## Supplemental Documentation for "Altair-LSFM A High-Resolution, Easy-to-Build Light-Sheet Microscope for Sub-Cellular Imaging"

### Altair Documentation

Dean Lab, UT Southwestern Medical Center

Nov 06, 2025

10 Future Releases 84

CHAPTER

ONE

### INTRODUCTION

#### 1.1 Why is Volumetric Imaging Important?

Biological processes occur in a fundamentally three-dimensional (3D) space, and the spatial organization of molecular structures is critical to cellular function. Traditional two-dimensional (2D) imaging provides only a limited cross-section of these dynamic events, often obscuring key interactions and leading to incomplete or misleading interpretations.

For instance, during mitosis, chromosomes undergo intricate rotations within the entire cellular volume, and microtubules dynamically polymerize and depolymerize as they engage and separate daughter chromosomes at their kinetochores. In 2D imaging, these structures inevitably drift in and out of the focal plane, making it challenging to track such dynamic processes over time. Similarly, in complex tissue environments, organelle positioning, cell-cell interactions, and subcellular signaling pathways are all inherently shaped by their 3D spatial context.

By capturing the full volume of a specimen, volumetric imaging enables a more accurate representation of cellular architecture and dynamic molecular events. This is particularly critical for understanding developmental biology, disease progression, and cellular interactions within their native environments. Emerging technologies such as light-sheet microscopy, expansion microscopy, and advanced computational imaging now allow researchers to visualize biological systems at unprecedented resolutions, bridging the gap between molecular detail and tissue-scale organization.

---

#### 1.2 The Importance of Parallelization

Historically, 3D imaging has been performed using spinning disk and laser scanning confocal microscopes. However, these microscopes waste significant excitation energy on out-of-focus regions, leading to unnecessary photodamage while requiring compensatory increases in laser power due to their low illumination duty cycle. This problem is further exacerbated by their sequential, point-by-point image acquisition, which limits speed and increases photobleaching.

**Illumination Confinement**

**Illumination confinement** refers to the restriction of excitation light to within the **depth of focus** of the detection objective, ensuring that only light contributing to image formation is used. This is a key advantage of **light-sheet fluorescence microscopy (LSFM)**, where the illumination plane is matched to the detection plane, minimizing out-of-focus excitation and reducing phototoxicity.

In contrast, depending on their design, light-sheet fluorescence microscopes (LSFM) restrict illumination primarily to the focal plane of interest, minimizing phototoxicity while maximizing imaging efficiency. Additionally, by leveraging highly sensitive scientific cameras (e.g., $4 \times 10^6$ pixels), LSFM enables massively parallelized image acquisition—recording entire planes in a single exposure rather than scanning point-by-point.

For instance, a typical laser scanning confocal microscope requires **~4.16s** to acquire a **2048 × 2048** voxel image, given a voxel dwell time of **1µs**. In contrast, an LSFM can image the same region in just **~5ms**, an **832-fold** increase in speed, all while maintaining a **5,000-fold longer** per-voxel dwell time. This extended dwell time enables improved signal accumulation, lower laser power requirements, and a significantly improved signal-to-noise ratio (SNR). As a result, LSFM is uniquely suited for long-term volumetric imaging at high spatiotemporal resolution while reducing photodamage.

**Spatial Duty Cycle**

The **spatial duty cycle of illumination** refers to the fraction of a sample that is actively contributing to image formation at any given moment during an imaging process. In **laser scanning confocal microscopy**, only a single point is illuminated at a time, resulting in a **low duty cycle** and requiring compensatory increases in laser power to maintain signal. In contrast, **light-sheet fluorescence microscopy (LSFM)** illuminates and images an entire plane at once, yielding a **duty cycle** that approaches 100%, enabling drastically reduced laser powers that reduce phototoxicity, .

#### 1.3 Challenges with 3D Imaging

To extract meaningful biological insight from volumetric imaging, microscopes must achieve sufficient **spatiotemporal resolution** while preserving molecular specificity and compatibility with advanced analytical techniques. This requires integration with biosensors, opto- and chemogenetic tools, and computer vision analyses that translate **high-dimensional 5D datasets** (x, y, z, $\lambda$, t) into quantitative biological readouts. Several key challenges must be addressed to ensure the accuracy and reliability of 3D imaging:

- **Nyquist Sampling in Space and Time:** To faithfully capture dynamic processes, the event of interest must be **Nyquist sampled** in both spatial and temporal dimensions. For example, the GTPase cycle times of Rho, Rac, and Cdc42 can be as short as 5s, necessitating volumetric acquisitions at $\leq$ 2.5s per volume with a spatial resolution of **<500nm**. Furthermore, resolution should ideally be **isotropic or nearly isotropic** to prevent morphology-dependent intensity artifacts—particularly for signaling events at the plasma membrane.
- **Multicolor Excitation and Detection:** To enable multiplexed cellular readouts, the microscope must support **simultaneous multicolor imaging** with achromatic performance across excitation and detection wavelengths. Microscope designs that rely on spatial light modulators (SLMs) for excitation often struggle to achieve this, as the diffraction efficiency of SLMs varies with wavelength.
- **Detection Sensitivity and Phototoxicity:** Imaging performance must be **photon-efficient** to minimize the need for high ectopic expression of signaling-active proteins and to mitigate **photobleaching and phototoxicity**—a challenge that is amplified when acquiring **10–100 slices per volume**. Maximizing quantum efficiency and minimizing out-of-focus excitation are crucial for maintaining live-cell viability.
- **Field of View Constraints:** When imaging dynamic processes in **extracellular matrix environments** (where cells can migrate in any direction) or within a **developing embryo**, the microscope must provide a sufficiently **large field of view (FOV)**—ideally **>100 × 100 × 100 µm**. Many leading **light-sheet fluorescence microscopes (LSFMs)** are optimized for small fields of view (e.g., 25µm), which limits their applicability to studies of large-scale tissue dynamics.
- **Avoidance of Computational Post-Processing Biases:** To ensure compatibility with **quantitative downstream analyses**, reliance on iterative **deconvolution** or structured illumination routines should be minimized. These techniques can introduce statistical artifacts that distort numerical analyses, particularly when measuring intensity distributions, localization precision, or dynamic molecular interactions.

**Nyquist Sampling**

**Nyquist sampling** is a fundamental principle in signal processing that dictates the minimum sampling frequency required to accurately reconstruct a signal without aliasing. According to the **Nyquist-Shannon sampling theorem**, a continuous signal must be sampled at least **twice the highest frequency component** present in the signal to ensure faithful reconstruction.

In microscopy, Nyquist sampling applies to both **spatial and temporal domains**:

- **Spatial Nyquist Sampling:** To accurately resolve features of size $d$, the sampling interval (pixel or voxel spacing) should be no greater than $d/2$. This ensures that high-frequency structural details are captured without loss of information.
- **Temporal Nyquist Sampling:** To track dynamic processes occurring at a characteristic frequency $f_c$, images must be acquired at a frequency of $\geq 2 \times f_c$ to prevent aliasing in time-lapse imaging.

Failure to meet the Nyquist criterion results in **aliasing**, where high-frequency components are misrepresented as lower-frequency artifacts, distorting biological measurements.

Meeting these criteria is essential for **accurate, high-throughput volumetric imaging** that captures cellular dynamics with sufficient fidelity to support rigorous biological interpretation.

---

#### 1.4 Why Build a Microscope?

The technology required to achieve **multiplexed volumetric imaging** with advanced probes and computer vision analyses already exists. However, the commercialization process imposes significant constraints on innovation. Microscope manufacturers prioritize **aesthetically attractive, highly engineered, and serviceable optical systems** that ensure a large return on investment. As a result, they tend to be **extremely conservative** in adopting emerging technologies.

Most commercially available microscopes take **over seven years** to develop, by which time they are often already obsolete due to rapid scientific advancements. A notable exception is the **Lattice Light-Sheet Microscope (LLSM)**, which was sublicensed by Zeiss to 3i shortly after its seminal publication. However, even in this case, it took another **six years** for Zeiss to release a consumer-friendly model—at a prohibitive cost of **~$1M USD**.

Beyond commercialization delays, **patent restrictions** further stifle innovation. The **highly fragmented and entangled intellectual property (IP) landscape** makes it difficult for new start-ups to enter the market. For example, despite their own **limited role in developing oblique plane microscopy (OPM)**, Leica has exclusively licensed a patent for **off-axis tertiary imaging systems**, effectively blocking broader commercialization of OPM.

As a result, reliance on commercial microscope development not only **delays technology adoption** but can actively **impede the dissemination of transformative imaging methods**. This reality makes in-house development of custom microscopy platforms essential for pushing the frontiers of biological imaging forward.

CHAPTER

### TWO

### OVERVIEW OF SUBCELLULAR LIGHT-SHEET MICROSCOPE TECHNOLOGIES

Light-sheet fluorescence microscopy (LSFM) has undergone significant advancements, leading to the development of specialized approaches tailored for **subcellular-resolution imaging**. These methods balance **resolution, field of view, optical sectioning, and phototoxicity** to accommodate different biological applications. Below, we provide an overview of existing high-resolution light-sheet microscope technologies, their strengths, and their limitations.

---

#### 2.1 Lattice Light-Sheet Microscopy (LLSM)

Lattice Light-Sheet Microscopy (LLSM) employs a **superposition of propagation-invariant beams** to generate a structured light-sheet that theoretically maintains a **narrow beam waist** over extended distances. However, in practice, these beams introduce **sidelobes** that increase with beam length, contributing to **out-of-focus blur** and degrading image contrast and resolution. As a result, LLSM is most effective for **small fields of view (~25µm)**, making it ideal for imaging **adherent cells and epithelial monolayers**.

A key limitation of LLSM is its reliance on a **spatial light modulator (SLM)** to sculpt its illumination beams. While this enables fine control over the excitation profile, it also introduces significant drawbacks: - **Limited Multicolor Excitation** – The SLM prevents simultaneous multicolor excitation, making LLSM suboptimal for **multiplexed biosensor imaging**. - **Reduced Light Throughput** – The optical train has an efficiency of only **~2.4%**, requiring higher-power and more expensive laser sources. - **Increased Complexity** – The integration of an SLM significantly increases the complexity and cost of the system, limiting accessibility.

---

#### 2.2 Field Synthesis

To address the limitations of Lattice Light-Sheet Microscopy (LLSM), the Fiolka lab developed **Field Synthesis**, a mathematical framework that reconstructs **time-averaged (dithered) lattice light-sheets** using **diffractive optics**. Unlike LLSM, which requires a **spatial light modulator (SLM)** to project complex amplitude and phase patterns onto the illumination objective, **Field Synthesis achieves the same illumination effect with greater light throughput and full compatibility with simultaneous multicolor excitation**.

Instead of directly sculpting the beam at the objective's pupil, Field Synthesis employs a **focused line scan over a binary pupil mask using a galvanometer during a single camera exposure**. This approach requires only:

- **A galvanometer for beam scanning**
- **A binary phase mask to shape the illumination**

- **A 4f telescope for beam relay**

Because of this design, **Field Synthesis provides resolution and photobleaching characteristics statistically indistinguishable from LLSM** while offering **2× higher acquisition speeds** in simultaneous multicolor imaging. For example, in a **sample-scanning configuration**, Field Synthesis was used to capture high-resolution, simultaneous imaging of **PI3K activity and dynamic filopodia and bleb formation** in **MV3 melanoma cells**. The increased imaging speed significantly reduced **motion blur**, which is particularly beneficial for visualizing rapid morphological transitions such as **filopodial buckling**.

---

#### 2.3 Dual Inverted Selective Plane Illumination Microscopy (diSPIM)

**Dual Inverted Selective Plane Illumination Microscopy (diSPIM)** is a commercially available light-sheet system that utilizes **two opposing objectives** to achieve isotropic resolution through sequential orthogonal illumination and detection. Unlike conventional LSFM, where a single objective delivers the light-sheet and another detects fluorescence, diSPIM alternates between two objectives, each serving both roles in successive acquisitions. This configuration enables **volumetric imaging with improved axial resolution**, as images from both orientations can be fused computationally to reconstruct a high-fidelity 3D volume. While diSPIM is **well-suited for live-cell imaging**, its reliance on sequential acquisition leads to **lower temporal resolution** compared to single-objective LSFM approaches. Additionally, alignment complexity and computational post-processing requirements can introduce challenges, particularly for rapid dynamic events. Despite these limitations, diSPIM remains a **flexible and widely adopted platform**, particularly in studies requiring high-resolution imaging of small organisms, embryos, and adherent cells.

#### 2.4 Axially Swept Light-Sheet Microscopy (ASLM)

Axially Swept Light-Sheet Microscopy (ASLM) overcomes many of the optical constraints of traditional light-sheet imaging by implementing a **rapidly scanned Gaussian beam** to create an extended, ultra-thin light-sheet. Unlike LLSM, ASLM minimizes **sidelobe artifacts**, resulting in superior **optical sectioning and contrast** across a larger field of view. Key advantages of ASLM include: - **Diffraction-Limited Isotropic Resolution** – Achieves **300–400 nm** axial resolution while maintaining high lateral resolution. - **Improved Optical Sectioning** – Reduces out-of-focus excitation, enhancing signal-to-noise ratios. - **Compatibility with Multicolor Imaging** – Unlike LLSM, ASLM does not require an SLM, allowing for **simultaneous multicolor excitation**, essential for biosensor applications.

However, ASLM is inherently **slower than conventional light-sheet methods**, as the sweeping mechanism imposes limits on acquisition speed. Additionally, its implementation requires precise **synchronization between beam scanning and camera acquisition**, adding some complexity to its control systems.

---

#### 2.5 Oblique Plane Microscopy (OPM)

The orthogonal illumination and detection geometry used in most **light-sheet fluorescence microscopes (LSFMs)** comes with several **underappreciated limitations**. This configuration is **incompatible** with many standard laboratory imaging setups, including **hardware-based autofocus systems** that mitigate thermal and mechanical drift, **standard imaging dishes**, **multi-well plates**, and **microfluidic devices** designed for establishing **chemotactic gradients** or delivering **controlled shear stress** in parallel plate flow chambers.

Additionally, LSFMs often require **large imaging chambers** (e.g., **~8.5 mL** for **LLSM**), leading to **high reagent consumption**, which can be cost-prohibitive for experiments involving **chemogenetic or pharmacological perturbations**. The use of **high numerical aperture (NA) water-dipping objectives** further **compromises sterility**, making long-term imaging of slow biological processes, such as **sarcomerogenesis over ~24 hours**, particularly challenging.

Although ultra-thin **fluorocarbon foil-based cuvettes** have been explored as a solution, even slight **refractive index mismatches** introduce **spherical aberrations**, degrading image resolution and sensitivity. These factors highlight the need for alternative light-sheet implementations that maintain **high optical performance while accommodating diverse experimental conditions**.

Oblique Plane Microscopy (OPM) represents a **single-objective light-sheet imaging approach** that overcomes these challenges. Here, owing to its unique non-coaxial design, an obliquely launched illumination beam is used to achieve volumetric imaging without requiring an orthogonally positioned objective. This method has gained popularity for **live-cell imaging** due to its advantages: - **Simplified Geometry** – Requires only a single high-NA objective, making it more compact and easier to integrate into existing microscope setups. - **High-Speed Volumetric Imaging** – Can acquire full 3D volumes at video rate. - **Compatible with Conventional Sample Mounting** – Unlike traditional LSFM, OPM does not require complex sample positioning or embedding techniques.

Despite its advantages, OPM suffers from **anisotropic resolution** and often requires **computational post-processing** (e.g., shearing correction) to reconstruct datasets accurately. Additionally, due to the oblique illumination geometry, axial resolution may degrade deeper into the sample.

| Microscope Type | Strengths | Limitations |
|---|---|---|
| **LLSM** | • High-resolution for small samples<br>• Excellent optical sectioning | • Limited field of view (~25µm)<br>• Incompatible with multicolor excitation<br>• Low light throughput (~2.4%)<br>• Requires high-power lasers |
| **Field Synthesis** | • Statistically indistinguishable from LLSM<br>• Higher light throughput<br>• Compatible with simultaneous multicolor imaging<br>• Faster acquisition (~2× speed vs. LLSM) | • Requires scanning with a galvanometer<br>• Less widely implemented than LLSM<br>• Slightly lower spatial control than LLSM |
| **diSPIM** | • Dual-objective design improves axial resolution<br>• Enables isotropic volumetric imaging via fusion<br>• Well-suited for live-cell and embryo imaging<br>• Commercially available and widely adopted | • Multiple volumes must be acquired<br>• Requires post-processing for image fusion<br>• Greater alignment complexity |
| **ASLM** | • High contrast and improved optical sectioning<br>• Isotropic resolution (~300–400 nm)<br>• Compatible with multicolor imaging | • Lower acquisition speed due to beam sweeping<br>• Requires precise synchronization<br>• Complex scanning mechanics |
| **OPM** | • Single-objective system (simplified geometry)<br>• High-speed volumetric imaging<br>• Compatible with conventional sample mounting | • Anisotropic resolution<br>• Requires computational post-processing<br>• Axial resolution degrades with depth |

CHAPTER

THREE

### SOFTWARE

The **software for optical simulations and computer-aided design (CAD)** is optional and only required for users who wish to **modify or customize Altair**. However, **navigate is mandatory** for microscope operation unless an alternative **open-source acquisition platform** is used. For **Altair-LSFM**, which operates in a **sample-scanning geometry**, **shearing** is necessary to properly align data spatially; while various software options exist, we provide **Python-based scripts** for this task. Lastly, **deconvolution is optional**, depending on the user's specific imaging and analysis needs.

---

#### 3.1 Optical Simulations

Performing accurate optical simulations is essential for designing and validating the illumination pathway of **Altair**. To facilitate this process, we used **Zemax OpticStudio (Ansys)** to model the full illumination system, optimizing the placement of each optical element to achieve the desired focusing and collimation properties.

To replicate or modify these simulations, a **Zemax OpticStudio (Ansys)** professional or academic license is needed. This enables optical modeling, ray tracing, and tolerance analysis. The **Zemax simulation files** used to design the Altair illumination system are included in the project's GitHub repository. These files allow users to inspect and modify the optical layout, test different lens configurations, and refine performance metrics as needed.

---

#### 3.2 Computer-Aided Design

For the design of custom parts and the baseplate, we used **Autodesk Inventor**. Academic licenses for Autodesk products are available free of charge on their website.

To ensure **Autodesk Inventor** correctly locates and manages the CAD files, it is necessary to set up a **new project** and define the workspace. We recommend following these steps:

##### 3.2.1 Setting Up a New Project in Autodesk Inventor

1. **Launch Autodesk Inventor**
2. Navigate to **File** → **Manage** → **Projects**

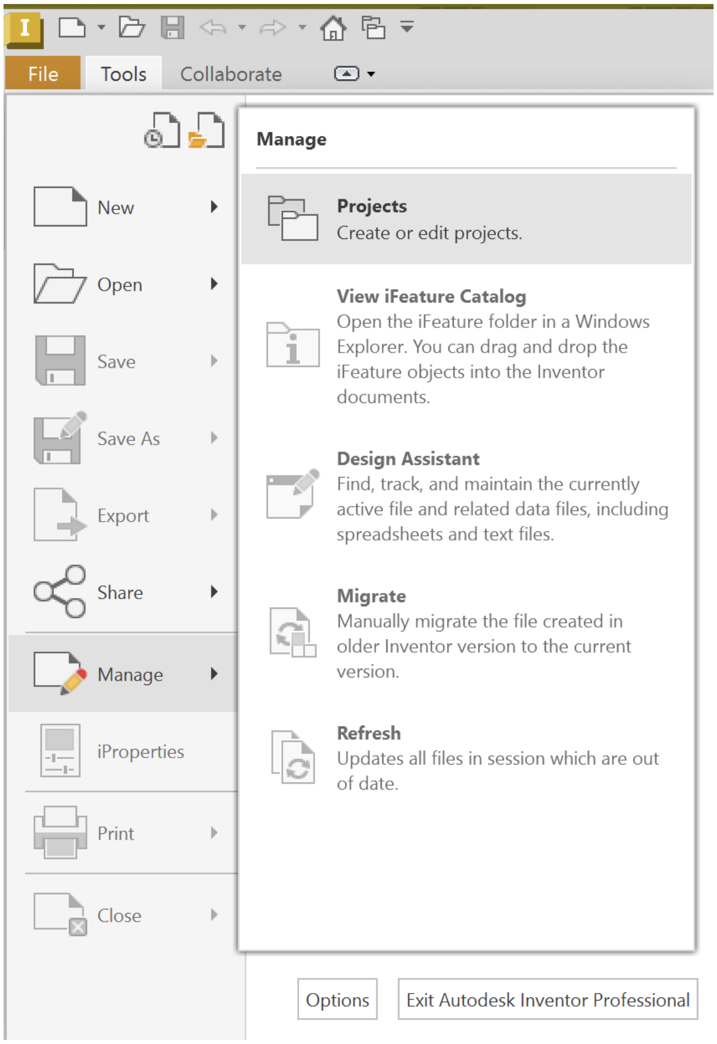

3. When the **Projects** window appears, select **New**

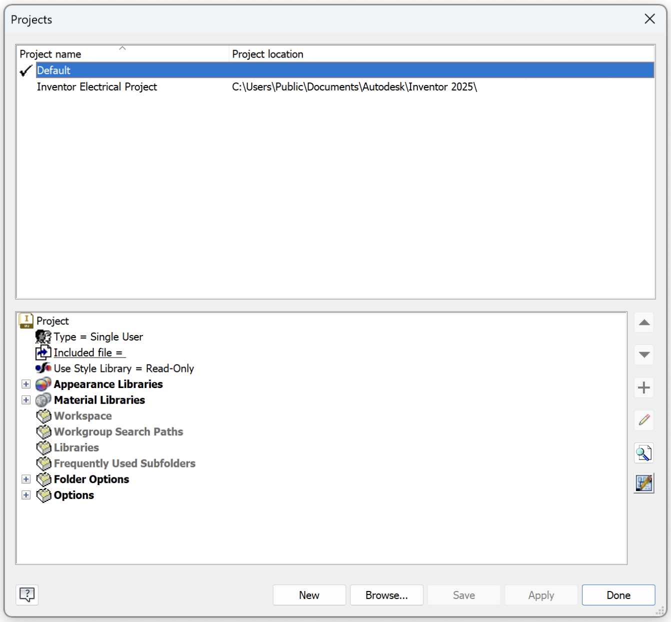

4. Choose **New Single User Project**

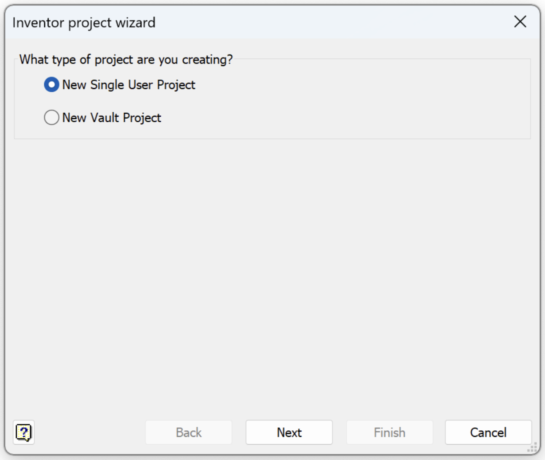

5. **Specify a Project Name** and set the **root directory of the cloned Altair GitHub repository** as the **Project (Workspace) Folder**

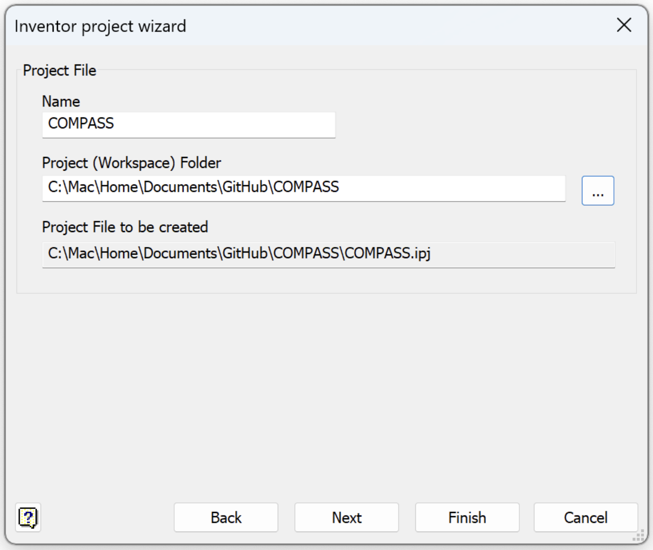

By configuring Autodesk Inventor in this way, all CAD files associated with **Altair** will be properly linked, ensuring seamless loading of all components and assemblies.

---

#### 3.3 Microscope Control Software: navigate

To control and operate **Altair**, we use **navigate**, a Python-based software package designed to provide flexible and intelligent microscope control. **navigate** enables users to interact with the system through a variety of advanced modalities, including **smart acquisition routines** that dynamically adjust microscope performance based on the biological specimen.

##### 3.3.1 Features of navigate

**navigate** provides key functionalities that enhance the efficiency and adaptability of the Altair system:

- **Automated Smart Acquisition** – Dynamically optimizes imaging parameters in real time, adapting to specimen properties to improve image quality.
- **Hardware Abstraction Layer** – Enables seamless control of multiple hardware components, allowing users to configure, modify, and expand the microscope's functionality.
- **High-Throughput Data Acquisition** – Facilitates volumetric imaging with high-speed acquisition routines, ensuring efficient data collection without compromising spatial or temporal resolution.

##### 3.3.2 Installation and Access

**navigate** is **open-source** and publicly available on GitHub. Installation instructions, along with comprehensive documentation, can be found here: navigate Installation Guide

---

#### 3.4 Post-Processing

After image acquisition, post-processing is often necessary to extract meaningful biological information. The **navigate** software package includes libraries for performing **basic image post-processing tasks**, such as **shearing correction**, which can be applied directly to acquired datasets. We provide Python-based scripts in the *post_processing* folder of this repository as an example.

##### 3.4.1 Deconvolution

For more advanced processing, such as **deconvolution**, we recommend using **PetaKit5D**. This software package is optimized for handling large-scale volumetric datasets and provides state-of-the-art deconvolution algorithms to enhance image quality and resolution[1]. **PetaKit5D** must be installed separately and can be found here: PetaKit5D GitHub Repository

###### Citations

[1] Ruan X, Mueller M, Liu G, Görlitz F, Fu TM, Milkie DE, Lillvis JL, Kuhn A, Gan Chong J, Hong JL, Herr CYA, Hercule W, Nienhaus M, Killilea AN, Betzig E, Upadhyayula S. *Image processing tools for petabyte-scale light sheet microscopy data.* **Nat Methods.** 2024 Dec;21(12):2342-2352. doi: 10.1038/s41592-024-02475-4. PMID: 39420143, PMCID: PMC11621031.

CHAPTER

FOUR

### ROOM AND HARDWARE CONSIDERATIONS

Installing a **Altair** system requires careful planning to ensure optimal performance and long-term stability. Proper preparation includes considerations for **room setup, optical table selection, acquisition computer specifications, laser identification, and environmental controls**. Each of these factors plays a significant role in instrument performance. Consequently, before beginning the installation process, we recommend evaluating the information, some of which is provided simply as educational resources.

#### 4.1 Preparation of the Imaging Room

A common question that we get is how to set up the room for imaging. This section will go over the steps that we took to set up the room for imaging. The stated directions assume that the system will be housed in the United States. If you are in a different country, you may need to adjust the electrical configuration accordingly.

---

##### 4.1.1 Power Outlets

The first thing to consider is the power outlets. For most imaging systems, you will need standard 120V outlets. If you plan on having a Ti-Sapphire laser, then 240V outlets are likely necessary. We typically like to have power strips on the walls, so that you have plenty to choose from.

---

##### 4.1.2 Temperature Control

Temperature control is important for imaging. We recommend having a temperature sensor in the room for controlling the temperature in that zone. Otherwise, the electronics and people can heat up the room significantly, which can cause drift in the imaging system. HVAC systems can also be a source of dust. Ideally, the ventilation to the room will be HEPA filtered and the air flow will be directed away from the microscope.

---

##### 4.1.3 Painting the Walls

We typically paint the walls matte black. The matte black walls are helpful for reducing the glare from the monitor and other sources of scattered light from being detected with the camera.

---

##### 4.1.4 Network Connections

We recommend having a 10G network or faster for transferring data. This will help with the transfer of large image files that are generated during imaging.

---

##### 4.1.5 Curtains

If you expect other people to work in the space at the same time, or if you want multiple microscopes, we recommend having blackout curtains to separate the spaces. If the imaging space does not have a fire sprinkler, then the curtain will likely need a gap between the ceiling and the curtain to meet building regulations.

---

##### 4.1.6 House Air

You will need house air operating at ~100 PSI to float the table. Having gas cylinders is not an option, as the tank will likely have to be replaced too frequently.

---

##### 4.1.7 Floor Flatness

The floor should be flat. If the floor is not flat, then the table will need special accessories to make it level.

---

##### 4.1.8 External Sources of Vibration

Make sure that the room is not near any external sources of vibration. This can include elevators, HVAC units, or other heavy machinery. Especially when imaging in soft substrates such as expanded tissues, or reconstituted extracellular matrix environments, vibrations can degrade imaging quality. In general, microscopy rooms in basements are preferable, as they are less likely to be affected by external sources of vibration.

---

##### 4.1.9 Water Damage

Also consider worst-case scenarios such as a water leak from the room above. If there is a significant risk that the fire sprinkler system could go off, then you may want to consider mechanisms of protecting the electronics and optics from water damage.

#### 4.2 Optical Table

The optical table is the most important part of the imaging room. We recommend a large table, as it is better to have too much table than not enough. Ideally, the room is large enough that the table is not completely against the wall, so that you have access to the entire table.

---

##### 4.2.1 Table Height

Having a table at the right height makes working at the table more comfortable. We recommend that the table surface be located approximately 38 - 40 inches from the floor.

---

##### 4.2.2 Table Thickness

We typically purchase tables with an 18” thickness. In theory, thicker tables better dampen vibrations arising from on the table itself, including fans from cameras, voice coils, stages, etc. Practically, in our experience, the table thickness helps mitigate vibrations arising from the building itself as well.

---

##### 4.2.3 Table Accessories

We recommend getting a table with a shelf and power strips. The shelf is useful for storing electronics and other equipment in a location where their vibrations will not be coupled to the table.

---

##### 4.2.4 Hole Pattern

All of our optical mounts are designed to fit the 1” grid pattern with 1/4”-20 holes. If you would like to have metric holes, you will need to customize the mounts to fit the table.

---

##### 4.2.5 Casters

Casters serve as a very convenient mechanism for moving the table around. We thus highly recommend getting a table with casters. Otherwise, most tables are sufficiently heavy that they will not be able to be moved once installed.

##### 4.2.6 Vibration Damping

Many vibration damping systems actually amplify vibrations at low frequencies, which are common building-associated vibrations. To mitigate this, we recommend using a tuned damping system. TMC provides and UltraDamp series of isolation systems that

##### 4.2.7 Table Installation

We recommend following the manufacturer's instructions for installing the table. Importantly the frame, and the table top should be level in order to achieve the best performance. Given the weight of the table, it is likely that professional installation will be required.

##### 4.2.8 Recommended Table

We typically purchase Performance Series CleanTop Research Grade 784 Series Optical Tables from TMC. These tables are available in a variety of sizes, and the weight can be estimated from the lateral dimensions of the table and its thickness. To increase the weight and damping capacity of the table, we opt for the 18" standard thickness. This is placed on a set of 14-Special UltraDamp legs that are equipped with casters and provide a 20" working height, and a total overall table height of 38".

#### 4.3 Computer Specifications

Modern scientific cameras are capable of capturing images at very high frame rates. For example, the Orca-Lightning from Hamamatsu, when imaging a region of interest of 4608x128, can image at 2203 frames per second, totalling 1.3 terapixel per second. It is important to have a computer that can handle this data rate. For many applications, we recommend Colfax International's SXP9000 workstation, which includes several convenient features, including:

- A large number of high-speed peripheral slots. While originally designed to host GPU cards, these slots can be used for other high-speed peripherals, such as frame grabbers.
- A high-speed NVMe SSD, which can achieve upwards of 20 GB/s under ideal conditions.
- 10GbE LAN - 10G Ethernet or faster, if supported by your institution, is highly recommended for transferring data to and from the computer. This computer provides this high-speed connection without necessitating an additional network card, freeing up precious space in the computer.
- USB-C ports that can be used for image display, keyboard, and mouse. Again, by eliminating the need for a graphics card, available space is available to other peripheral devices.
- Redundant power supplies - in the event of a power supply failure, the computer will continue to operate. Moreover, they will be able to drive some of the more power-hungry components, such as frame grabbers.
- Aggressive cooling - the computer is designed to handle the heat generated by the high-speed components.

##### 4.3.1 Standard Build

A general build for a imaging computer is as follows:

- Windows 10 Pro. This is the most common operating system for scientific imaging applications.
- 2x Intel Xeon Silver CPUs Total of 16 Cores/32 Threads @ 3.2GHz. Higher speeds, and a greater number of cores is advantageous but expensive.
- 128 GB of 3200 MHz DDR4 RAM. This is more than sufficient for most applications.
- 800 GB M.2 NVMe SSD. Used to host and run the operating system.
- 20 TB 7200 RPM SATA HDD. Used as a 'cold storage' for long-term data storage.
- NVIDIA T1000 Video Card. This is a low-end video card that can be used to drive displays and perform some basic image processing. The priority for this computer is driving the microscope and acquiring data, so a high-end video card is not necessary.
- Intel X710-T2L 10GbE Card. If the motherboard does not provide 10GbE, this card can be used to provide high-speed network connectivity.
- 7.68 TB NVMe SSD. Used as the primary data drive.

**Note:** Recently, Hamamatsu released a Linux driver for their cameras. Linux provides several advantages to Windows-based operating systems, including read and write operations with lower overhead. This is especially important for next-generation file formats, such as N5, OME-Zarr, and Zarr, all of which break large images up into smaller 'chunks' that can be read and written independently. To learn more, we recommend reading Moore et al. 2021, and Moore et al. 2023.

#### 4.4 Laser Sources

When designing a microscope for imaging, the choice of light source is critical. In general, an ideal imaging system will provide fast spectral switching, high light throughput, provide stable power emission, have a long lifetime, be cost-effective, and be low complexity.

Several light have been developed for use in microscopy, including arc lamps, light emitting diodes (LEDs), solid state lasers (including fiber lasers, optically pumped semiconductor lasers, diode pumped solid state lasers), supercontinuum lasers, and ultrafast lasers (including ti-sapphire lasers). A few key parameters to consider are discussed below.

##### 4.4.1 Linewidth

One key aspect of any light source is the linewidth of its emission. This is the spectral width of a light source in nanometers of the spectral density of the emitted electric field. It can also be reported in wavenumbers ($cm^{-1}$) or frequency (Hz). Narrow linewidths enable more aggressive detection bands, reduced bleedthrough, and a reduced background. LEDs typically have a linewidth of 20-50 nm, which is much broader than the 0.1-1 nm linewidth of a laser. As such, we prefer lasers for our imaging systems.

##### 4.4.2 Beam Divergence

The $M^2$ value of a beam is the degree at which it can be focused to a given beam divergence angle (e.g., numerical aperture). If you have a higher $M^2$ value of ~1.6, it will focused to a laser spot that is 1.6x larger than the diffraction limit. As such, for high-resolution imaging, it is important to have a low $M^2$ value. We prefer lasers with an $M^2$ value of <1.2 for our imaging systems. Fiber lasers, such as those offered by MPB Communications, provide very low $M^2$ values. However, these systems require external mechanisms for controlling the power output, which is a major disadvantage. For lasers with high $M^2$ values, additional optics may be necessary to clean up the beam profile, such as a spatial filter.

---

##### 4.4.3 Modulation

The ability to modulate the power of a light source is important for many imaging applications. It is not uncommon to use an acousto-optic modulator (AOM), acousto-optic tunable filter (AOTF), or electro-optic modulator (EOM) to modulate the power of a laser. AOMs induce sound waves within a crystal to change the refractive index of the crystal, which alters the angle of the light at a single wavelength. Likewise, AOTFs use a variable frequency sound wave to operate at multiple wavelengths. EOMs, such as a Pockels cell, induce a directionally dependent change in refractive index in a crystal, which can change the polarization of the light passing through it. These are especially common in ultrafast laser systems. Both AOMs and EOMs can be used to modulate the power of the laser, with rise times of ~5-50 ns and Acousto-optic devices offer rise times 0.25 - 1 ns, respectively. However, especially for continuous wave (CW) lasers, these devices serve as a source of additional complexity, cost, and potential failure points. For example, we have noted that AOMs can be sensitive to temperature changes, which can lead to drift in the power output. As such, we prefer lasers with built-in modulation capabilities.

---

##### 4.4.4 Vibrations

Many lasers require active cooling, and are often placed directly on the optic table, which can lead to vibrations. These vibrations can be detrimental to imaging quality, especially when imaging in soft contexts such as expanded tissues. Nonetheless, if insufficient cooling is provided, fluctuations in laser emission or instability can occur.

---

##### 4.4.5 Power

The power of the laser that is needed depends critically on the light throughout of the illumination train, as well as the size of the field of view. While some light sources such as supercontinuum lasers can provide high power (e.g., >3W), their emission is spread across a large spectral range, reducing their spectral power density at any given wavelength to a few mW/nm. We typically recommend lasers with a power of 100 mW or greater for imaging.

---

##### Recommended Light Sources

For all of these reasons, we recommend the use of optically pumped semiconductor lasers (OPSLs) for imaging. OPSLs are solid state lasers that use a diode to optically pump a semiconductor gain medium which can be tuned to adjust the emission wavelength of the laser. Importantly, OPSLs have a narrow line width (<10 pM), low $M^2$ value, can modulate the laser on microsecond timescales, and achieve high and stable emission powers.

OPSLs are available from several manufacturers and can typically be driven in either a constant power or constant current mode. We typically use the former, as the laser uses active feedback to stabilize the emission power. To eliminate challenges associated with vibrations, we use single mode fiber-coupled OPSLs, which can be removed from the optical table provide very nice beam quality.

For our imaging systems, we have used the following lasers:

- Coherent Galaxy laser combiner and OBIS lasers
- Oxxius LaserBoxx with LuxX lasers.
- Omicron LightHub Ultra with LuxX and OBIS lasers.

#### 4.5 Remote Focus Actuator

In Axially Swept Light-Sheet Microscopy (ASLM), axial scanning is achieved by driving a mirror placed at the focus of a remote objective. To maintain high-fidelity imaging, the remote focus actuator must precisely synchronize its motion with the rolling shutter of the camera. However, designing for speed and accuracy involves trade-offs. As the stroke of the actuator increases, its mechanical bandwidth decreases, limiting the maximum acquisition rate for large field of view imaging. Similarly, higher scan frequencies tend to reduce the scan amplitude and introduce phase delays, which can degrade image quality, especially at the edges of the field of view. Therefore, it is crucial to select an actuator that balances these factors effectively.

Early implementations of ASLM used piezoelectric actuators, but modern systems often employ voice coil actuators for improved performance. We have tested two commercially available systems: the LFA2004 from Equipment Solutions and the Blink from Thorlabs. The LFA2004 is reliable, easy to configure, and sufficient for most imaging applications, while the Blink offers higher performance at the cost of greater complexity and sensitivity to tuning. For most users, we recommend the LFA2004.

When ordering the LFA2004, we typically request the stage configured with the SCA814 amplifier and SPS15 power supply. For standard ASLM configurations, we recommend tuning the system for a ~2 gram mirror with a 30µm sawtooth waveform (90% duty cycle). The factory can preconfigure the unit to accept an analog position command by default (servo active). If oscillation or high-frequency noise is observed when the servo is engaged, you can fine-tune the control loop via the RS232 interface by adjusting the “A” and “B” parameters. A common stable configuration is A 40 20 and B 80 5, though slight adjustments may be necessary depending on your optical load and mechanical setup.

### CHAPTER FIVE

### DESIGN PROCESS

#### 5.1 Initial Lens Selection

Prior to starting optical simulations in Zemax, it's convenient to start with straightforward calculations to determine which lenses to use in the optical train to achieve the desired field of view (FoV) for your detection path. In our case, our detection path consisted of a 400 mm tube lens and a Nikon 25x/1.1 numerical aperture (NA) immersion detection objective.

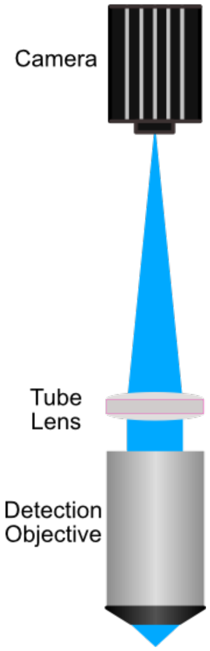

Fig. 1: **Figure 1:** A simple schematic for a widefield detection path. Here, a detection objective captures light, which is then focused onto the active sensor of a CMOS camera using a tube lens.

**Calculating the Focal Length of an Objective**

The focal length ($f$) of a microscope objective can be determined using the **nominal focal length** of the manufacturer's standard tube lens and the **objective's magnification**:

$$f = \frac{f_{\text{tube lens}}}{M}$$

where:

- $f_{\text{tube lens}}$ is the standard focal length of the tube lens.
- $M$ is the nominal magnification of the objective.

Standard tube lens focal lengths vary by manufacturer:

- **Nikon / Zeiss**: 200 mm
- **Olympus**: 180 mm

**Example Calculation** For a **Nikon 25× NA 1.1 objective**, the focal length is calculated as:

$$f = \frac{200 \text{ mm}}{25} = 8 \text{ mm}$$

To determine the target FoV, start with determining the final magnification (M) of the system using the ratio of the focal lengths of the tube lens ($f_{\text{TL}}$) and the detection objective ($f_{\text{DO}}$, Equation 1.).

$$M = \frac{f_{\text{TL}}}{f_{\text{DO}}}$$

From there, determine the resulting FoV of the detection path by dividing the total camera sensor size (in mm) by the magnification, and then converting into microns (Equation 2).

$$\text{FoV } (\mu m) = \left(\frac{W_{\text{Sensor}}}{M}\right) \times 1000$$

For our system, where $f_{\text{TL}}$ = 400 mm, $f_{\text{DO}}$ = 8 mm, and W $_{\text{Sensor}}$ = 13 mm, this resulted in a FoV of ~266 $\mu m$, meaning that we want to select lenses in our illumination path to produce a light sheet as close to 266 $\mu m$ in length as we can achieve.

The overarching goal of a standard optical system is to both mold light into a particular shape and direct it to a particular location. In our case, our optical system works to convert an input gaussian beam into a thin light sheet that illuminates our sample. There are a few sets of criteria that help guide our potential lens selection:

- As mentioned, we want our final light sheet size to ideally cover the full FoV of our detection path (~266 $\mu$m)
- At the focus of our cylindrical lens, we want the beam spot size to stay under the size of our resonant galvo (12 mm diameter)
- We need the focal distance between the cylindrical lens and the galvo mirror system to be greater than ~55 mm due to mechanical considerations of the mirror mount used

With these criteria in mind, we can calculate a theoretical estimate of what our beam size is after each of our lenses. We do this by considering every pair of lenses (i.e. Lens 1 & 2, Lens 2 & 3, $\ldots$) as a sort of 4F magnification system, where the resulting image size of the pairs is determined by the ratio of their focal lengths ($f_{\text{n}}$) as follows:

Essentially, we can cascade these calculations through our lenses and make sure that our choices in their focal lengths produce our desired beam characteristics as light propagates through the system. In our case, our chosen system featured 4 lenses from Thorlabs: L1 = 30 mm, L2 = 80 mm, L3 (Cylindrical) = 75 mm, and L4 = 250 mm.

We can then take these lens choices and load them into Zemax OpticStudio to verify the characteristics of our system.

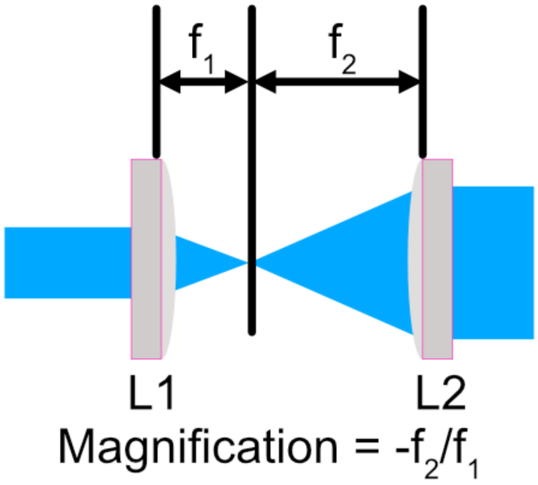

Fig. 2: **Figure 2:** A diagram of a 4f optical system. Here, the ratio of the focal lengths of the lenses determines the magnification of the system.

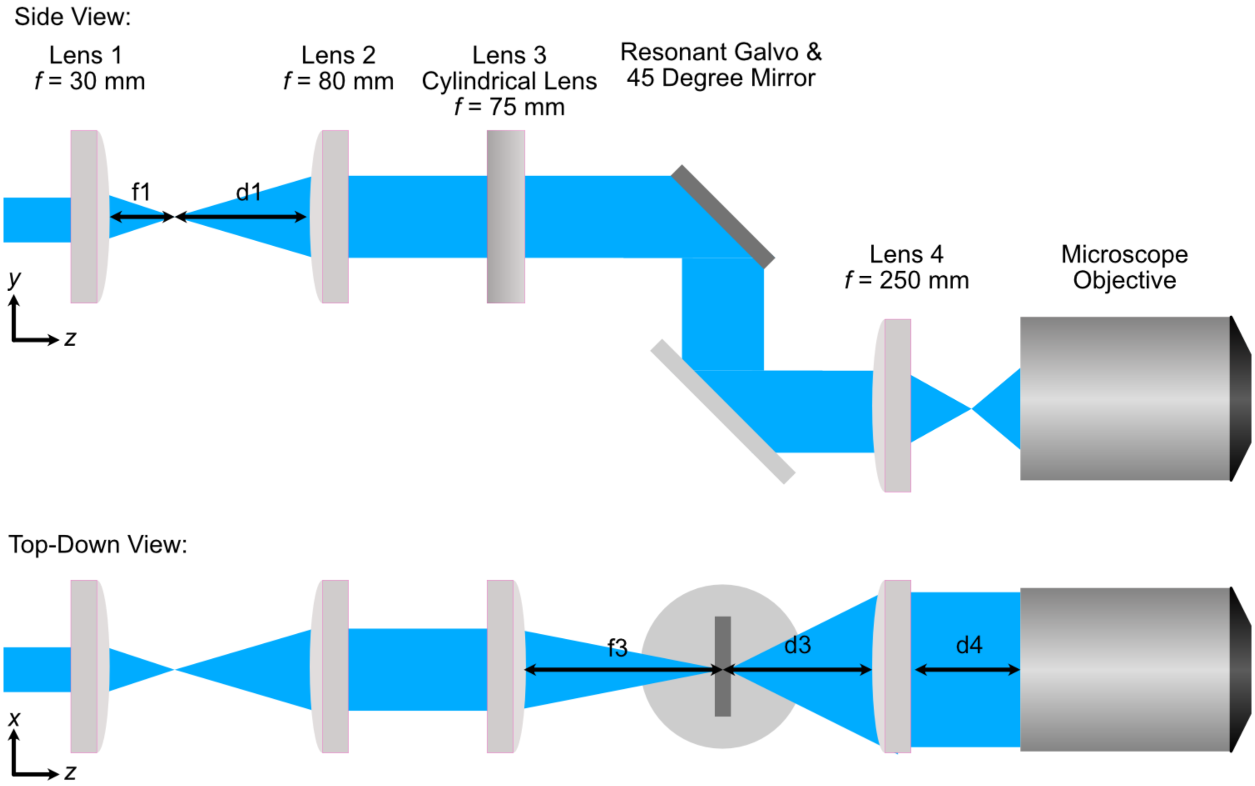

Fig. 3: **Figure 3:** A schematic of the optical path for a light-sheet microscope. The system consists of four lenses, a cylindrical lens, a resonant galvo, a 45-degree mirror, and an illumination objective. Top and bottom views are shown.

#### 5.2 Zemax Simulation Setup Process

With our chosen lenses in mind, we can download Zemax files associated with each lens directly from Thorlabs website and set up our simulation.

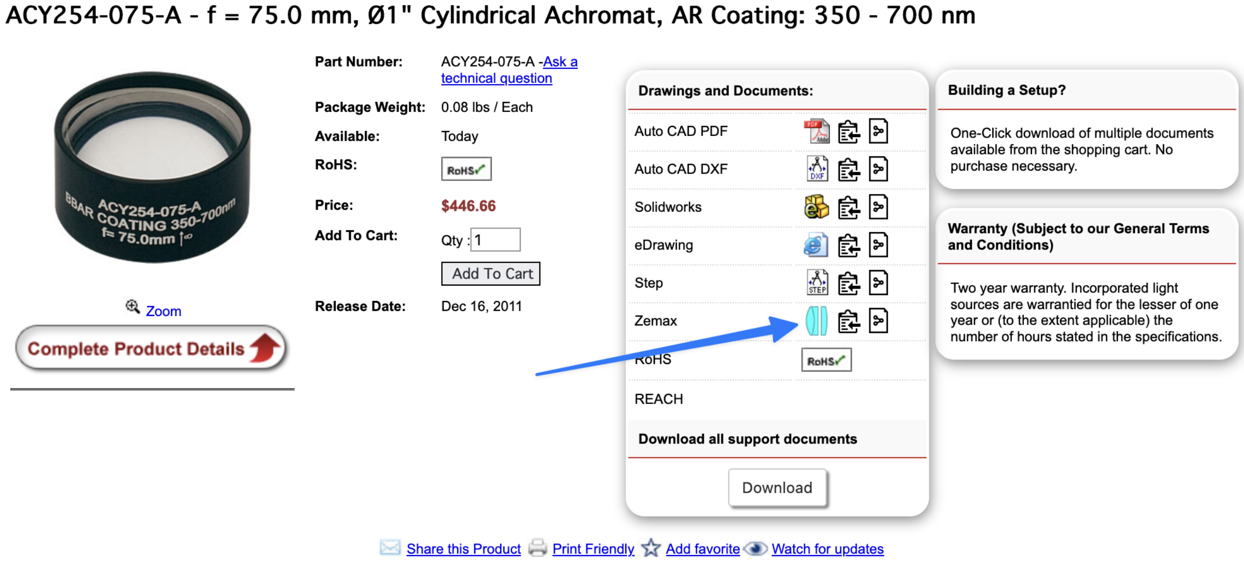

Fig. 4: **Figure 4:** An example of a Zemax file download from Thorlabs. The file contains the optical properties of the lens, which can be imported into Zemax OpticStudio.

Here, we use Zemax as a tool to find the optimal placement of all the lenses of our system based on whether or not the input beam should be focusing or collimated after each lens. As a general rule of thumb, one should build optical systems in Zemax in an element-by-element manner instead of adding all the optical elements and trying to then optimize aspects of it. Our general flow involves adding a lens to the system and then optimizing for either either a focused or collimated beam, and then adding in the next lens and doing the same process until all lenses are placed in the system. This is described in more detail below.

For our particular system, our generalized process went as follows:

1. Create a new file that will be used as our lens assembly file
2. Set aperture size in Zemax to match our original laser spot size (2.4 mm).
3. Open the Zemax file associated with Lens 1, then copy and paste the surfaces into our assembly file.
4. Use the optimization wizard to set a focusing optimization with the distance after L1 (f1) as the variable to find the correct position of L1's focus.

**Figure 5:** The optimization wizard in Zemax. Here, the user can set the optimization type and the variable to optimize. In this case, the user is optimizing for spot size.

5. Run the optimization, then remove the variable for f1.
6. Open the Zemax file associated with Lens 2, then copy and paste the surfaces into our assembly after Lens 1
7. Use the Optimization Wizard to set an angular (collimation) optimization, with the distance between L1's focus and L2 (d1) as the variable.

**Figure 6:** The optimization wizard in Zemax. Here, the user can set the optimization type and the variable to optimize. In this case, the user is optimizing for collimation.

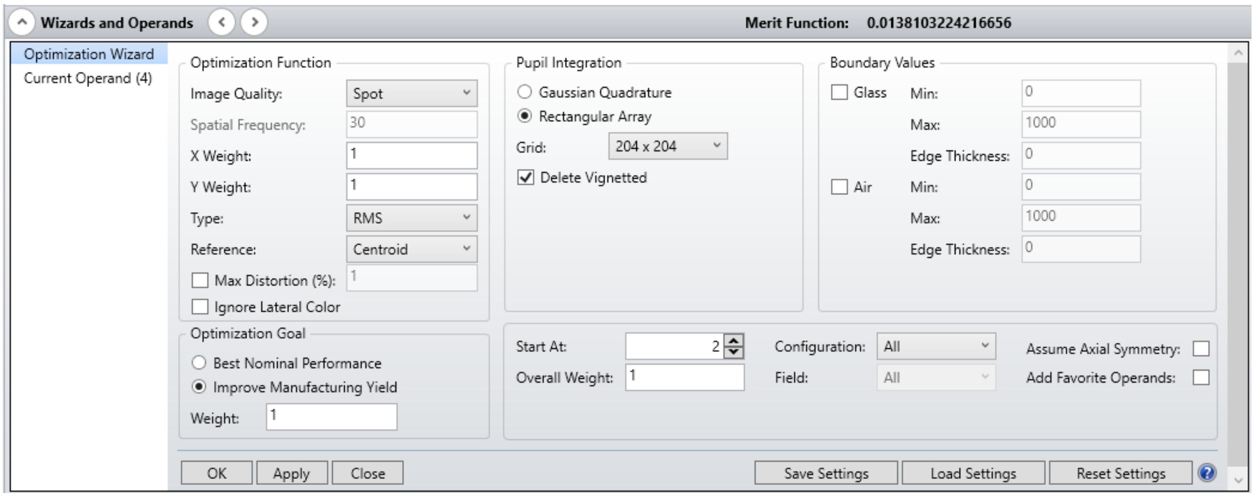
Wizards and Operands
Merit Function: 0.0138103224216656
Optimization Wizard
Current Operand (4)
Optimization Function
Image Quality: Spot
Spatial Frequency: 30
X Weight: 1
Y Weight: 1
Type: RMS
Reference: Centroid
Max Distortion (%): 1
Ignore Lateral Color
Optimization Goal
Best Nominal Performance
Improve Manufacturing Yield
Weight: 1
Pupil Integration
Gaussian Quadrature
Rectangular Array
Grid: 204 x 204
Delete Vignetted
Boundary Values
Glass Min: 0
Max: 1000
Edge Thickness: 0
Air Min: 0
Max: 1000
Edge Thickness: 0
Start At: 2
Overall Weight: 1
Configuration: All
Field: All
Assume Axial Symmetry:
Add Favorite Operands:
OK
Apply
Close
Save Settings
Load Settings
Reset Settings

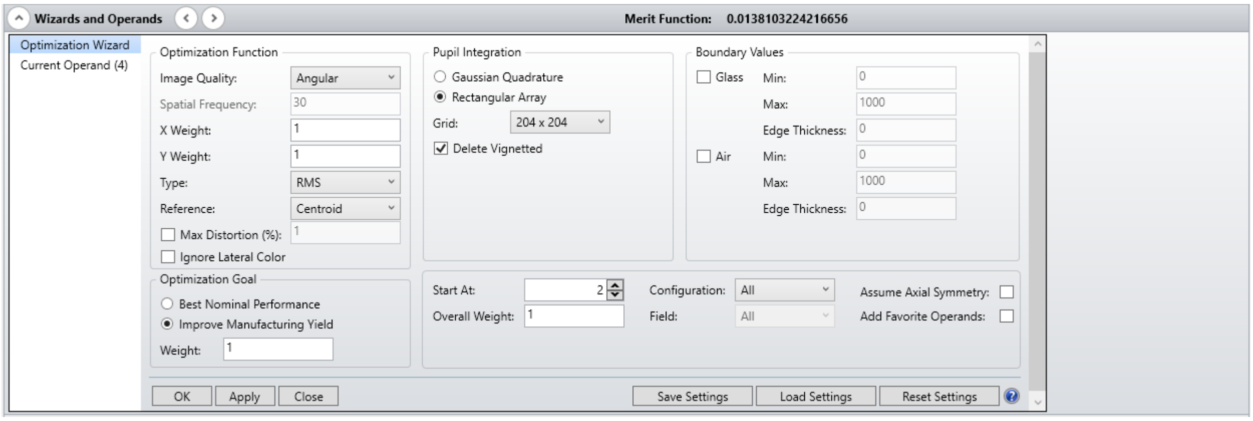
Wizards and Operands
Merit Function: 0.0138103224216656
Optimization Wizard
Current Operand (4)
Optimization Function
Image Quality: Angular
Spatial Frequency: 30
X Weight: 1
Y Weight: 1
Type: RMS
Reference: Centroid
Max Distortion (%): 1
Ignore Lateral Color
Optimization Goal
Best Nominal Performance
Improve Manufacturing Yield
Weight: 1
Pupil Integration
Gaussian Quadrature
Rectangular Array
Grid: 204 x 204
Delete Vignetted
Boundary Values
Glass Min: 0
Max: 1000
Edge Thickness: 0
Air Min: 0
Max: 1000
Edge Thickness: 0
Start At: 2
Overall Weight: 1
Configuration: All
Field: All
Assume Axial Symmetry:
Add Favorite Operands:
OK
Apply
Close
Save Settings
Load Settings
Reset Settings

8. Optimize, then remove the variable for d1.
9. Open the Zemax file associated with Lens 3, then copy and paste the surfaces into our assembly after Lens 2.
10. Use the optimization wizard to set an $X$-focusing optimization with the distance after L3 (f3) as the variable.
11. Optimize, then remove the variable for f3.
12. Place in resonant galvo and 45 degree mirror surfaces at the location of f3.
13. Open the Zemax file associated with Lens 4, then copy and paste the surfaces into our assembly after the 45 degree mirror.
14. Use the optimization wizard to set an $X$-collimation optimization with the distance between the 45 degree mirror and L4 (d3) as the variable.
15. Optimize, then remove the variable for d3.
16. Open the Zemax file associated with our Illumination Objective, then copy and paste the surfaces into our assembly after L4.
17. Use the Optimization Wizard to set an $X$-focusing Optimization with the distance between L4 and the objective (d4) as the variable.
18. Optimize

---

#### 5.3 Zemax Simulation Analysis

Within Zemax, there are numerous analysis tools available to investigate different characteristics of optical systems. Our analysis will primarily be guided by the Geometric Image Analysis, Huygen's PSF, and Through Focus Spot tools. Zemax innately uses geometric ray tracing in most all of its operations like beam optimization. This is generally-acceptable for most optical systems; however, as our output light sheet size approaches the diffraction limit ( $\frac{\lambda}{2NA}$ ), we need to make sure to also consider the effects of diffraction in our analysis.

The Huygen's PSF analysis tool is how we incorporate effects of diffraction into our analysis; where we anticipate results from this analysis to be more in-line with what would be seen on the physical system. Based on the cross section of our Huygen's PSF analysis, we can see that our expected Full-Width Half-Max (FWHM) of the light sheet is expected to lie somewhere around 0.376 $\mu m$.

We compare the results of these two analyses for our optimized illumination path below, where we show the full XY profile as well as cross-sections through the center row of both beam profiles. In this case, the FWHM of both analyses ends up being quite similar at ~0.37 $\mu m$.

Through Focus Spot analysis allows us to essentially see the evolution of the light sheet through the point of focus, where we can then estimate a sort of range where we expect the width of the light sheet to be thin enough for our imaging purposes, where the maximum usable light sheet width is the FWHM at the focus multiplied by $\sqrt{2}$. The optimized illumination path simulation files are available in the Zemax folder of our repository.

---

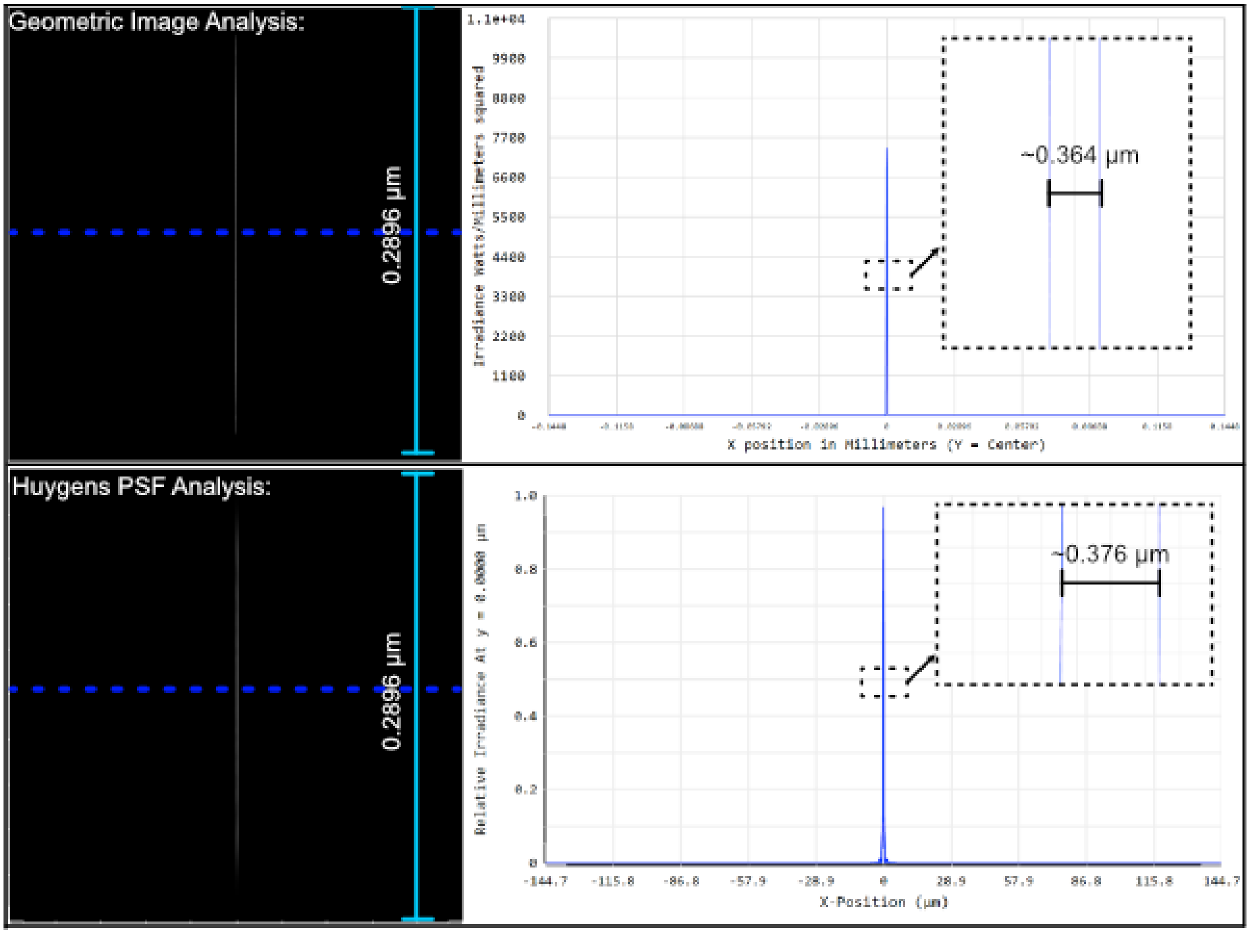

Fig. 5: **Figure 7:** A comparison of the Geometric Image Analysis and Huygen's PSF analysis for our optimized system. The FWHM of the light sheet is expected to be around 0.376 $\mu m$.

#### 5.4 Zemax Tolerance Analysis

When considering building physical systems using Zemax, an additional analysis tool known as tolerancing becomes increasingly important. No physical system is perfect, and tolerancing is essentially the process of understanding how sensitive different elements in a system are to various perturbations. This can be along the lines of how sensitive the collimation or magnification of a 4F system is to small physical displacements of the two lenses that comprise it. Similarly to Zemax's optimization process, tolerancing also utilizes a merit function. This merit function is fully customizable, and serves to define how well a particular system is performing. In the case of our system, we chose our merit function to factor in both the size and displacement of the output light sheet relative to the perfectly optimized instance. Our merit function used in Zemax is also shown below, where there are 4 operands that track the size and position of the beam in both x and y.

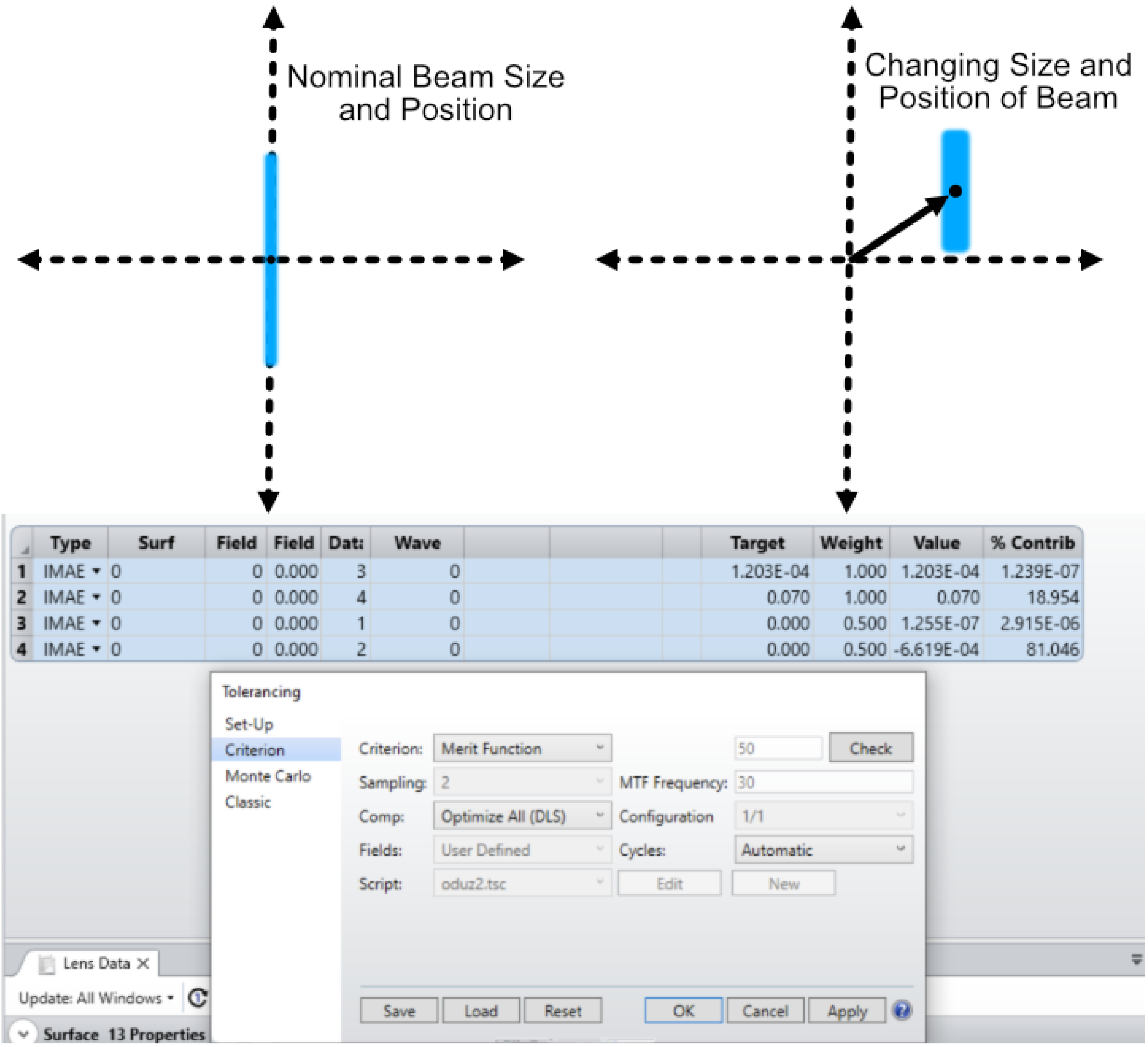

Fig. 6: **Figure 8:** The merit function used in our Zemax tolerancing analysis. The merit function is used to evaluate the performance of the system based on the size and position of the output light sheet.

With a merit function criteria set, the next step is to designate which elements of the system will change and by how much. In our case, we wanted to associate our tolerance analysis with the machining tolerances given by fabrication companies. In general, looking across different companies, the standard machining tolerance is around +-0.005” and the finer machining tolerance is around +-0.002”. For our analysis, we wanted to understand how angular deviations in elements due to machining tolerances in the alignment dowel pins would affect overall system performance. This is depicted below, where in the worst case scenario of one pin being offset +0.005” and the other -0.005” the resulting angular offset would be around 1.45 degrees.

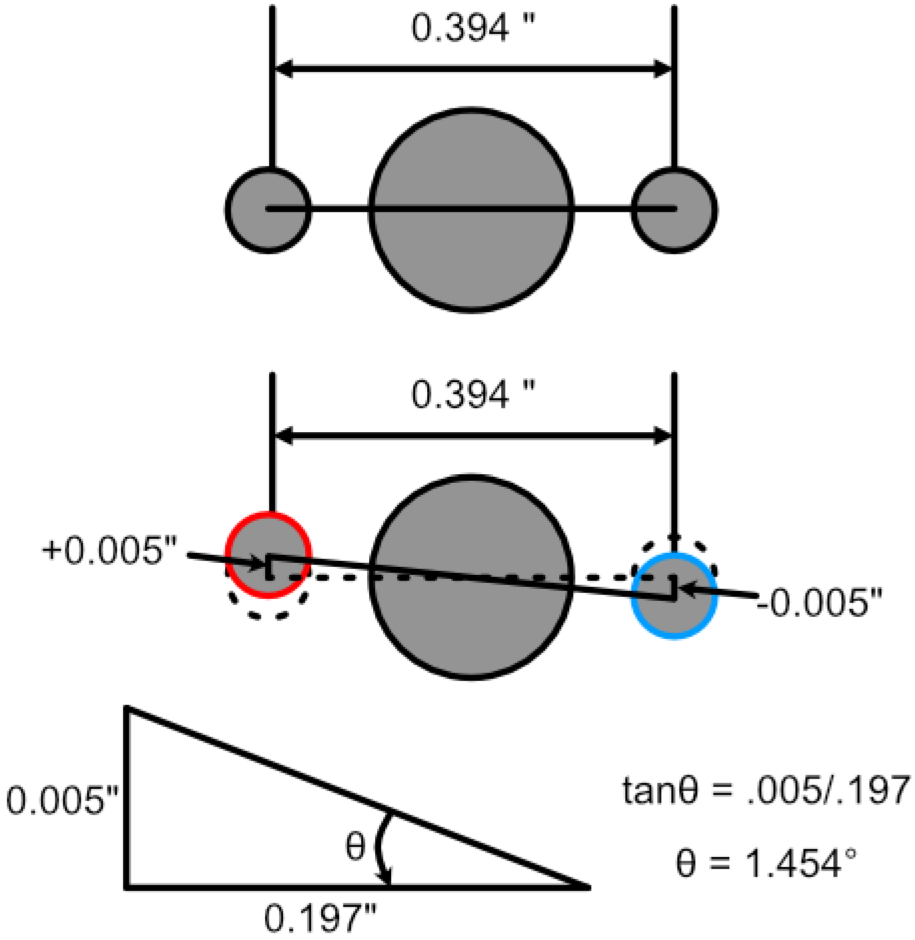

Fig. 7: **Figure 9:** The angular offset of elements due to machining tolerances of alignment dowel pin holes. In the worst case scenario, the angular offset is around 1.45 degrees.

In addition to perturbations to a system, in tolerancing analysis a compensator can also be defined as a sort of designated element that can be changed in ways to try to mitigate effects of other elements in the system being tweaked. In our case, we define the xy position of our illumination objective as a compensator with a range of +-0.25mm, which matches the xy translation adjustment associated with our objective mount used.

The basic way in which this analysis works is that Zemax performs a designated number of Monte Carlo simulations, each with a different perturbation made to the system, and evaluates the merit function for each of those systems. Based on the change to the merit function for each of these instances, tolerancing outputs a report that describes the sensitivity of the merit function to each of the different elements in the system. In some cases, tolerancing analysis gives information as to how much . An example of this is for a lens designer tolerancing the radii or material properties of a lens to ensure it’s focal length stays above or below a certain value. For our system though, even with our designated merit function, it is difficult to directly ascribe a sort of cutoff value of the merit function as acceptable, and so we primarily use tolerancing analysis as a way to guide us as to general trends of sensitivity in the elements of our system.

This is shown below, where in this instance we can see that in the case of our system, the element corresponding to the

24th surface (the galvo mirror) causes the most change to the merit function as it becomes perturbed. In all cases, the largest perturbations in the system (i.e. when the angular offset of an element is maximum at +-1.45 degrees) results in the largest changes to the merit function.

We also set our tolerance analysis to output the best and worst instances from the Monte Carlo simulations as individual files, and the corresponding geometric image analysis windows are shown for each as well as the nominal optimized case for comparison. It's clear that in the worst case scenario, it looks like the resulting light sheet is shorter in span than that of the nominal and best cases.

| | | \|----------- ---- Minimum -------------\| | | | \|------------ ---- Maximum ---- ---------\| | |
|---|---|---|---|---|---|---|---|
| Type | | Value | Criterion | Change | Value | Criterion | Change |
| TUTY | 3 | -1.45400000 | 0.00029706 | 0.00016637 | 1.45400000 | 0.00021864 | 8.7953E-05 |
| TUTY | 10 | -1.45400000 | 8.9203E-05 | -4.1483E-05 | 1.45400000 | 6.6077E-05 | -6.4608E-05 |
| TUTY | 18 | -1.45400000 | 0.00011300 | -1.7687E-05 | 1.45400000 | 0.00010052 | -3.0168E-05 |
| TUTY | 24 | -1.45400000 | 0.01370568 | 0.01357499 | 1.45400000 | 0.01335287 | 0.01322218 |
| TUTY | 37 | -1.45400000 | 0.00015180 | 2.1119E-05 | 1.45400000 | 0.00014712 | 1.6439E-05 |

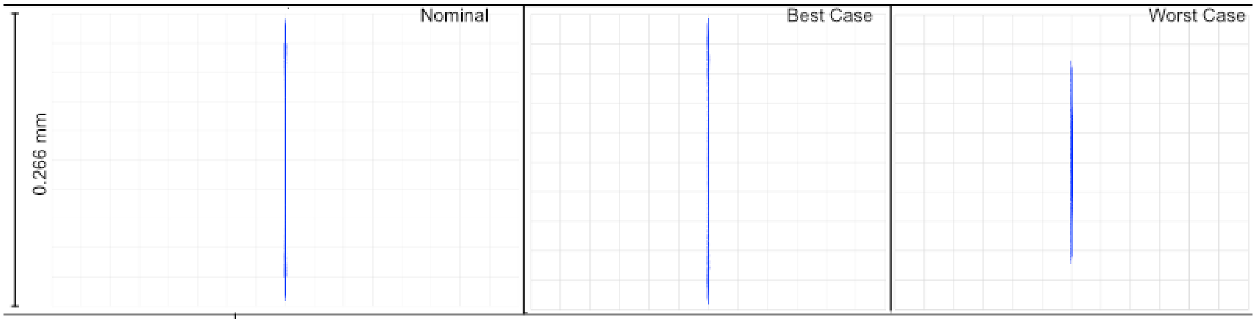

| | |
|---|---|
| 98% > | 0.01021605 |
| 90% > | 0.00651932 |
| 80% > | 0.00450740 |
| 50% > | 0.00168385 |
| 20% > | 0.00013947 |
| 10% > | 9.9676E-05 |
| 2% > | 2.8750E-05 |

| | | | |
|---|---|---|---|
| Nominal | 0.00013069 | | |
| Best | 4.3493E-06 | Trial | 452 |
| Worst | 0.01648946 | Trial | 243 |
| Mean | 0.00252931 | | |
| Std Dev | 0.00274570 | | |

Fig. 8: **Figure 10:** Results of the tolerancing analysis when the offset corresponded to +-0.005". The merit function is used to evaluate the performance of the system based on the size and position of the output light sheet.

To understand how tighter tolerances might affect system performance, we set our angular offset to correspond to tighter machining tolerances offered online at +-0.002". Typically, tighter machining tolerances correspond to an increase in price, so understanding if higher tolerances would benefit a system is beneficial. We can the same tolerance analysis as before, but this time with an angular offset of +-0.581 degrees, and show the results below. In this analysis, once again the element that affects the system most adversely is the galvo mirror element. The deviations in the resulting merit functions from this element are about a tenth of that of the larger machining tolerance case. Visually, in the worst case example, one can see that the resulting light sheet looks much closer to the nominal case than before as well.

The results of our tolerancing analysis, as well as the associated lens files for our best and wort case instances for both fine and coarse tolerancing are available here.

| Type | | Minimum Value | Minimum Criterion | Minimum Change | Maximum Value | Maximum Criterion | Maximum Change |
|---|---|---|---|---|---|---|---|
| TUTY | 3 | -0.58100000 | 5.7648E-05 | -7.3037E-05 | 0.58100000 | 9.4611E-05 | -3.6074E-05 |
| TUTY | 10 | -0.58100000 | 0.00011160 | -1.9086E-05 | 0.58100000 | 0.00010886 | -2.1820E-05 |
| TUTY | 18 | -0.58100000 | 0.00012106 | -9.6250E-06 | 0.58100000 | 0.00011500 | -1.5686E-05 |
| TUTY | 24 | -0.58100000 | 0.00278300 | 0.00265232 | 0.58100000 | 0.00259014 | 0.00245946 |
| TUTY | 37 | -0.58100000 | 0.00013506 | 4.3707E-06 | 0.58100000 | 0.00012988 | -8.0569E-07 |

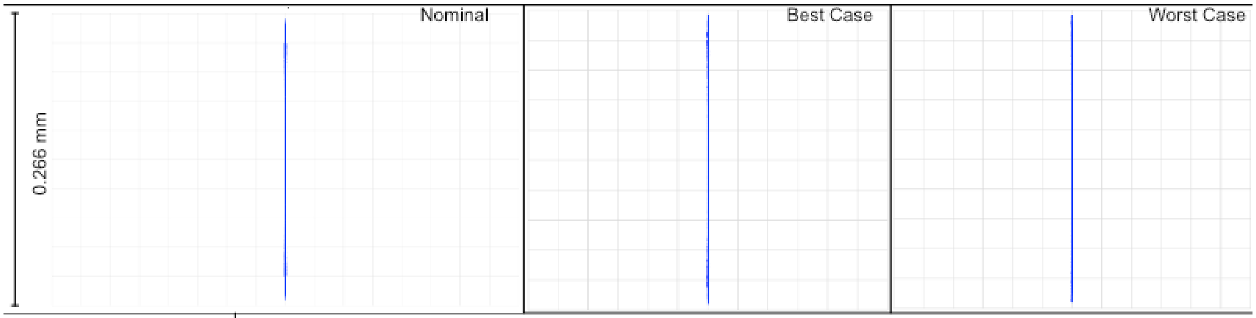

| | |
|---|---|
| 98% > | 0.00250521 |
| 90% > | 0.00176802 |
| 80% > | 0.00118280 |
| 50% > | 0.00028634 |
| 20% > | 9.9463E-05 |
| 10% > | 5.9625E-05 |
| 2% > | 1.4822E-05 |

| | | | |
|---|---|---|---|
| Nominal | 0.00013069 | | |
| Best | 9.6097E-07 | Trial | 705 |
| Worst | 0.00302942 | Trial | 739 |
| Mean | 0.00062233 | | |
| Std Dev | 0.00070550 | | |

Fig. 9: **Figure 11:** Results of the tolerancing analysis when the offset corresponded to +-0.002”. The merit function is used to evaluate the performance of the system based on the size and position of the output light sheet.

#### 5.5 Baseplate Design

When satisfied with the results of simulations, the optimized values in Zemax can then be used to design our baseplate. This process involves taking the optimized distances between our various optical elements and then considering how each of those elements are mounted in a physical system, as in Zemax all of the elements are effectively suspended in midair like below:

For mounting our elements, we utilize the Polaris line from Thorlabs, which are designed with long-term stability and alignment in mind. Each component is characterized in part by two dowel pin alignment holes to ensure subsequent mounted elements are aligned along a specific axis. In the baseplate design, we are essentially deciding on the location for the mounting holes of the Polaris posts we’re using, which is not the same as the locations of the elements themselves from Zemax.

While we are able to use most of our element mounts from the Polaris line, for the cylindrical lens L3 we needed a mount capable of rotating the lens, which at this time is not something available from Thorlabs. In our case we designed an additional mounting element that allows the use of a basic Thorlabs RSP1 rotation mount, but still ensures alignment with the other Polaris elements. The CAD file for this mount is available for download in our CAD directory.

With the method in which each of the elements needs to be mounted decided upon, we then went over the product schematics for each mount to understand the z-displacement that they impart upon the element mounted within them relative to where the Polaris post central mounting hole would need to be. This idea is depicted below, where when considering how to space two lenses from each other there is essentially three components to take into account:

1. The distance between the lenses decided from simulation
2. The thickness of the lenses themselves
3. The distance between the center of the Polaris post and the start of the lens in the mount

Once the locations of the mounting holes were determined, we used Autodesk Inventor to design the full baseplate. The

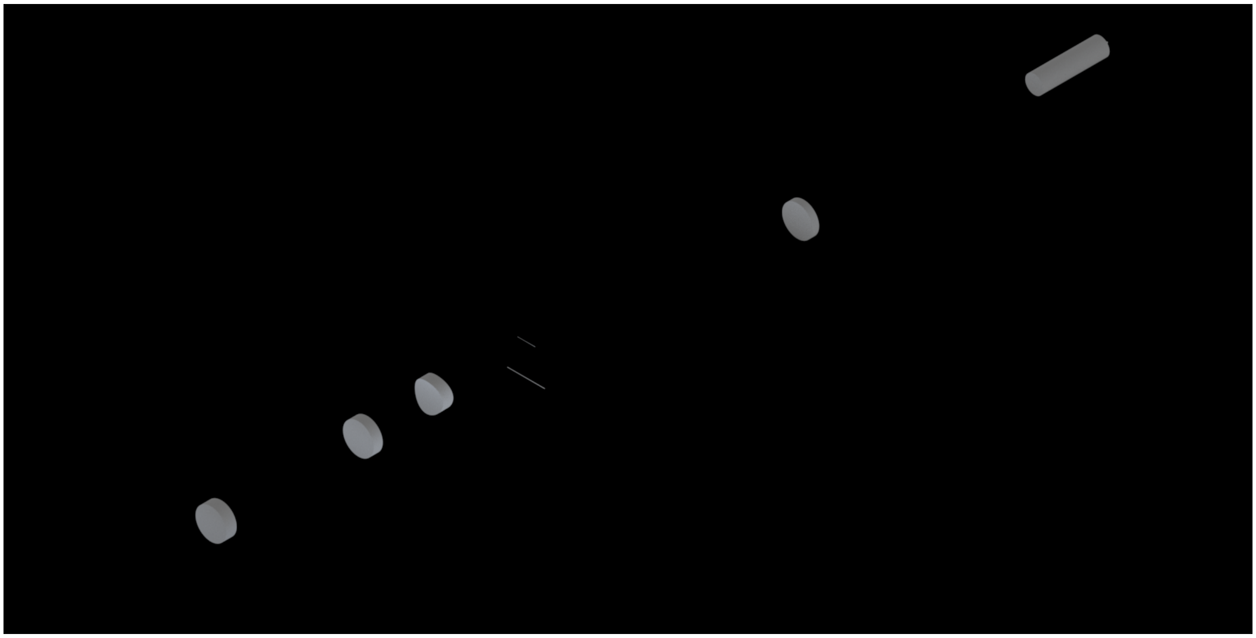

Fig. 10: **Figure 12:** A Zemax diagram of the elements in our system. Here, the elements are shown floating in midair.

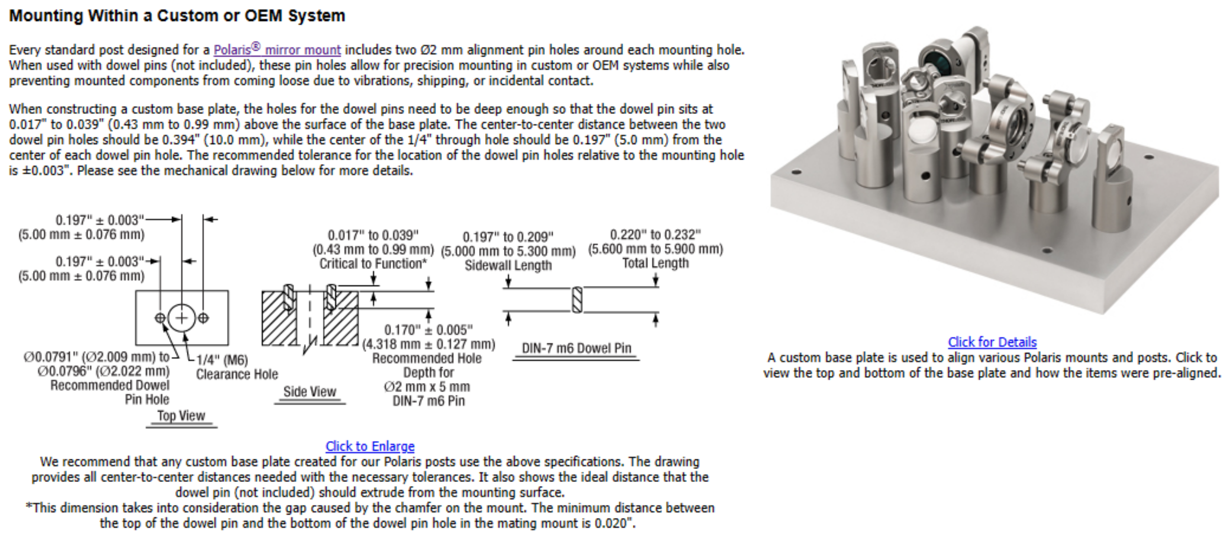

Fig. 11: **Figure 13:** A schematic of the Polaris mounting system. The system is characterized by two dowel pin alignment holes to ensure subsequent mounted elements are aligned along a specific axis.

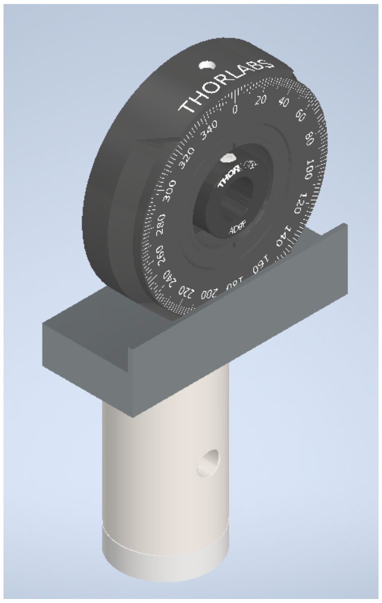

Fig. 12: **Figure 14:** A schematic of the rotation mount adapter. The adapter allows for the rotation of the cylindrical lens while ensuring alignment with the other Polaris elements.

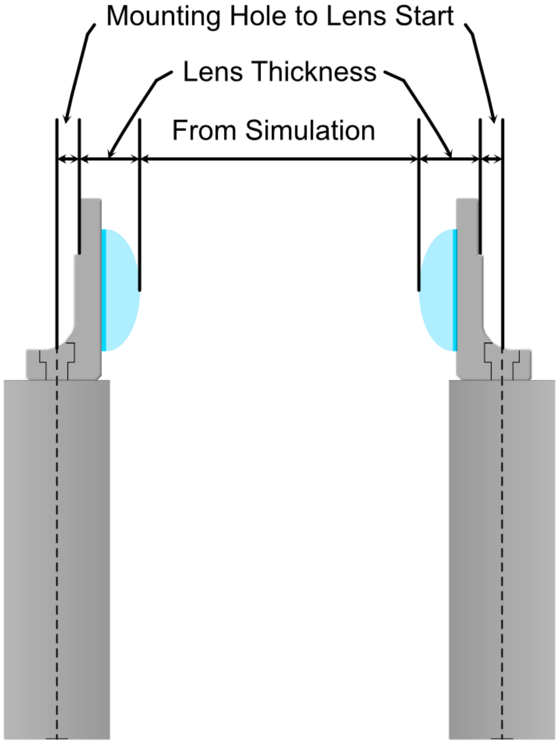

Fig. 13: **Figure 15:** A schematic of the considerations for spacing two lenses from each other. The distance between the lenses is decided from simulation, the thickness of the lenses themselves, and the distance between the center of the Polaris post and the start of the lens in the mount.

baseplate is essentially just a mounting hole and the two dowel pin holes for every element, as well as four mounting holes for the baseplate itself. These four baseplate mounting holes were spaced in increments of inches such that the baseplate can either be screwed directly into an optical breadboard table or into additional posts that can keep the assembly at a desired height.

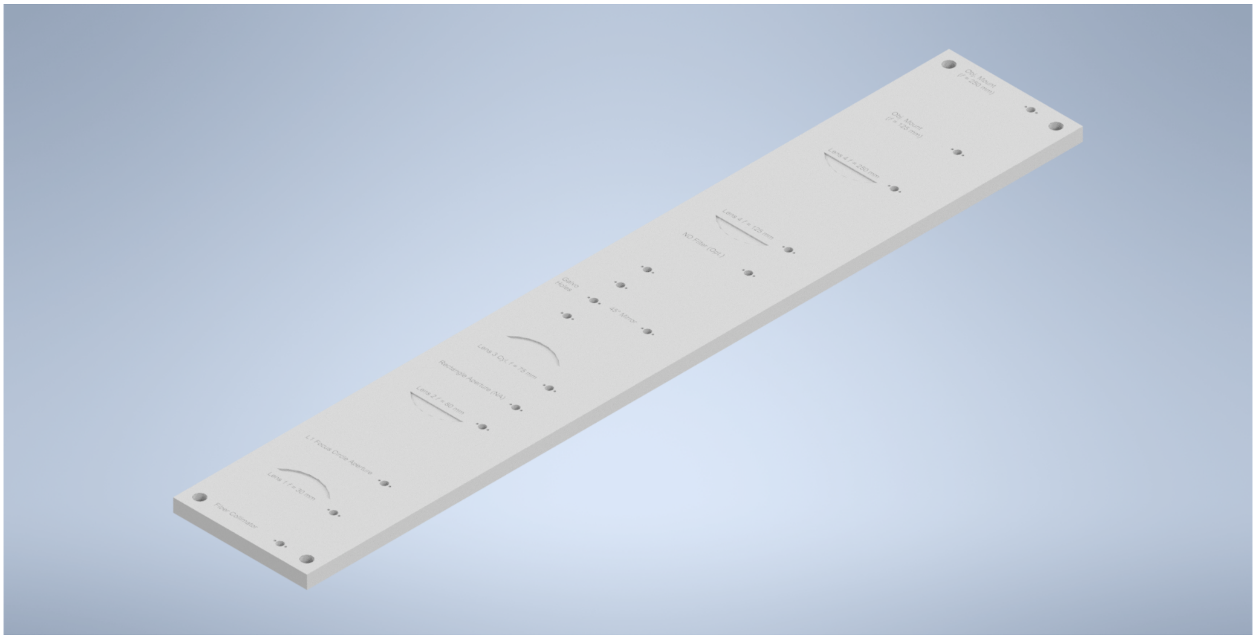

Fig. 14: **Figure 16:** A schematic of the baseplate. The baseplate is characterized by mounting holes for each element and four mounting holes for the baseplate itself.

With the baseplate designed, our final assembly for our illumination path looks as follows:

The CAD files for our baseplate design are available in the following GitHub repository.

---

#### 5.6 Physical Coordinate Definitions

It should be noted briefly that when discussing our physical microscope systems using navigate software, the definitions for the coordinate axes is different than that of our simulations. This is due to a difference in standardized definitions for the axes in our previous systems and how Zemax defines these same axes. This difference is depicted in the picture below:

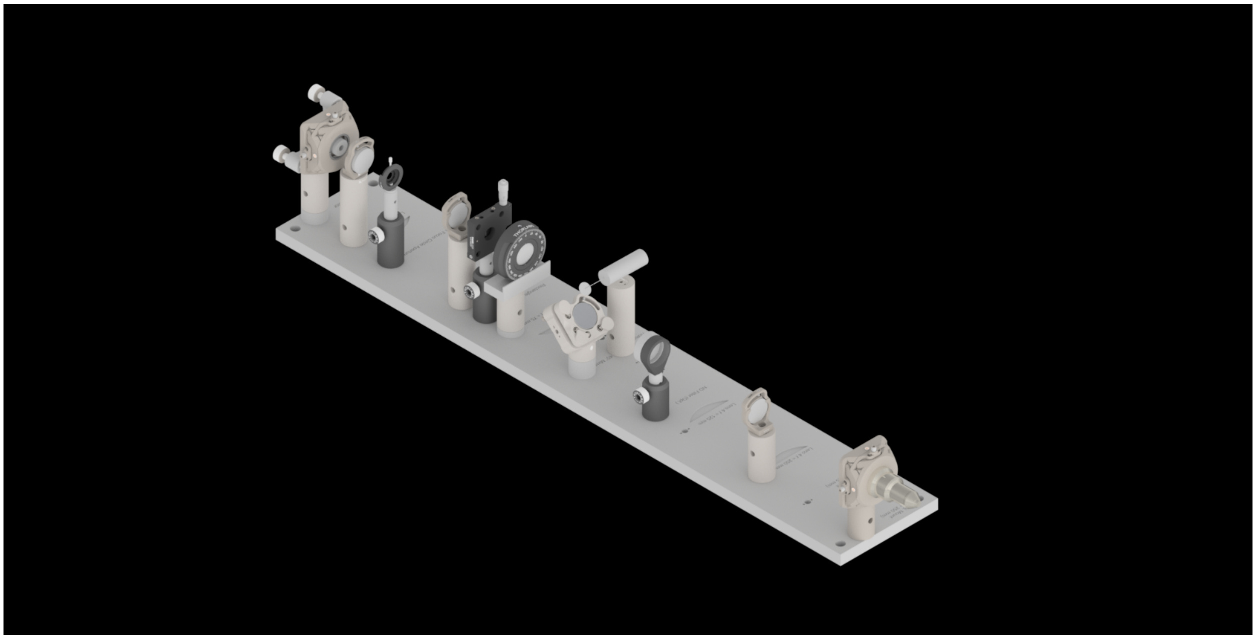

Fig. 15: **Figure 17:** An isometric view of the baseplate assembly.

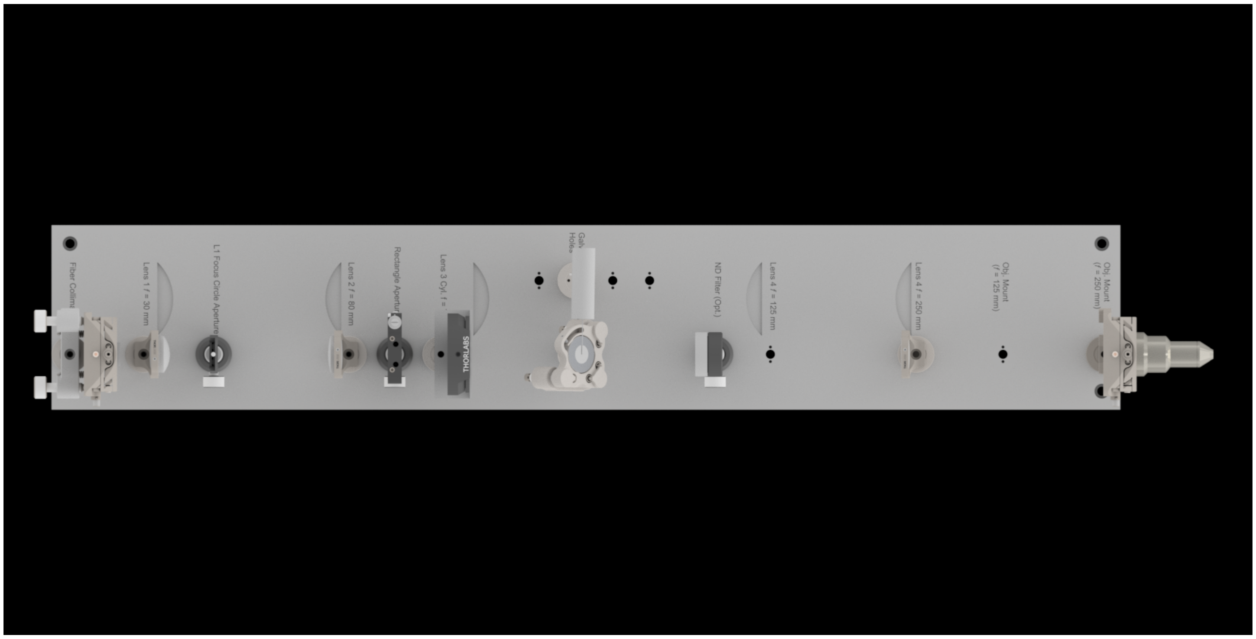

Fig. 16: **Figure 18:** A top view of the baseplate assembly.

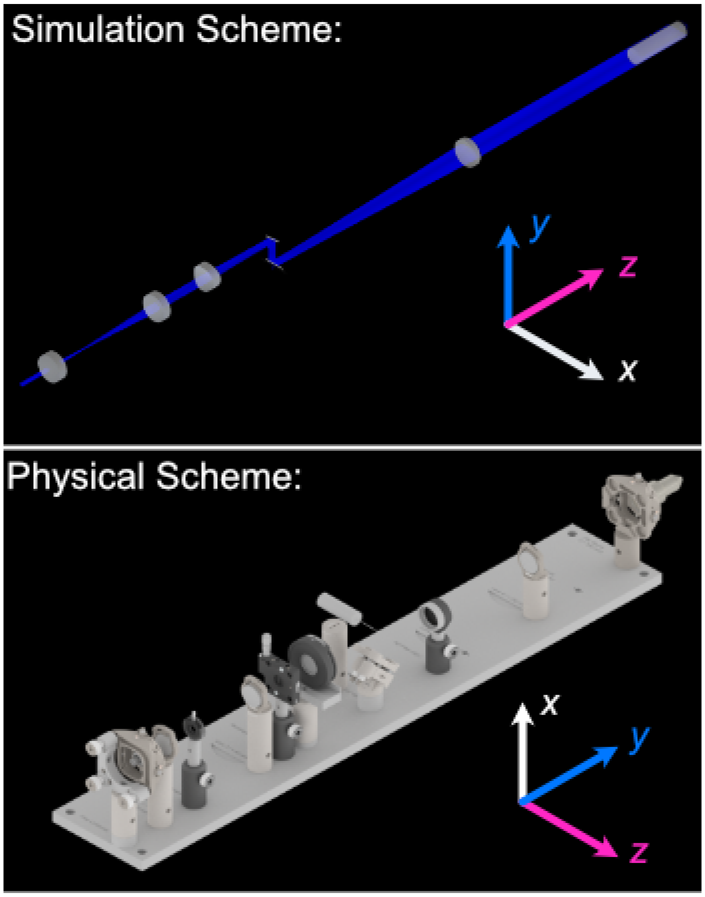

Fig. 17: **Figure 19:** A schematic of the difference in coordinate axes for simulation and physical setup.

CHAPTER

SIX

### MICROSCOPE ASSEMBLY

#### 6.1 Parts List and Cost

A breakdown of all components used in Altair, and the approximate cost are included in the collapsable sections below:

| Standard Parts | Vendor | Purpose |
|---|---|---|
| TL20X-MPL | Thorlabs | Illumination Objective |
| Polaris-1XY | Thorlabs | Mounting |
| Polaris-K1XY | Thorlabs | Mounting |
| Polaris-MA45 | Thorlabs | Mirror Mounting |
| Polaris-P150 | Thorlabs | Mounting |
| Polaris-P200 | Thorlabs | Mounting for L4 |
| Polaris-P3 | Thorlabs | Mounting |
| Polaris-P250 | Thorlabs | Mounting |
| Polaris-P225 | Thorlabs | Mounting |
| Polaris-P075 | Thorlabs | Mounting |
| Polaris-P175 | Thorlabs | Mounting |
| Polaris-B1S | Thorlabs | Lens Mounting |
| Polaris-K1S4 | Thorlabs | Mirror Mounting |
| CFC11A-A | Thorlabs | Fiber Laser Collimator |
| P3-405B-FC-1 | Thorlabs | Single Mode Fiber |
| AC254-080-A-ML | Thorlabs | L2 |
| AC254-030-A-ML | Thorlabs | L1 |
| VA100CP | Thorlabs | Rectangular Aperture |
| ACY254-075-A | Thorlabs | L3 |
| PF10-03-P01 | Thorlabs | 1" Mirror |
| RSP1 | Thorlabs | Rotation Mount For Cylindrical Lens |
| AC254-250-A-ML | Thorlabs | L4 |
| SM1A12 | Thorlabs | Illumination Objective Mounting |
| 1-Axis 4kHz Resonant Galvo and Servo | Novanta | Galvo Mirror |
| TGPOW-12-3 | ASI | Galvo Low Noise Power Supply +12V 3A |
| Polaris B1F | Thorlabs | Mount for Mirror 2 |
| ID12 | Thorlabs | Mounted Circular Iris (12 mm Max Aperture) |
| L4CC | Oxxius | Multicolor Laser Source |
| **Custom Machined Parts** | **Vendor** | **Purpose** |
| RSP1 Polaris Adapter | Xometry | Polaris Adapter for RSP1 Cylindrical Lens |
| Illumination Baseplate | Xometry | Baseplate |
| Galvo Holder | Protolabs | Galvo Mounting |

continues on next page

Table 1 – continued from previous page

| Standard Parts | Vendor | Purpose |
|---|---|---|
| VA100CP Polaris Adapter | Xometry | Aperture Mounting |
| IDA12 Polaris Adapter | Xometry | Polaris Adapter for ID12 Iris |

| Part | Vendor | Purpose |
|---|---|---|
| N25X-APO-MP | Thorlabs (Nikon) | Detection Objective |
| LS-100-AMCCH | ASI | 100 mm Linear Focusing Stage |
| TGDCM2 | ASI | 2-Axis Stage Control Card |
| C60-EXT-15 | ASI | 15 mm Tube Extension |
| RAO-0051 | ASI | M32x0.75 Threaded Sleeve |
| FW-0002-8 | ASI | 8-Position Filter Wheel |
| FW-C-MNT-K1 | ASI | Filter Wheel to MIM Adapter Kit |
| TGFW | ASI | Filter Wheel Control Card |
| C60-TUBE-400 | ASI | 400 mm Achromatic Tube Lens |
| C13440-20CU | Hamamatsu | ORCA Flash4.0 V3 Camera |
| Semrock Brightline Filter 605/15-25nm | IDEX | Filter |
| Semrock Brightline Filter 445/20nm | IDEX | Filter |
| Semrock Brightline Filter 676/29 nm | IDEX | Filter |
| Semrock Brightline Filter 529/24-25nm | IDEX | Filter |

| Part | Vendor | Purpose |
|---|---|---|
| KBT1X1T | Thorlabs | Magnetic Mount for Sample Holder Top |
| KBB1X1 | Thorlabs | Magnetic Mount for Sample Holder Bottom |
| LS-5012 | ASI | Breadboard Adapter for Translation Stages |
| LS-5013 | ASI | Right Angle Bracket for Translation Stages |
| DV-6010-C | ASI | Dovetail Mount Pair |
| LS-50-AMCLLS | ASI | 50 mm Linear Stage with Stainless Steel Slide |
| LS-100-AMERL | ASI | 100 mm Linear Stage, 16 TPI, Extended, Right |
| LS-50-AMELL | ASI | 50 mm Linear Stage |
| TGADEPT | ASI | Piezo Control Card |
| HS1.100 | PiezoConcept | 100 Micron Piezo Stage |
| ADAPTHS1BB | PiezoConcept | Adapter Plate for PiezoConcept HS1 |
| **Custom Machined Parts** | **Vendor** | **Purpose** |
| Angle Bracket Adapter | Xometry | Mounting the piezo to ASI translation stages at an angle |
| 5mm Coverslip Holder | Xometry | Sample holder for 5 mm Coverslips |
| 5mm Coverslip Holder Adapter | Xomtery | Adapter to mount coverslip holder onto KBT1X1T linearly |

| Part | Vendor | Purpose |
|---|---|---|
| TG16-BASIC | ASI | Tiger Controller - 16 Bay System |
| T3283 | Thorlabs | BNC Adapter (F-F) |
| PAA272R | Thorlabs | SMB F to BNC M cable adapter |
| CA2106 | Thorlabs | BNC to Terminal Pin Cable |
| PCIe-6738 | NI | Data Acquisition Card |
| SHC68-68-A2 | NI | Test Cable Assembly |
| SCB-68A | NI | Noise Rejecting Terminal Block |
| SX6300 | Colfax International | Colfax SC6300 Workstation |
| 14UD-42X-24 | TMC | Ultradamp Vibration Isolation with Casters |
| 784-736-02R | TMC | 36x60x18" Performance Series Optical Top Table |

| Section | Cost (USD) |
|---|---|
| Illumination Path | $42,419.67 |
| Detection Path | $61,635.82 |
| Sample Positioning | $18,869.78 |
| Shared Equipment | $32,300 |
| **TOTAL** | **$155,224.78** |

#### 6.2 Illumination Path

Our baseplate design was made with ease of assembly in mind. The basic process involves aligning Polaris posts with dowel pins and screwing them using 1/4"-20 Screws in at the predetermined hole locations on the breadboard. This general process is depicted below:

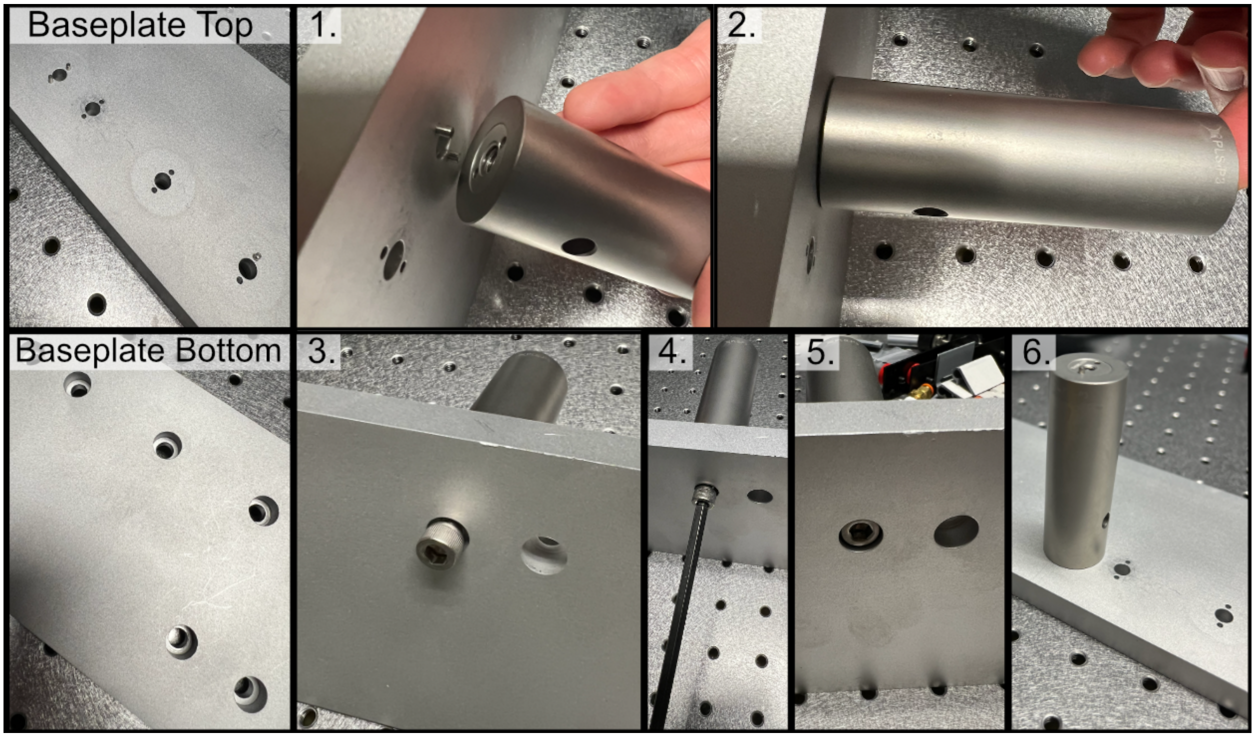

Fig. 1: **Figure 1:** General process for baseplate assembly

To either mount the baseplate onto an optical table or onto separate posts, the process is similar in that just requires screwing 1/4"-20 screws into either an optical breadboard or onto separate posts at the four corner holes. For our system, we opted to mount our illumination baseplate on 1.5" tall 1" diameter Thorlabs pedestal posts. As a general note, having the illumination path elevated instead of affixed directly into the table is both helpful to enable access to mounting and unmounting the illumination optics as well as ensuring that the illumination objective and detection objective are able to be aligned vertically with each other.

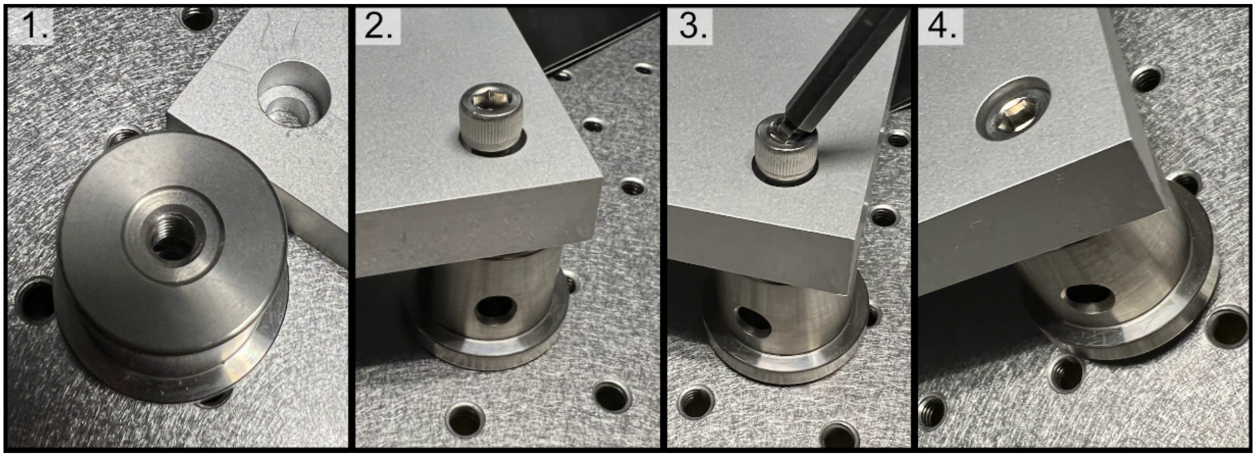

Fig. 2: **Figure 2:** General process to place posts on baseplate corners

##### 6.2.1 Ordering Custom Parts from Xometry

We utilize a number of different custom fabricated elements in our completed system, . There are a number of different companies available for this, and we opted for Xometry. The basic ordering process we go through is as follows:

1. **Begin by creating an account with Xometry**

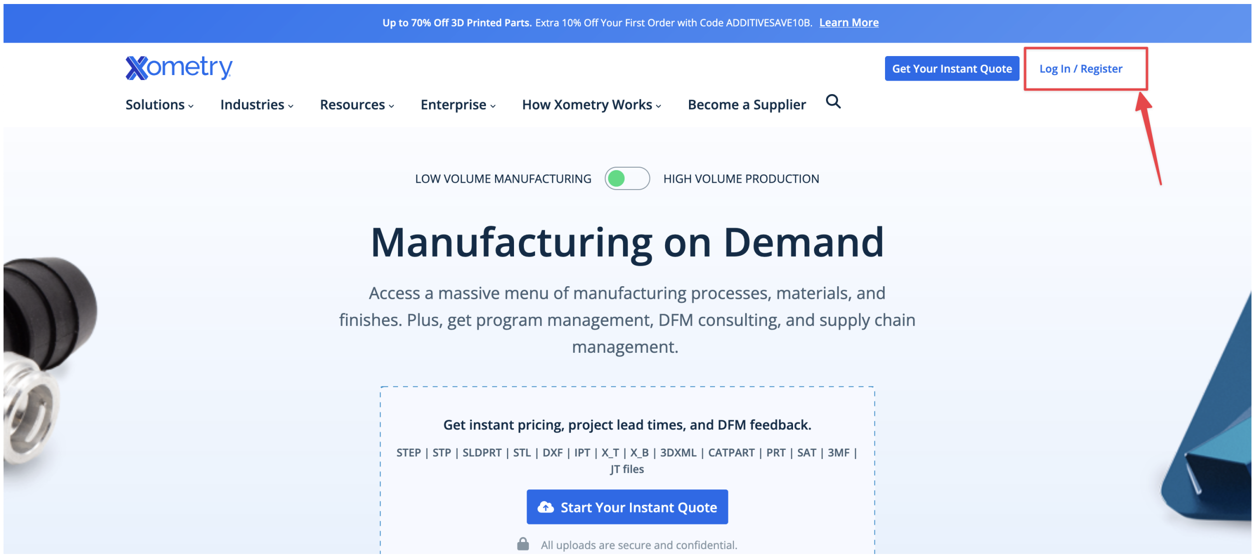

2. **Start a quote by uploading your desired file (in .stp format)**
3. **Edit the configuration**
4. **Adjust the fabrication process to be CNC machining**
5. **Select Aluminum 6061-T6x as the material for machining**
6. **Adjust the finish to be bead blasted**
7. **If you have a part that uses threads or tapped holes select that option and upload associated pdf drawings**
8. Select shipping details and checkout

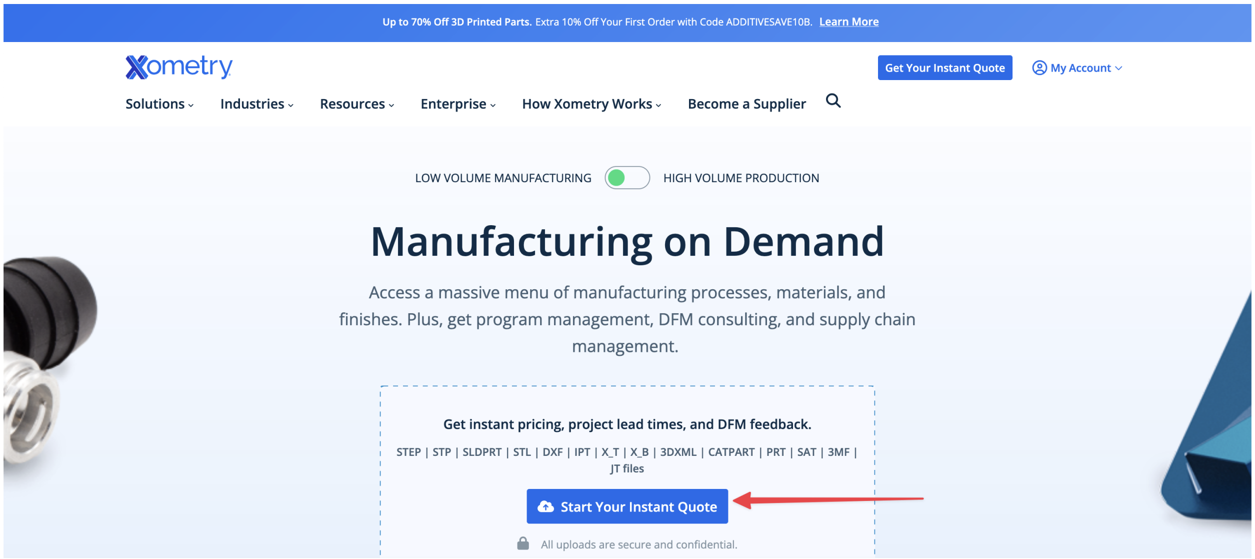
Up to 70% Off 3D Printed Parts. Extra 10% Off Your First Order with Code ADDITIVESAVE10B. Learn More
Xometry
Get Your Instant Quote
My Account
Solutions
Industries
Resources
Enterprise
How Xometry Works
Become a Supplier
LOW VOLUME MANUFACTURING
HIGH VOLUME PRODUCTION
Manufacturing on Demand
Access a massive menu of manufacturing processes, materials, and finishes. Plus, get program management, DFM consulting, and supply chain management.
Get instant pricing, project lead times, and DFM feedback.
STEP | STP | SLDPRT | STL | DXF | IPT | X_T | X_B | 3DXML | CATPART | PRT | SAT | 3MF | JT files
Start Your Instant Quote
All uploads are secure and confidential.

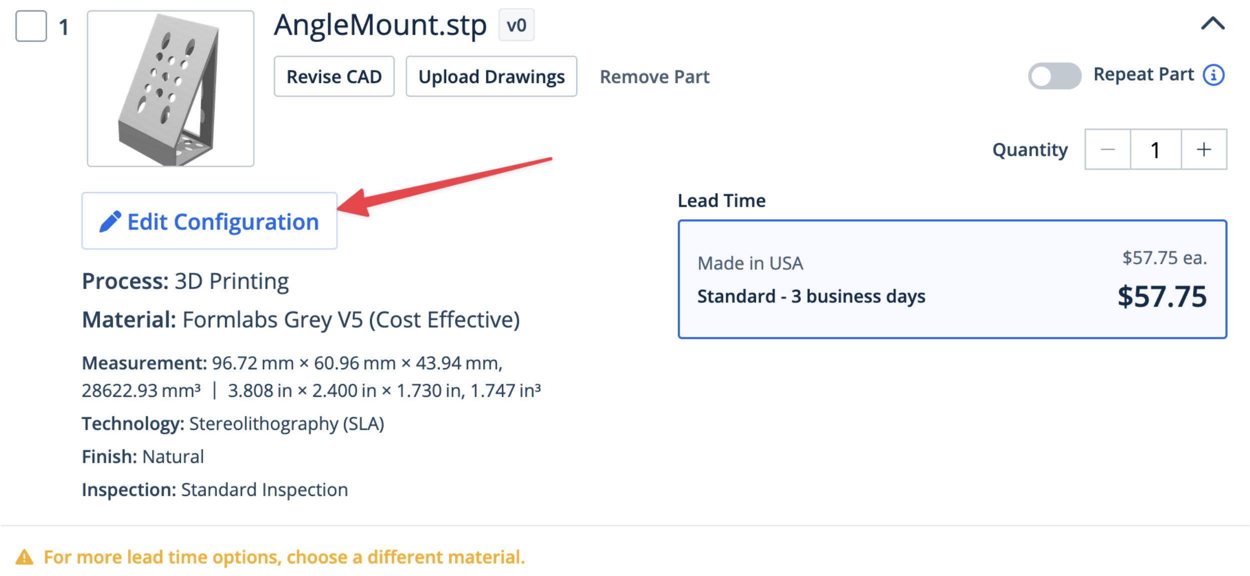
1
AngleMount.stp
v0
Revise CAD
Upload Drawings
Remove Part
Repeat Part
Quantity
1
Edit Configuration
Lead Time
Made in USA
$57.75 ea.
Standard - 3 business days
$57.75
Process: 3D Printing
Material: Formlabs Grey V5 (Cost Effective)
Measurement: 96.72 mm × 60.96 mm × 43.94 mm, 28622.93 mm³ | 3.808 in × 2.400 in × 1.730 in, 1.747 in³
Technology: Stereolithography (SLA)
Finish: Natural
Inspection: Standard Inspection
For more lead time options, choose a different material.

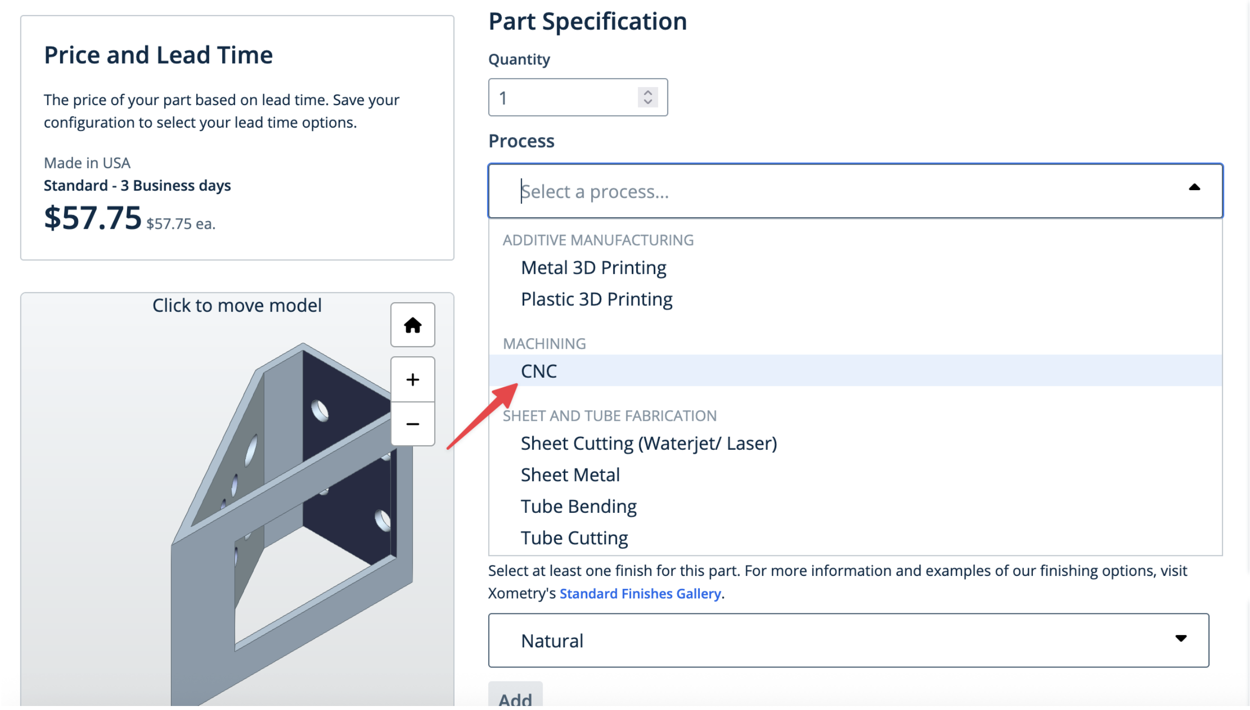
Price and Lead Time
The price of your part based on lead time. Save your configuration to select your lead time options.
Made in USA
Standard - 3 Business days
$57.75 $57.75 ea.
Click to move model
Part Specification
Quantity
1
Process
Select a process...
ADDITIVE MANUFACTURING
Metal 3D Printing
Plastic 3D Printing
MACHINING
CNC
SHEET AND TUBE FABRICATION
Sheet Cutting (Waterjet/ Laser)
Sheet Metal
Tube Bending
Tube Cutting
Select at least one finish for this part. For more information and examples of our finishing options, visit Xometry's Standard Finishes Gallery.
Natural
Add

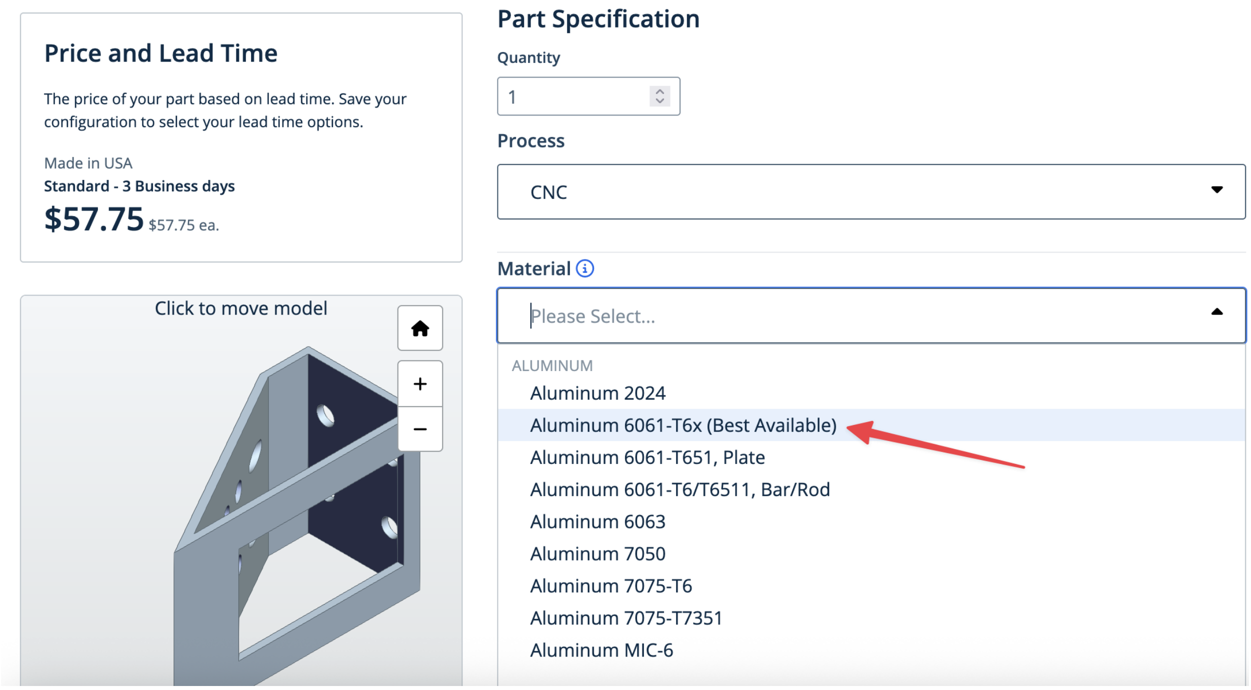
Price and Lead Time
The price of your part based on lead time. Save your configuration to select your lead time options.
Made in USA
Standard - 3 Business days
$57.75 $57.75 ea.
Click to move model
Part Specification
Quantity
1
Process
CNC
Material
Please Select...
ALUMINUM
Aluminum 2024
Aluminum 6061-T6x (Best Available)
Aluminum 6061-T651, Plate
Aluminum 6061-T6/T6511, Bar/Rod
Aluminum 6063
Aluminum 7050
Aluminum 7075-T6
Aluminum 7075-T7351
Aluminum MIC-6

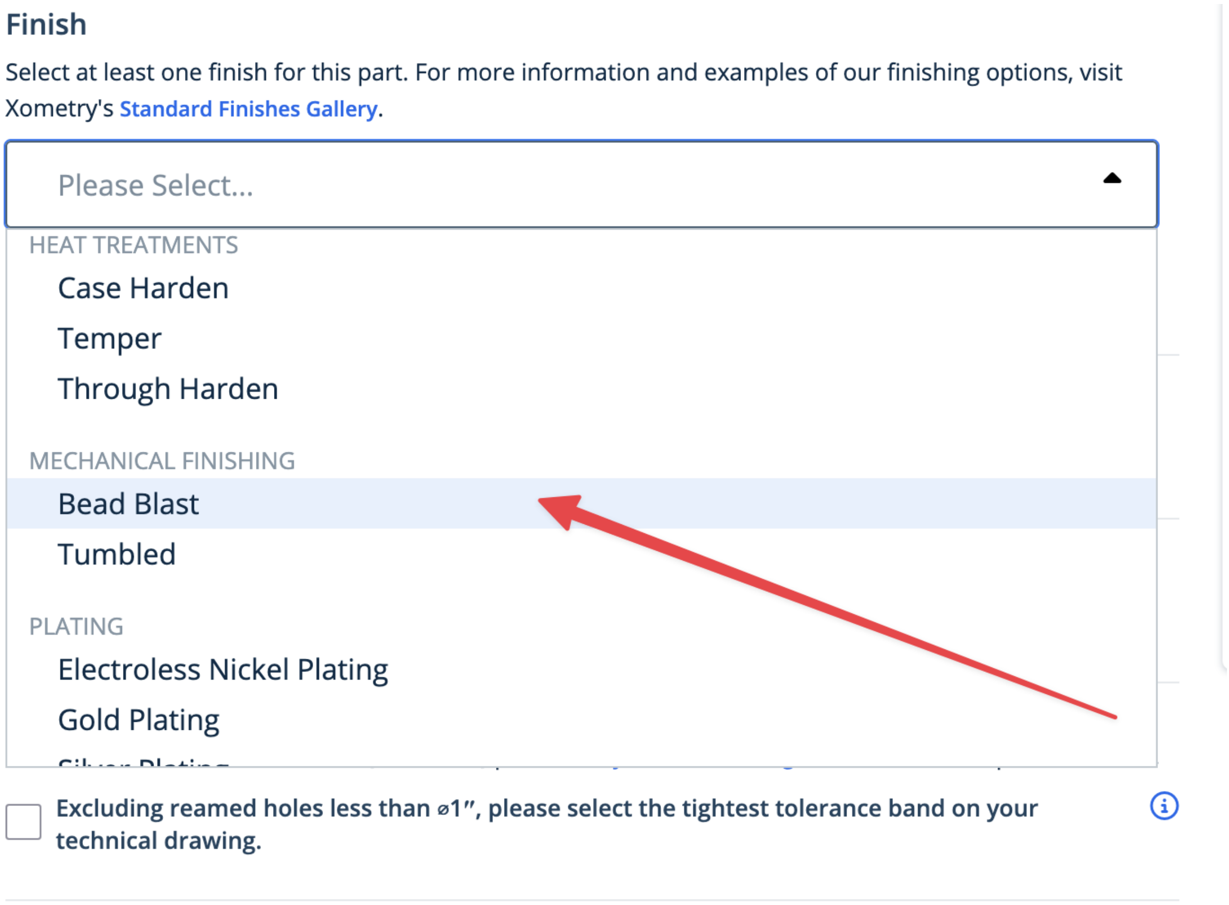
Finish
Select at least one finish for this part. For more information and examples of our finishing options, visit Xometry's Standard Finishes Gallery.
Please Select...
HEAT TREATMENTS
Case Harden
Temper
Through Harden
MECHANICAL FINISHING
Bead Blast
Tumbled
PLATING
Electroless Nickel Plating
Gold Plating
Excluding reamed holes less than ⌀1″, please select the tightest tolerance band on your technical drawing.

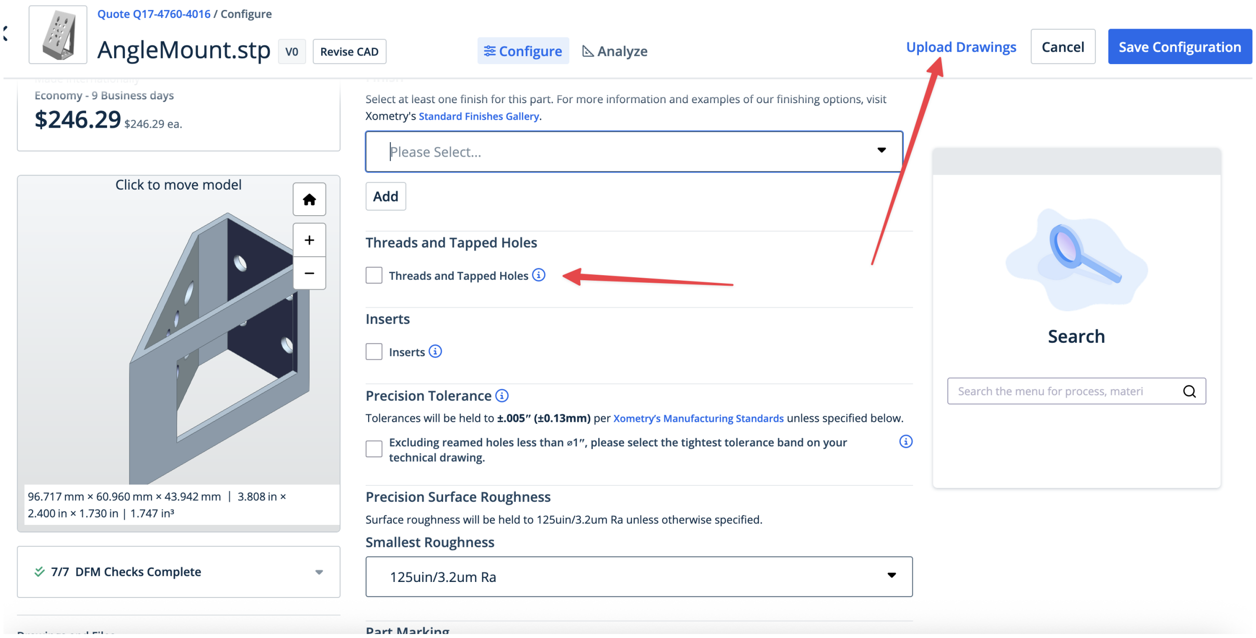
Quote Q17-4760-4016 / Configure
AngleMount.stp
V0
Revise CAD
Configure
Analyze
Upload Drawings
Cancel
Save Configuration
Economy - 9 Business days
$246.29 $246.29 ea.
Click to move model
96.717 mm × 60.960 mm × 43.942 mm | 3.808 in × 2.400 in × 1.730 in | 1.747 in³
7/7 DFM Checks Complete
Select at least one finish for this part. For more information and examples of our finishing options, visit Xometry's Standard Finishes Gallery.
Please Select...
Add
Threads and Tapped Holes
Threads and Tapped Holes
Inserts
Inserts
Precision Tolerance
Tolerances will be held to ±.005″ (±0.13mm) per Xometry's Manufacturing Standards unless specified below.
Excluding reamed holes less than ⌀1″, please select the tightest tolerance band on your technical drawing.
Precision Surface Roughness
Surface roughness will be held to 125uin/3.2um Ra unless otherwise specified.
Smallest Roughness
125uin/3.2um Ra
Part Marking
Search
Search the menu for process, materi

##### 6.2.2 Mounting Lenses

Mounting lenses into a Polaris lens mount and onto an associated post is a fairly straightforward process. The general flow is shown in the image below, where first the flatter face of the desired lens should be placed such that it is touching the metal boundary on the lens mount itself. Then the lens should be fixed into place by screwing the lock screw on the top of the lens mount. With the lens secured in the mount, then two dowel pins should be placed in the appropriate holes on the Polaris post, and then the lens mount should be placed such that the two holes on the lens mount align with the pins on the Polaris post. Then the lens mount should be anchored into place by screwing it into the Polaris post. Additional information can be found at Thorlabs.

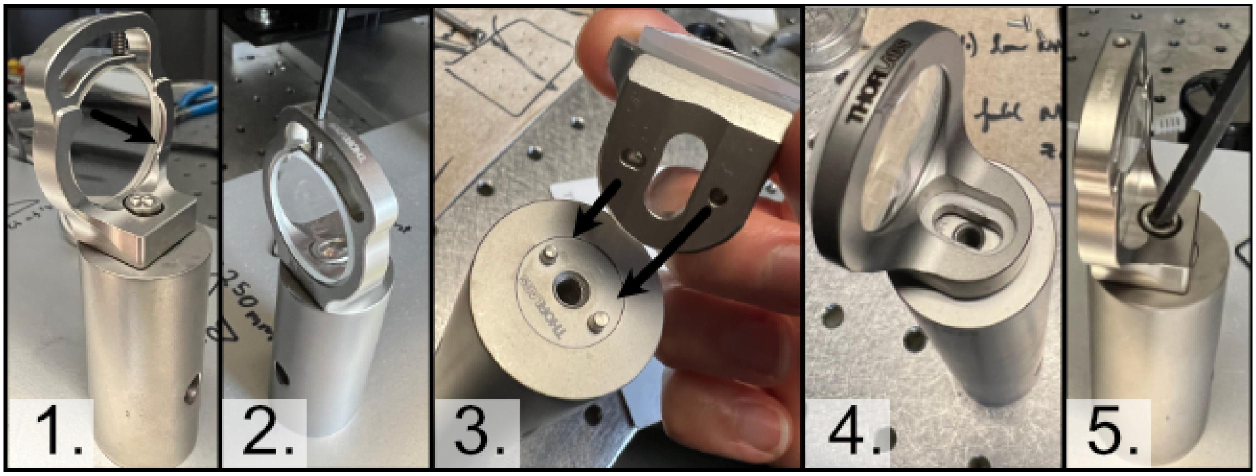

Fig. 3: **Figure 3:** General process for mounting a lens into a Polaris holder and onto a post

##### 6.2.3 Initial Laser Collimation and Alignment

When first assembling the system, ensuring proper output collimation from the fiber laser source is critical. There are multiple checks that one can take for this step, but we utilize a combination of a shear-plate interferometer and two pinhole apertures placed at opposite ends along the length of the baseplate. Shear-plate interferometers are designed to split and interfere an input beam of coherent light, such that when the beam is collimated there are interference fringes aligned vertically with a reference line. The fiber laser collimator we used for this system is the Thorlabs CFC11A-A, which features an adjustable barrel which controls the position of collimation optics within the element.

The basic assembly process involves first inserting and fixing the CFC11A-A into a Thorlabs AD15S2 adapter, which allows it to then be mounted into a 2.5" Polaris K1XY mount. This assembly is then mounted onto the respective Polaris post at the start of the baseplate. The fiber laser source is then able to be directly mounted into the CFC11A-A, making sure that the protrusion on the fiber wire aligns with the open section of the CFC11A-A port. The basic process of ensuring collimation then involves turning on the laser source, and placing the shear-plate interferometer such that the input port aligns with the output of the laser unit. Then, by slowly adjusting the barrel of the CFC11A-A and observing the interference fringe orientations along the top display of the interferometer, one is able to adjust the beam until it is properly collimated.

With the beam collimated, the process of beam alignment involves adjusting the position control knobs on the K1XY to have the beam pass through two pinhole apertures along the optical path. The height of the initial laser output is designed to be at 3.75" above the top surface of the baseplate, so selecting appropriate post heights for the apertures such that their centers rest at 3.75" is essential. In our case, we use Thorlabs ID12 pinhole apertures, so using a post height of 3.25" will ensure that they are at the proper height for alignment. We designed a custom ID12 to Polaris adapter to ensure the aperture is at the proper height and properly aligned along the designated Polaris axis. When

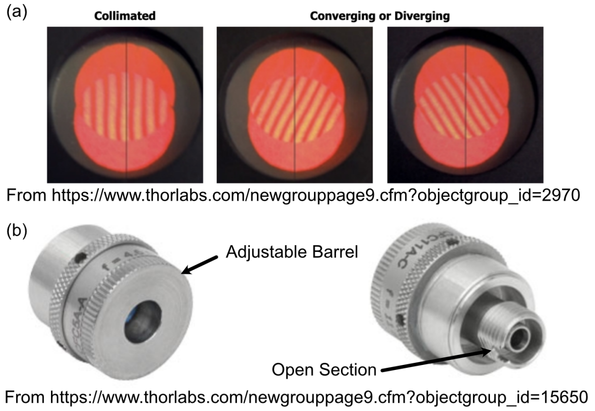

Fig. 4: **Figure 4:** Shear Plate interferometer and collimator lens

using this method, the ID12 to Polaris Adapters can just be directly mounted onto the holes designated for L1 and the Illumination Objective, depicted below, to cover the length of the baseplate. With the pinholes placed, the process becomes iterative by making small adjustments on the K1XY tip/tilt knobs and XY position screws until the beam passes through both pinholes.

---

##### 6.2.4 Mounting of Optic Posts

After ensuring basic collimation and alignment of the laser system, the next step is mounting the appropriate polaris post size for each optical element in the system. The overall breakdown of which size posts went with each hole location is listed below. Where holes (3), (4), and (10) are placements for optional elements in the system. (3) is meant for the placement of an adjustable pinhole aperture, where the same ID12 to Polaris Adapter system used in the alignment step can be placed. Otherwise a 0.5" post holder can be placed here, and then the ID12 or ones pinhole aperture of choice can be mounted on a 0.5" post. (4) is dedicated placement for an electronic shutter in the event that the laser being used for the system isn't directly controlled via a computer, and helps to ensure that laser light is not photobleaching a sample unless imaging is taking place. (10) is dedicated space for a neutral density (ND) filter, in the event that the power of the laser used needs additional reduction not directly addressable through the laser controls itself.

---

##### 6.2.5 Mounting of Optics

With the posts in place, the next step involves mounting the optics elements themselves onto the posts. When mounting elements (6), the VA100 rectangular aperture, and (7), the RSP1 rotation mount, it should be noted that the corresponding element should first be fixed onto their respective Polaris adapter and then onto the Polaris posts. In addition, take measures to mount the 45 degree mirror (9) fully prior to mounting the resonant galvo to ensure that the galvo is not accidentally hit during installation of the 45 degree mirror.

To construct the 45 degree mirror assembly, first mount the Polaris K1S4 onto the Polaris-MA45 45 degree mounting adapter, and then place a 1" diameter mirror into the K1S4 and secure it using the K1S4 mounting screw.

To construct the illumination objective assembly, first mount the Polaris 1XY onto its respective 1.5" post. The threading of the 1XY (SM1) does not match that of our illumination objective used (M25x0.75), so an SM1A12 adapter must first be installed onto either the 1XY or the illumination objective. Then, the objective can be mounted onto the 1XY.

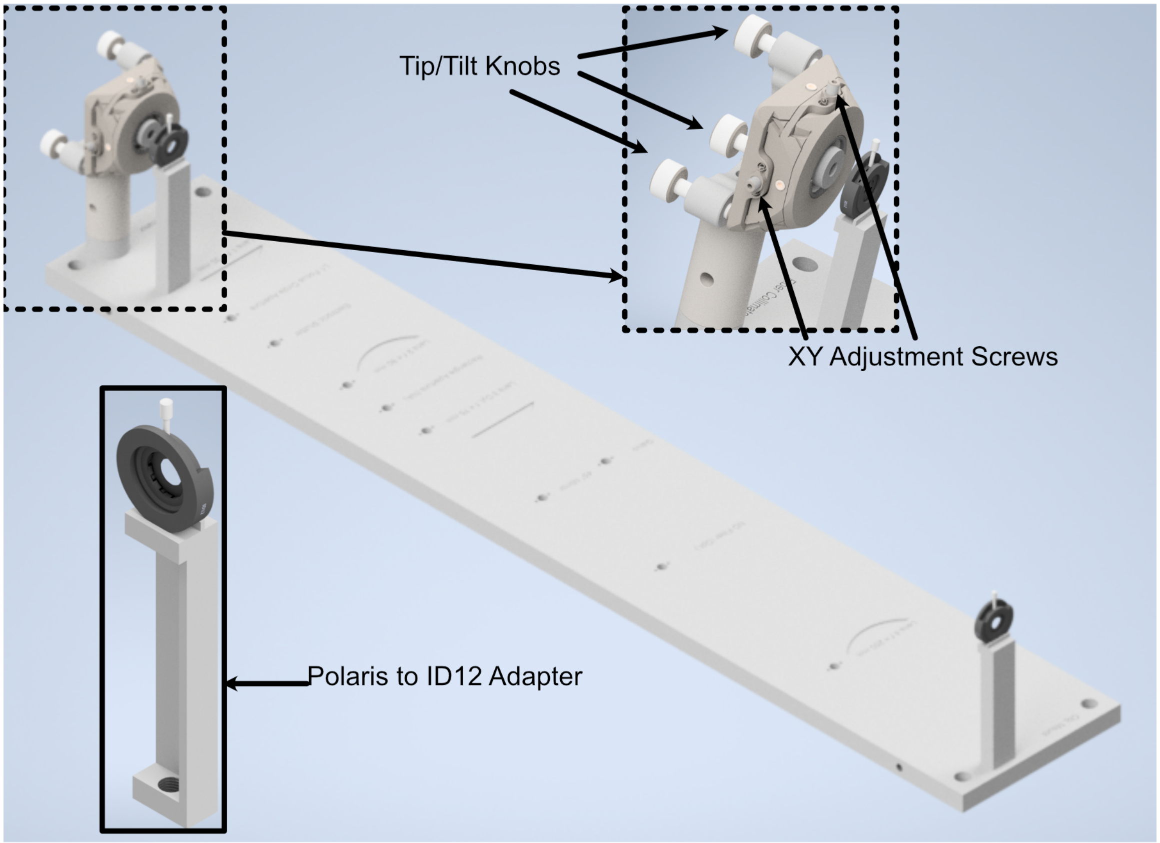

Fig. 5: **Figure 5:** Performing beam alignment across the baseplate

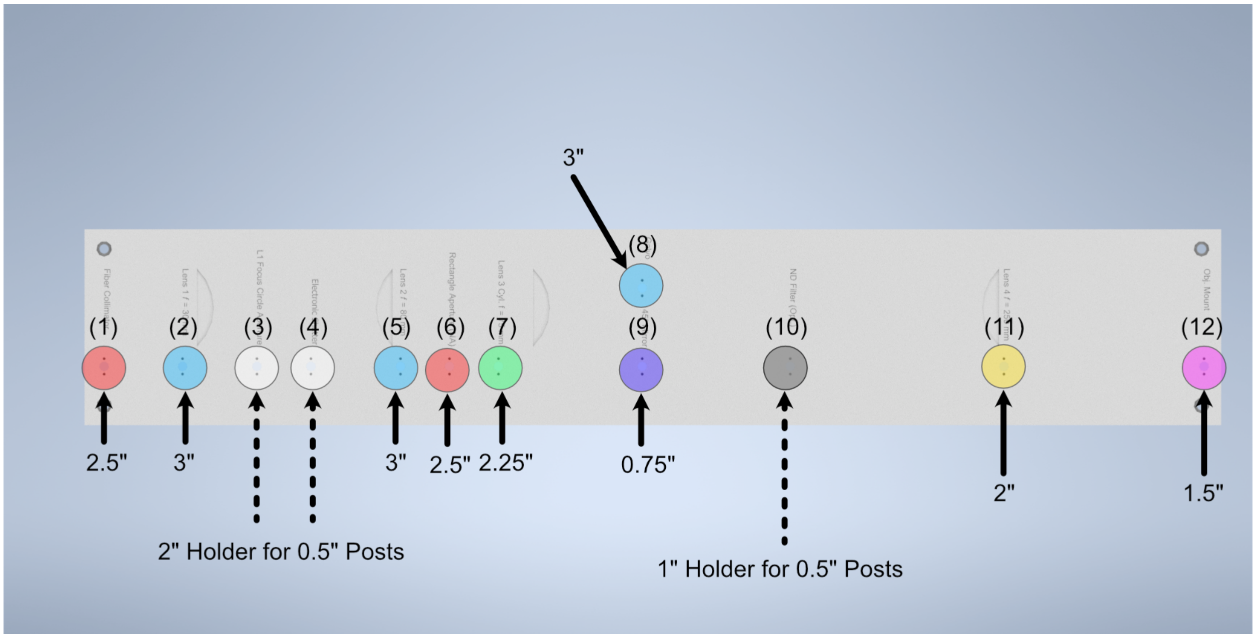

Fig. 6: **Figure 6:** Schematic of which holes use which post heights

##### 6.2.6 Final Alignment Adjustments

Once all the optics have been properly mounted, the final step for ensuring a working illumination path is to center the beam after the 45 degree mirror onto the center of back lens surface of the illumination objective. The primary method of doing this is through adjusting the xy tip/tilt knobs on the K1S4 Polaris mount that houses the 45 degree mirror.

#### 6.3 Detection Path

##### 6.3.1 Detection Path Assembly

Our detection path consists of our Nikon N25X-APO-MP detection objective, Hamamatsu ORCA Flash4.0 V3 Camera, ASI C60-TUBE-400 tube lens, and ASI FW-0002-8 8-position filter wheel unit. These components are mounted together and affixed onto an ASI LS-100-AMCCH translation stage for focus control. We currently use a prototype detection path baseplate (available here) as a mounting stage for these elements and the sample chamber; however, this additional baseplate is still undergoing design iterations and is not critical for a functional detection path.

We utilize two additional custom adapter elements in the construction of the detection path: a shell casing around the tube lens that mounts to the translation stage and an adapter for the translation stage to mount it to an optical breadboard table. The height thicknesses of these elements were chosen such that the height of the detection objective center should match that of the illumination objective (which with the 1.5" tall posts our illumination baseplate rests on is 4.75" above the optical table surface). These elements can be custom machined if desired; however, we have found 3D printed PLA variants to perform their functions effectively as well.

The assembly of the detection path begins with the translation stage and it's associated breadboard adapter (available here in two variants, based on whether or not one is using the 0.5" thick detection path baseplate):

1. Turn the translation stage upside-down
2. Place the breadboard adapter upside-down on top of the inverted translation stage (such that the raised platform of the adapter is touching the bottom side of the translation stage)
3. Align the recesses on the bottom of the adapter with the holes on the bottom of the translation stage
4. Fix the adapter onto the translation stage by screwing M6 screws into the recesses aligned with the translation stage holes.

The next step is flipping the translation stage assembly right side up again, and then fixing the first of two halves of the tube lens adapter onto the top of the translation stage:

1. Place the tube lens adapter half onto the top of the translation stage such that the block with two sets of five recessed holes is touching the top of the translation stage.
2. Align the Recess holes on the adapter with the holes on the top of the translation stage.
3. Fix the adapter onto the translation stage by screwing M6 screws into the aligned recess holes

Next, we'll focus on assembling the tube lens and filter wheel:

1. Take the MIM to Filter wheel adapter and fix it onto the front port of the filter wheel using the associated screw ports
2. With the adapter fixed, now screw the 400 mm tube lens into the adapter.

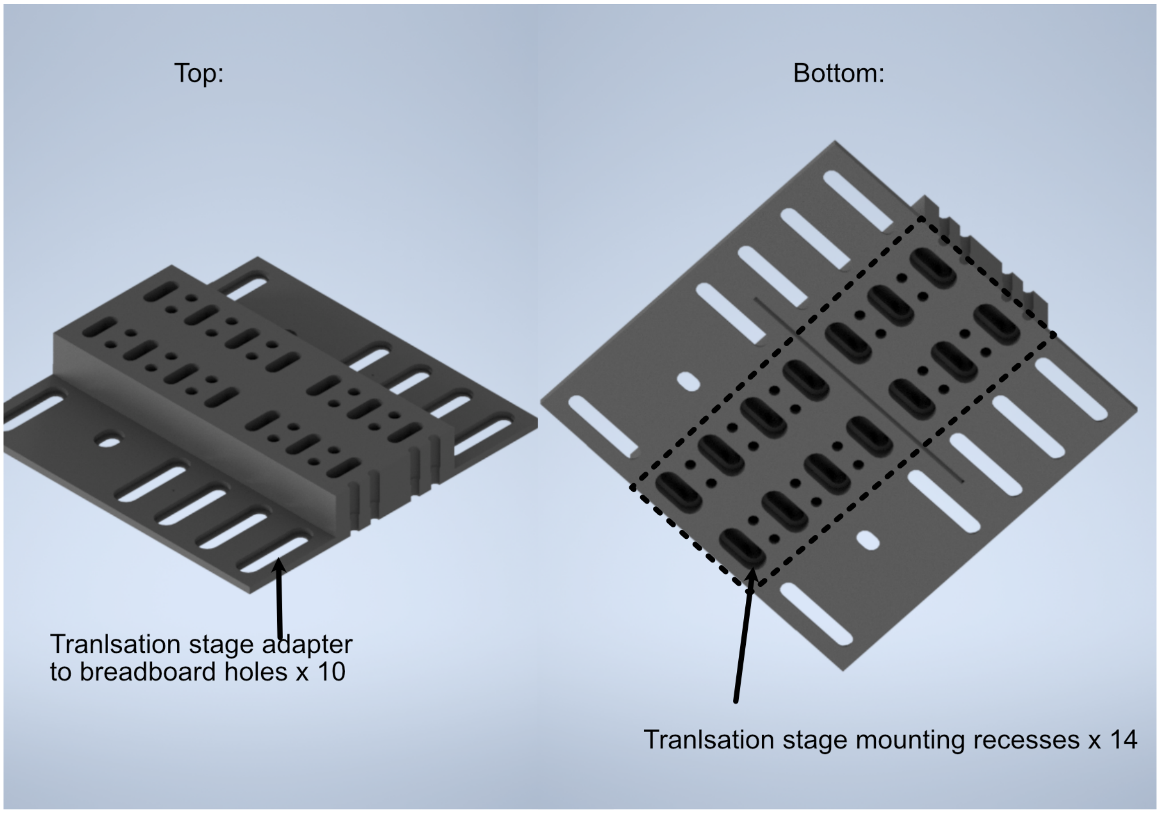

Fig. 7: **Figure 7:** Schematic of the translation stage breadboard adapter

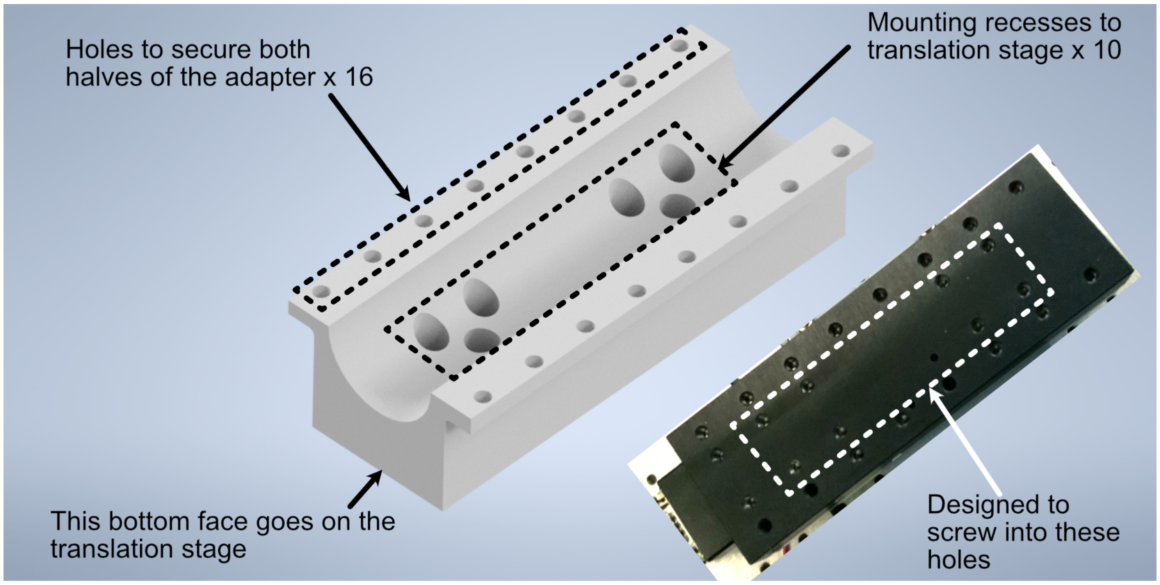

Fig. 8: **Figure 8:** Schematic of the tube lens to translation stage adapter

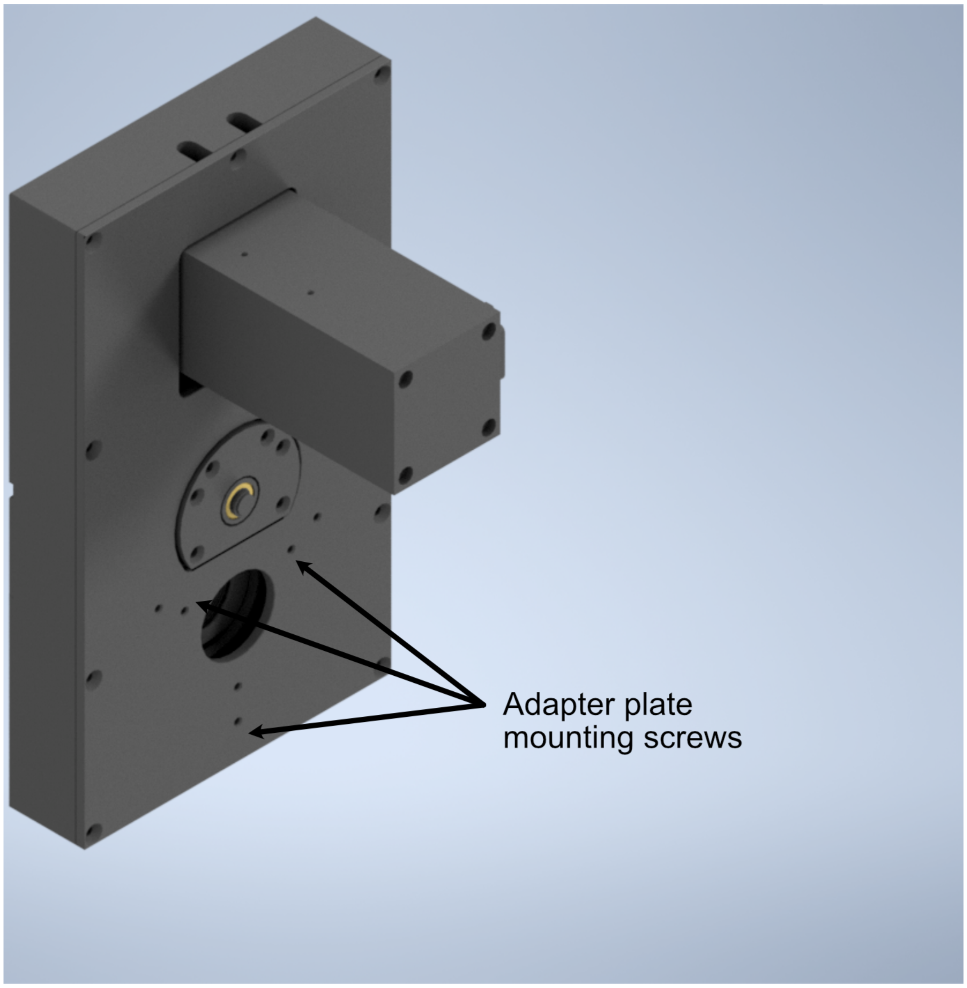

Fig. 9: **Figure 9:** Schematic of the filter wheel port for the tube lens

**In order to fix our detection objective onto the tube lens, we must first prepare an extension and threading adapter:**

1. Take the C60-EXT-15 15 mm Tube extension piece and place the RAO-0051 M32x0.75 threaded sleeve inside
2. Using the screws on the top of the extension piece, fix the threaded sleeve in place
3. Insert/screw the extension piece into the front of the tube lens.

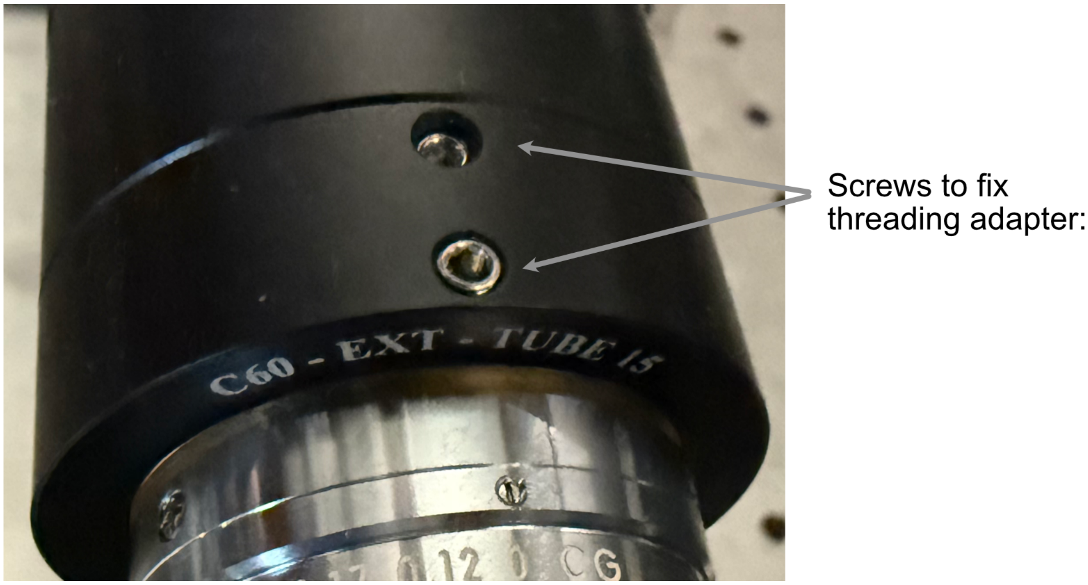

Fig. 10: **Figure 10:** Showcase of the screws used to secure the thread adapter for the tube lens

**The tube lens assembly is now ready to be fixed onto the translation stage assembly:**

1. Place the tube lens assembly such that the tube lens lies within the curved region of the tube lens adapter
2. While there isn't an exact science to the relative placement of the tube within the adapter, try to position it such that more of the tube is extended out on the side where the objective will be mounted (our setup is shown below for reference).
3. Place the second half of the tube lens adapter such that the curved side fits onto the lube lens and position it such that the holes of both halves of the adapter align with each other.
4. Using your choice of either M6 or 1/4"-20 screws and associated washers/nuts, place the screws with a washer placed on them first into the aligned holes of the adapter. We used 4 of the adapter holes on each side, but more can be used for extra security if desired.
5. Screw a washer onto each of the screws until they're secured against the bottom lip of the adapter.

The detection path assembly can now be fixed into place onto either the detection path baseplate or the optical table, keep in mind this process is meant to essentially place the unit in the ballpark of where it should be, finer adjustments will be made afterwards:

1. Using the mounting holes on the translation stage assembly, place the assembly such that the edge of the translation stage adapter facing the illumination path is roughly 9-10" away from the location of the illumination objective.
2. Using the adjacent edge of the translation stage adapter (the one that should be perpendicular to the orientation of the illumination path), try to align the side of the adapter with the mounting hole of the illumination objective.

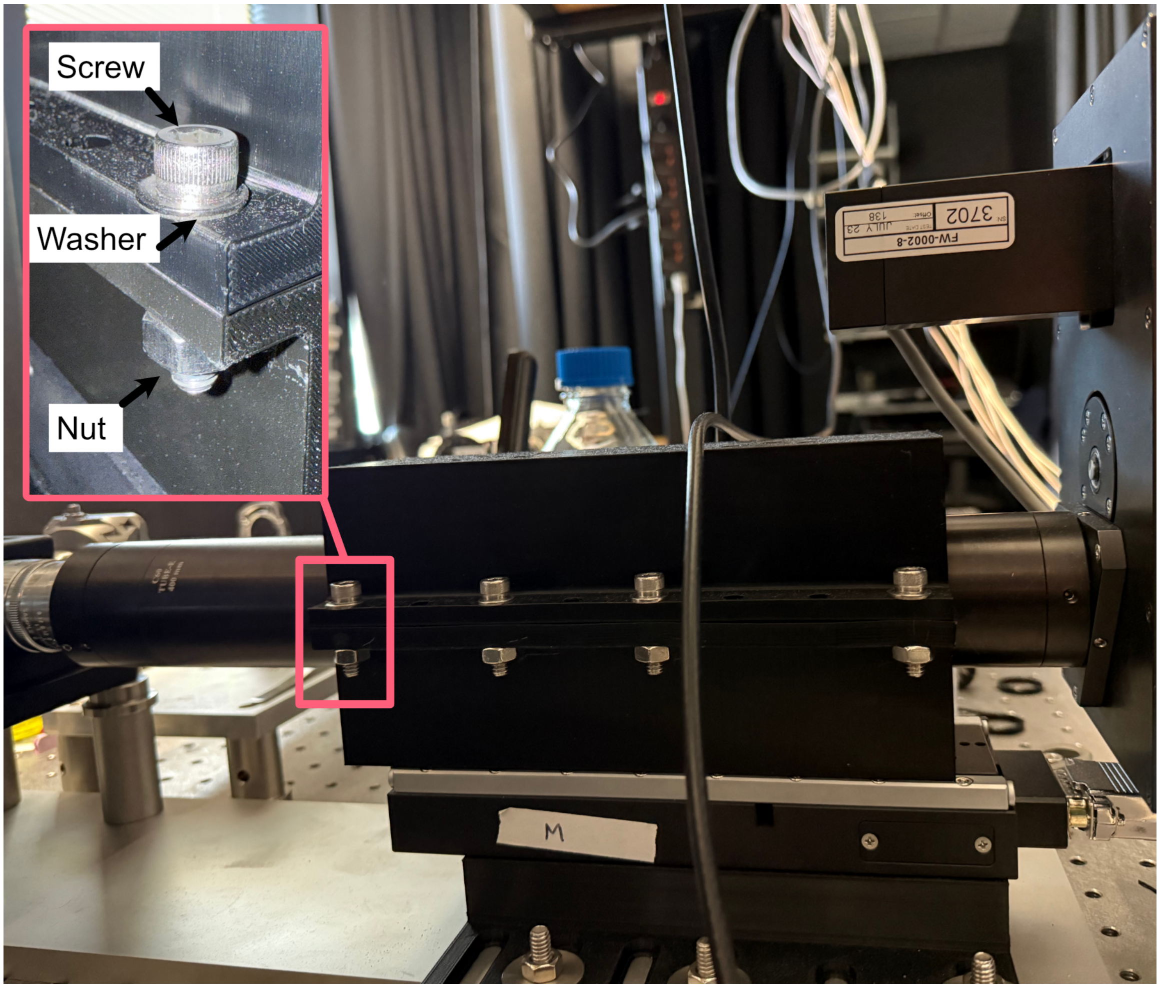

Fig. 11: **Figure 11:** Example of tube lens mounted in the corresponding adapter

3. Screw the translation stage adapter into either the optical table to the detection path baseplate (we recommend using Thorlabs 1” Spacers in place of washers here).

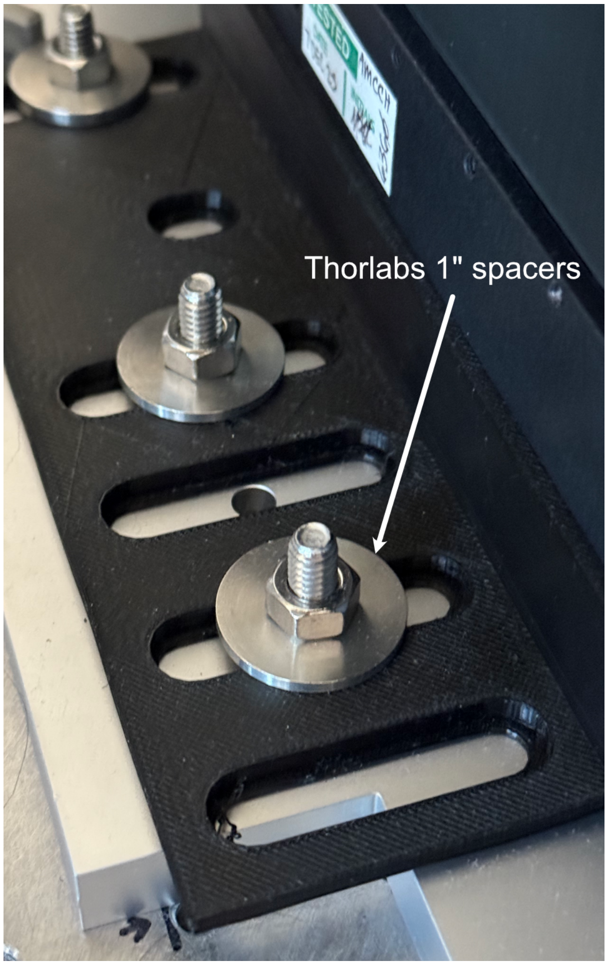

Fig. 12: **Figure 12:** Setup for securing the translation stage to breadboard adapter onto our detection path baseplate

**With the assembly fixed in place, the camera can then be screwed into the filter wheel:**

1. Align the front thread of the camera with the back port of the filter wheel
2. Screw the camera into the filter wheel until there is resistance
3. Slowly adjust the camera tilt until the top surface is leveled (we use a bubble leveling tool for this, shown below)

The final steps to to assemble the detection path are to screw the detection objective into the front of the tube lens and attach all associated wires to the camera, filter wheel, and translation stage. If there’s not enough clearance between the front of the tube lens and the sample chamber to screw in the detection objective, the translation stage might need to be wired up first and translated backwards manually using either navigate or the Tiger Control Panel software.

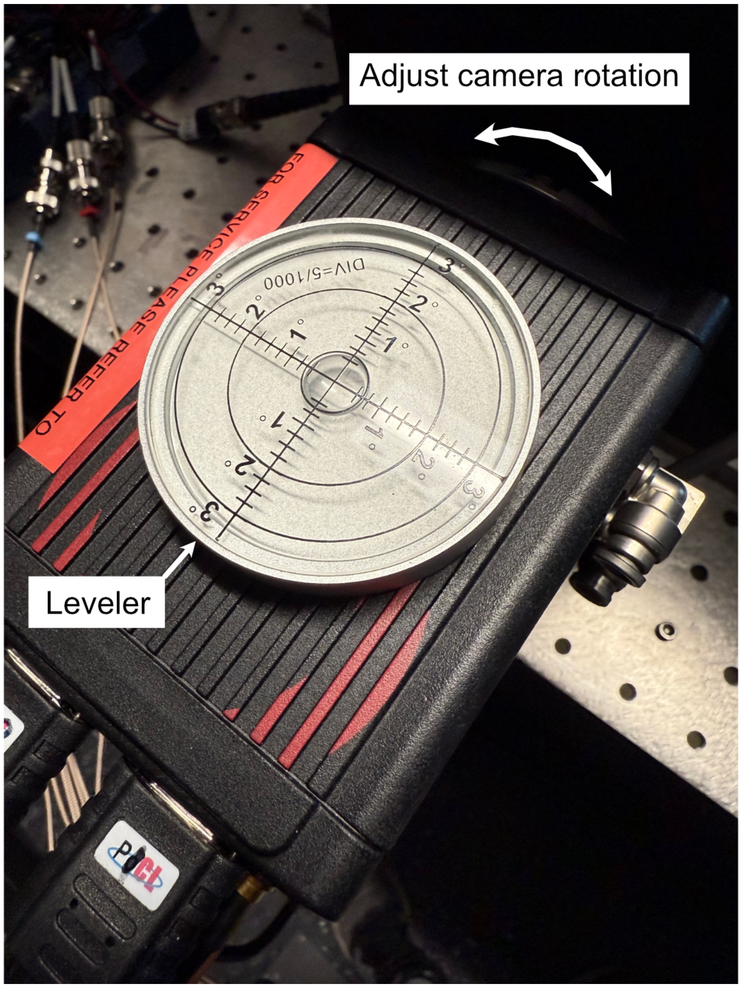

Fig. 13: **Figure 13:** Mounting of the camera

##### 6.3.2 Detection Path Positioning

To position the detection path correctly, there's two primary steps: Initially ensuring the detection objective slides into the correct port on the fixed sample chamber and then ensuring that the light sheet is centered in the FoV of the camera, typically done by putting fluorescein into the water-filled chamber.

**Step 1: Ensuring the detection objective slides into the sample chamber port**

1. It's recommended to start with your detection path translation stage set far enough back such that there is anywhere from 0.5-1" or more of space between the front of the detection objective and the sample chamber.
2. Within navigate or the Tiger Controller software, start to slowly move the objective forward (initially increments of 1000 um should be okay, but as the objective approaches the port the distance should be significantly lowered to 100 um or below to ensure the objective isn't damaged).
3. If the objective isn't centered properly, the screws/nuts on the translation stage to breadboard adapter (see figure 12) can be loosened slightly and the detection path assembly can be carefully positioned horizontally, taking care that the unit isn't tilted at an angle going into the objective port, and then re-tightened.

Once the detection objective properly translates into it's port, the tilt alignment can be further refined by slowly translating the detection objective forward towards the illumination objective, taking great care that they don't touch each other. If there is a significant tilt in the detection path compared to the illumination path, it should be fairly evident in the diagonal space between the objectives (ideally it should look roughly equal in width between the two objectives). As example of the relative position of our objectives when the detection objective is properly placed at the focus of the illumination objective is shown below to give a visual reference for users.

Once the position of the detection objective is acceptable, further refinement of it's position can be done by first filling the chamber with water and then adding roughly ~100 microliters of f fluorescein into the chamber. Fluorescein is useful as a visualization tool to be able to see the beam coming out of the illumination objective with the naked eye. With fluorescein in the chamber, the position of the detection objective can be translated to align with the focus of the beam itself (shown below)

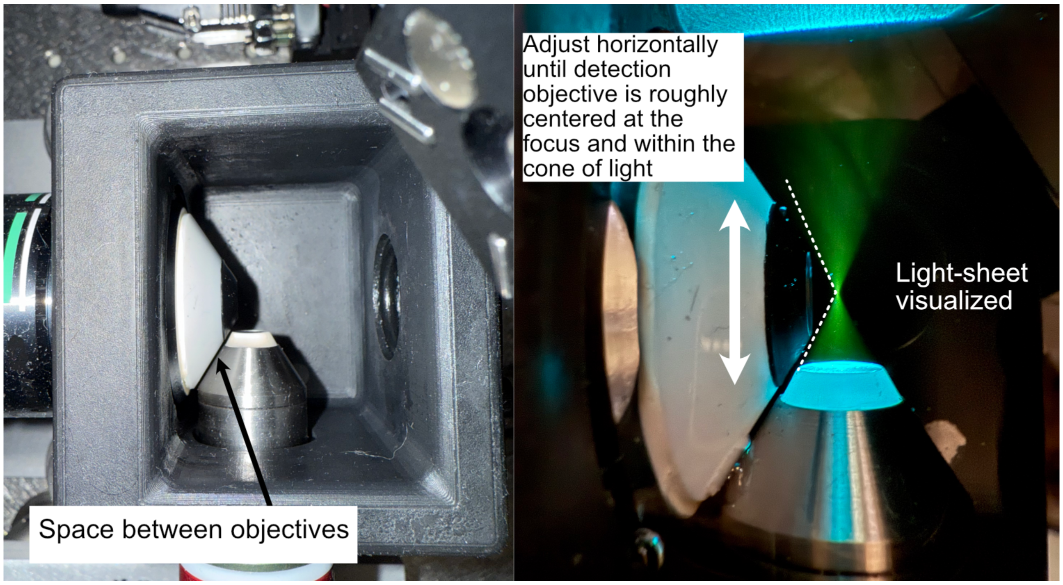

Fig. 14: **Figure 14:** Physical alignment of the detection objective

**Step 2: Focus refinement**

With the detection objective roughly in the correct location, further refinement is done by turning on the camera and imaging the fluorescein light-sheet to find the focus by adjusting the focus translation stage in small increments (50 microns for large movements, 1-5 microns for small movements). The goals during this step are to ensure that the detection objective focus is at the focus of the illumination objective, to make sure the light-sheet focus is centered on the image FoV from the camera, and that the light-sheet is aligned vertically in the image FoV.

**Finding the focus is often something that just comes with experience of using the system itself; however, here are some general methods to follow that can help get one's bearings:**

1. In general, as the detection objective approaches the light sheet focus, an envelope of light should start to appear (see below), ideally this envelope will be horizontal across the FoV. If it horizontal or has some tilt to it, adjusting the tip/tilt of the 45 degree mirror and the rotation of the cylindrical lens. In addition, ideally this envelope will be centered vertically in the FoV, which can be set by adjusting the y adjustment screw on the illumination objective mount.
2. Towards the focus location, a dark vertical region with horizontal striations should appear. Once this region is visible, using small translation distances (~1 um or finer) to find the position that makes both the vertical region and horizontal striations look as sharp as possible should be done.
3. If the dark region isn't centered on the image FoV, adjusting the x and y adjustment screws on the illumination objective mount and making slight horizontal translations on the detection path can help.
4. In general doing these adjustments in small increments is helpful, as once adjustments to the various elements may require readjusting the focus translation stage position to keep track of the focus.

#### 6.4 Optomechanics

##### 6.4.1 Visualization of Axes Mapping

In our system we have 5 different translation stages at work: the standard x,y, and z axes, an additional stage along the z axis to control the focus of the detection path (f), and and axis associated with the piezo positioned such that its normal is 60.5 degrees away from the y-axis.

**Note: Coordinate System Differences**

The coordinate axes used in **optical simulations** differ from those in the **physical microscope system** controlled by **navigate**. This discrepancy arises from **standardized axis conventions** in Zemax versus those historically used in our previous systems. Users should be mindful of this distinction when interpreting simulation results versus real-world microscope operation. The differences are illustrated below:

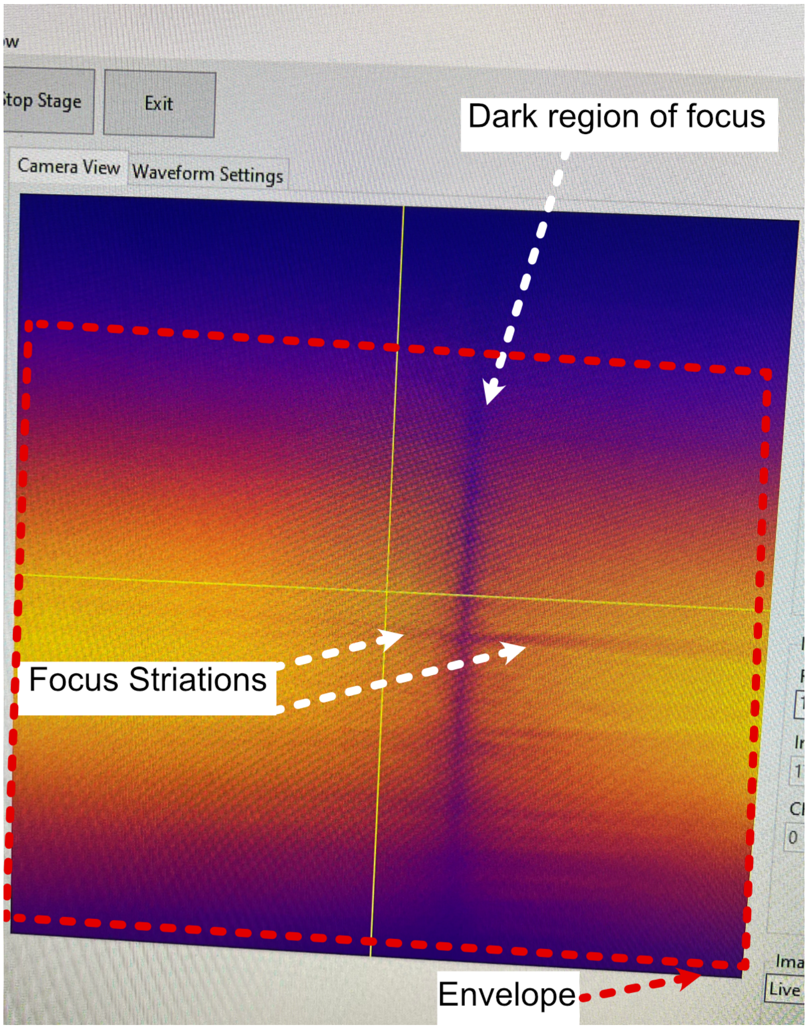

Fig. 15: **Figure 15:** Breakdown of the light sheet focus within navigate

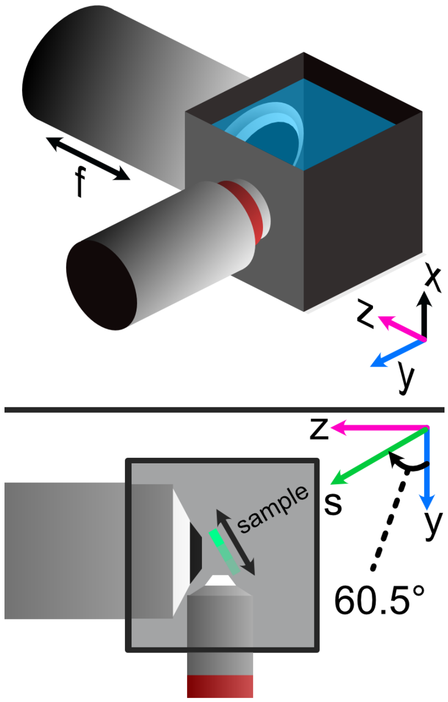

Fig. 16: **Figure 16:** Layout of how the axis of the system are mapped

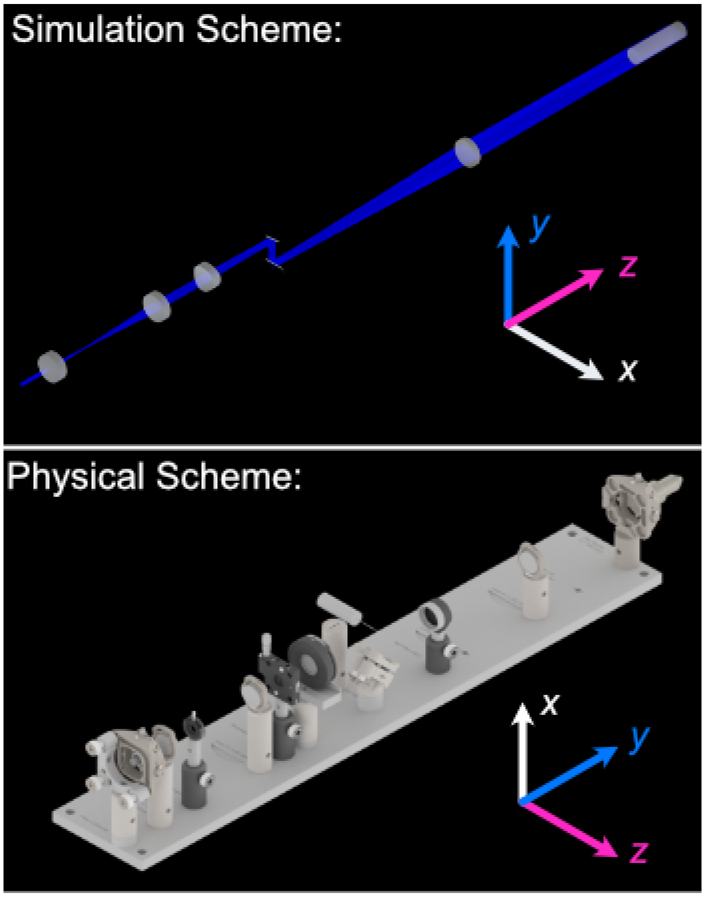

##### 6.4.2 The Piezo Angle Mount

We designed a custom angled mount for our Piezoconcept HS1 piezo in order to be able to scan our sample easily between our two objectives by translating a single motorized unit (in this case the piezo), instead of having to calculate and program the movement of two translation stages in tandem for both the y and z directions. The anatomy of our angled mount is broken down in the figure below, where there are four translation stage mounting holes to attach the unit to an ASI translation stage, nine Piezo mounting holes that correspond to the mounting scheme of our piezo unit, as well as four through-holes and a window for ease of access for the mounting process. We provide the CAD files for this mount here, and have had success in using both 3D printed and aluminum machined versions of the unit. It's recommended to first mount the angle mount onto the translation stage unit before mounting the piezo on the angle mount to ensure access to all the through-holes.

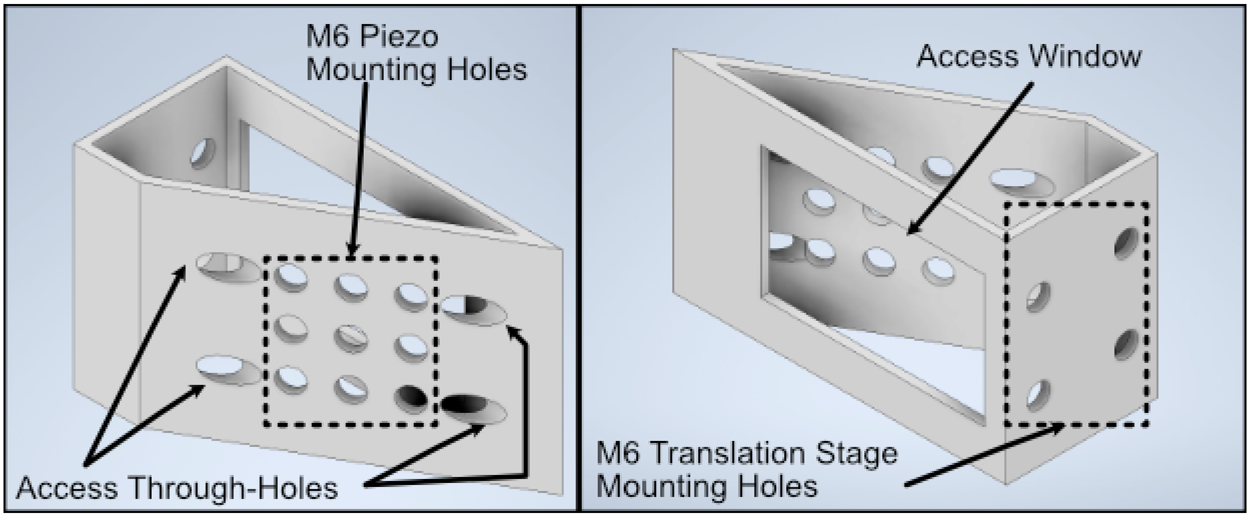

Fig. 17: **Figure 17:** Breakdown of our custom angle piezo angle mount.

The installation of our custom angled piezo mount is designed to be directly compatible with ASI translation stages. ASI translation stages feature M6 hole pairs that are spaced along the length of the translation stage at intervals dependent on the specific stage one is using. The mounting process involves aligning these 4 holes with 4 of the M6 holes on the translation stage and screwing them in. For ease of screwing in the base, there are four through holes on the angled face of the mount shown in B that a screwdriver is able to pass directly through to screw as shown in C. An alternative method of mounting is shown in D, where the window on the back of the angle mount is able to be screwed through as well.

##### 6.4.3 Sample Holder Design

Our sample holder design is built for imaging cells on a 5 mm coverslip, and is shown below. The design features a clamp-like method of securing the 5 mm coverslip in place, where the coverslip rests in an inset region and the clamp is screwed in via an M1.6 screw in the back of the holder. All associated files for this design and other custom parts can be found here.

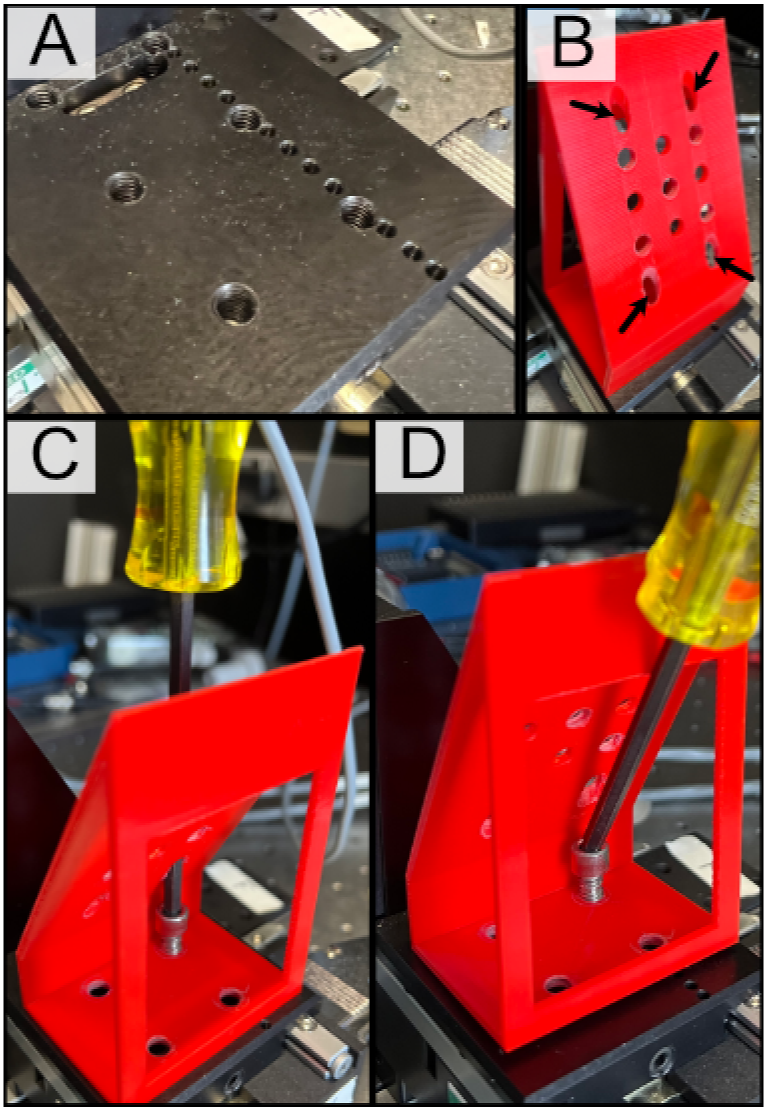

Fig. 18: **Figure 18:** General process for mounting our piezo angle mount onto an ASI translation stage

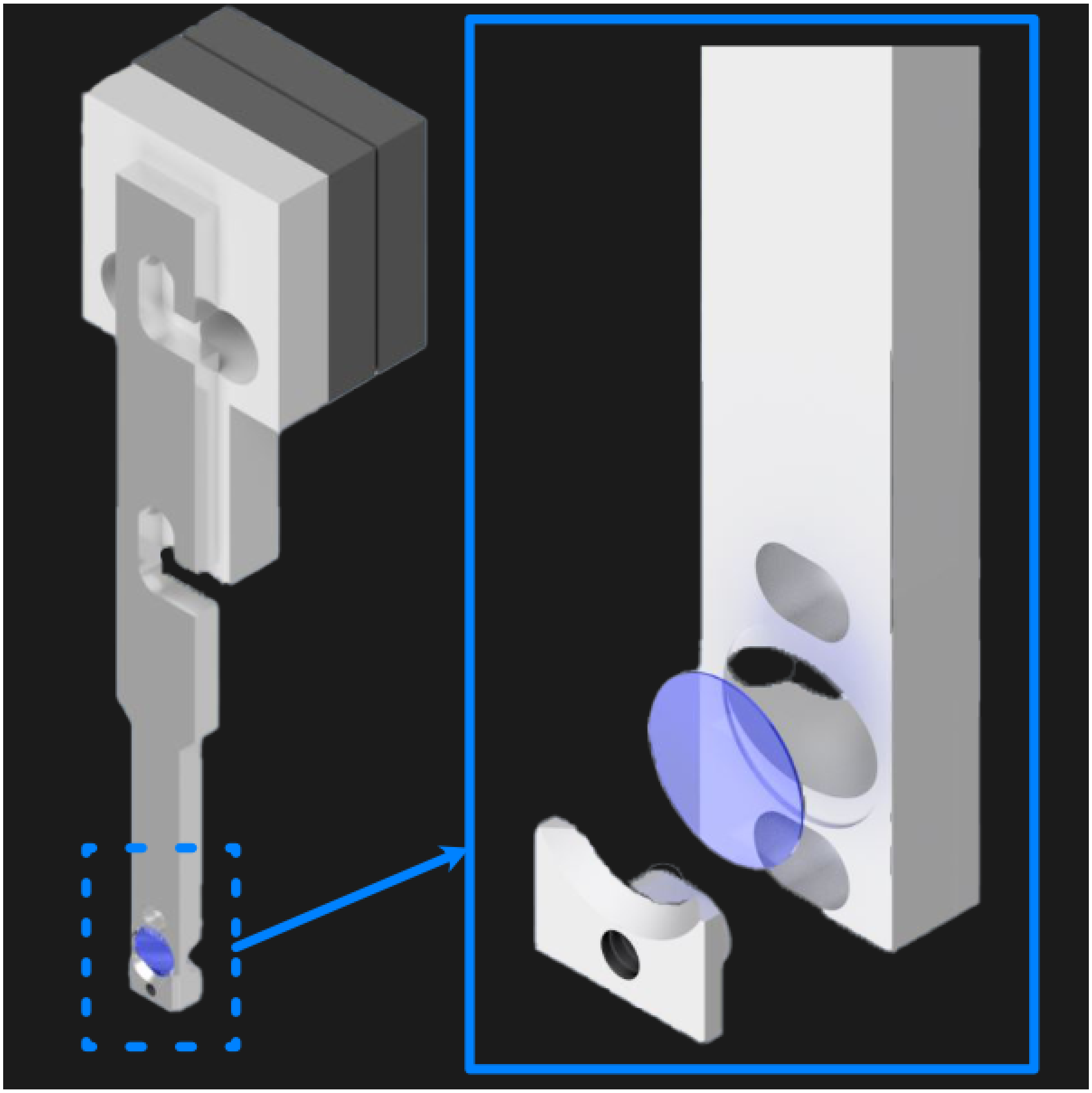

Fig. 19: **Figure 19:** 5 mm coverslip sample holder design.

##### 6.4.4 Assembling the Magnetic Sample Mount

As a safeguard for the risk of the sample crashing into either the illumination or detection objective during sample positioning or imaging, we opted to incorporate a magnetic mount for our sample holder. We use a Thorlabs KBT1X1T and KBB1X1 as our magnetic mount pair, and then mount our sample holder onto the KBT1X1T using a custom adapter. The KBB1X1 is affixed to the piezo using an M2.5 screw, and using a small leveling tool (shown below) during this step is helpful to ensure that the magnetic base is mounted as level as possible for the imaging process.

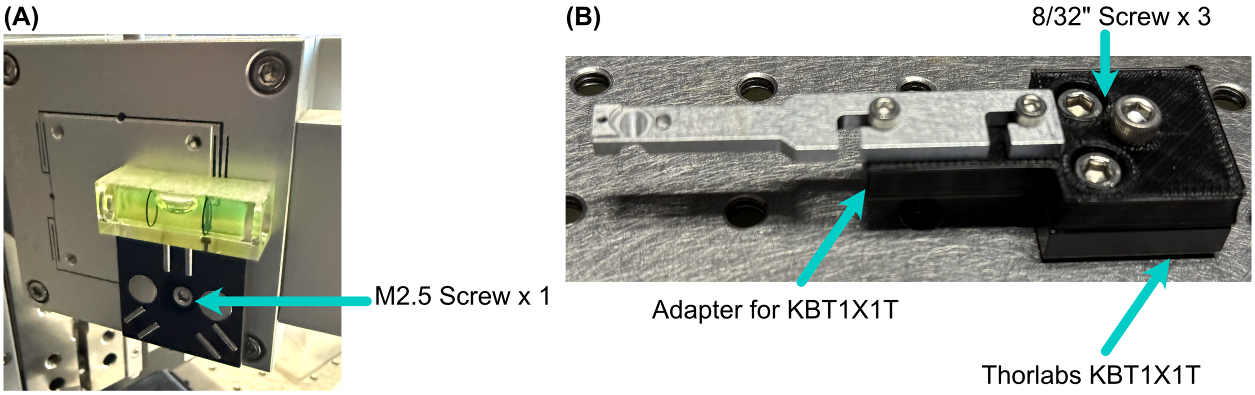

Fig. 20: **Figure 20:** Basic assembly of magnetic sample holder mount

---

#### 6.5 Optoelectronics

##### 6.5.1 Wiring Diagram

Our complete microscope assembly features a variety of different optoelectrical and optomechanical elements. These elements are primary controlled via our NI DAQ (PXIe-6738) or our ASI Tiger Controller (TG16-BASIC), which are then controlled via navigate during the imaging process. The diagram below depicts how these elements are wired together, as well as an individual pinout designation table for the pin configurations we used on our DAQ.

---

##### 6.5.2 Piezo Setup & Troubleshooting

On the PCI Board, connect the positive and negative wires to the corresponding analog output (AO) you want, in our case we used AO 0, so we connected the positive wire to pin 10 and the ground to pin 11, then plug the BNC cable connected to those wires into the EXT IN input on the Tiger controller panel corresponding to the piezo.

Plug the piezo cable into the PIEZO input on the Tiger controller panel corresponding to the piezo, and verify the range of the piezo in the Tiger Controller software:

Listing 1: Verifying Piezo Range in Tiger Controller Software

```
# Verify the range of the piezo in the Tiger controller software
5 cca x?

# Example output:
```

(continues on next page)

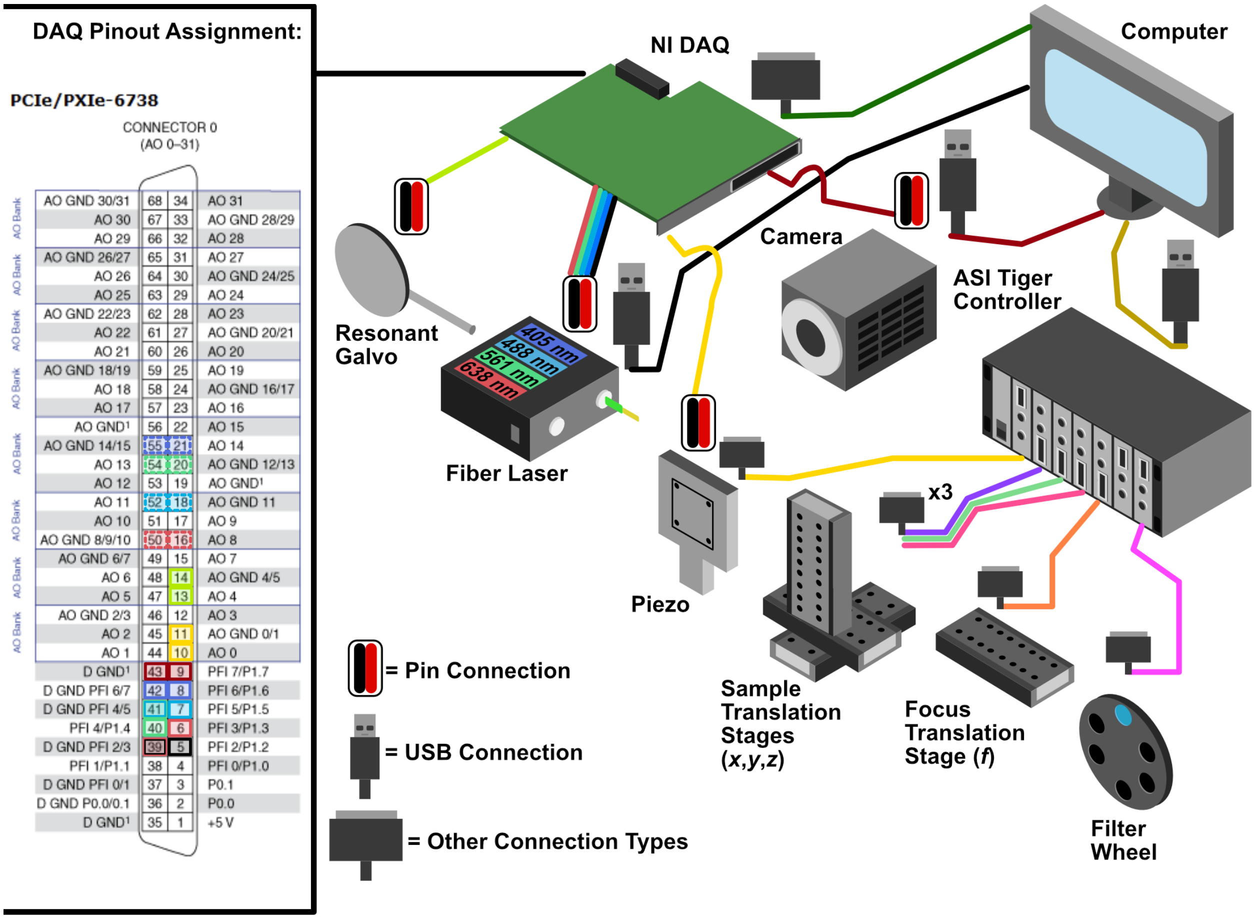

| Component | Analog/Digital? | Ground Pin | Active Pin |
|---|---|---|---|
| Resonant Galvo | Analog | 14 | 13 |
| Piezo Motor | Analog | 11 | 10 |
| Camera | Digital | 43 | 9 |
| Fiber Laser (FL) Output Shutter | Digital | 39 | 5 |
| FL λ = 405 nm AO Port | Analog | 55 | 21 |
| FL λ = 405 nm IO Port | Digital | 42 | 8 |
| FL λ = 488 nm AO Port | Analog | 18 | 52 |
| FL λ = 488 nm IO Port | Digital | 41 | 7 |
| FL λ = 561 nm AO Port | Analog | 20 | 54 |
| FL λ = 561 nm IO Port | Digital | 41 | 40 |
| FL λ = 638 nm AO Port | Analog | 50 | 16 |
| FL λ = 638 nm IO Port | Digital | 39 | 6 |

Fig. 21: **Figure 21:** Wiring diagram for the Tiger Controller, DAQ, and associated elements.

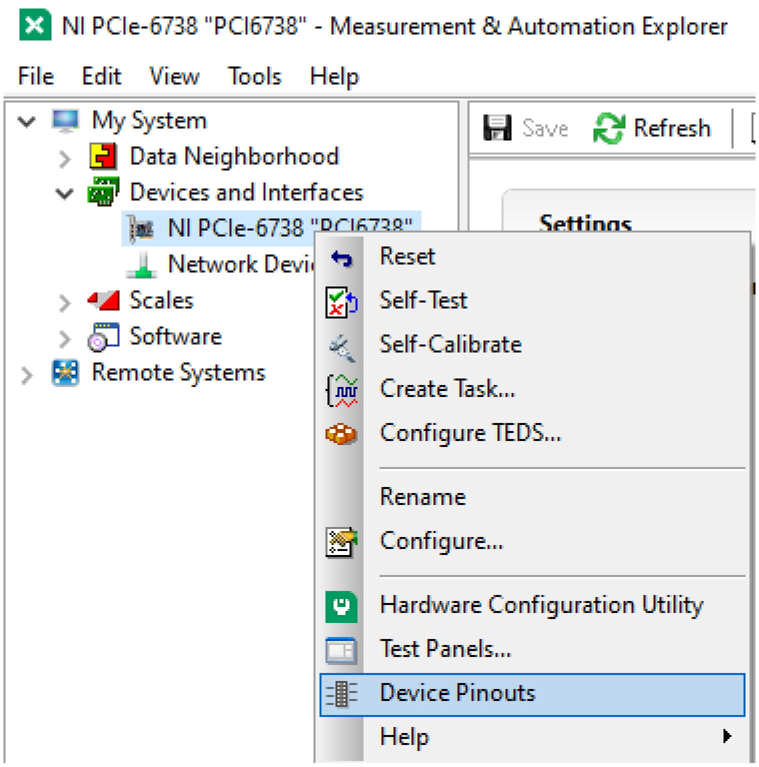

Fig. 22: **Figure 22:** How to find the Device Pinout panel in NI MAXX.

(continued from previous page)

```
:A  Q:P1
23 P 1 100um RANGE
24 P 2 200um RANGE
35 P S 150um RANGE
36 P 3 300um RANGE
37 P 5 500um RANGE
34 P f 50um RANGE
25 P 4 350um RANGE:N-4
```

This tells us that our Piezo (Panel 5/Q) corresponded to P1 or a 100 um range, but ASI requires the piezo needed to be set to a 50 um range to be able to be initialized instead. To change this, we used the command `5 cca x = 34` and power cycled the controller. Once the controller was powered back on, we verified the range again with the `5 cca x?`

Listing 2: Verifying Piezo Range in Tiger Controller Software

```
# Verify the range of the piezo in the Tiger controller software
5 cca x?

# Example output:
:A  Q:Pf
...
```

Now we can see that the piezo is set to the correct range (Pf). With that verified, now confirm that the voltage output from the PCI Board is working:

1. Put the BNC cable input currently in EXT IN on the Tiger control panel into the input of the oscilloscope instead.
2. Go to the test panels for the PCI board in NI MAX.

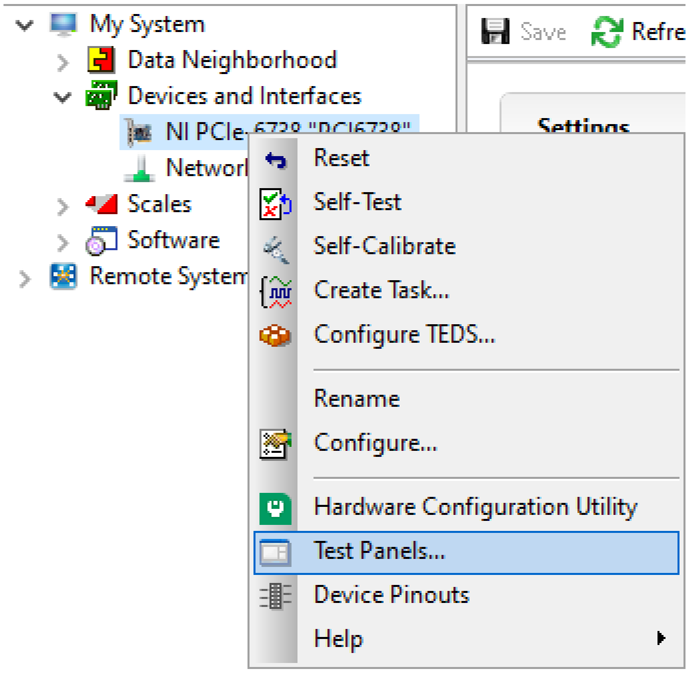

Fig. 23: **Figure 23:** How to find the Test Panels panel in NI MAXX.

3. Set the voltage mode to sinewave generation.
4. Set the voltage range to be between 0 to 10 V.
5. Set the frequency to a desired value (we ended up setting it pretty high at 10000 Hz for ease of viewing on the oscilloscope).

With the voltage output of the PCI board verified, plug the PCI Board voltage cable output back into the EXT IN slot and verify that the position output of the Piezo reads similarly on the oscilloscope:

1. Plug a BNC Cable into the SENSOR OUT connection on the tiger controller panel.

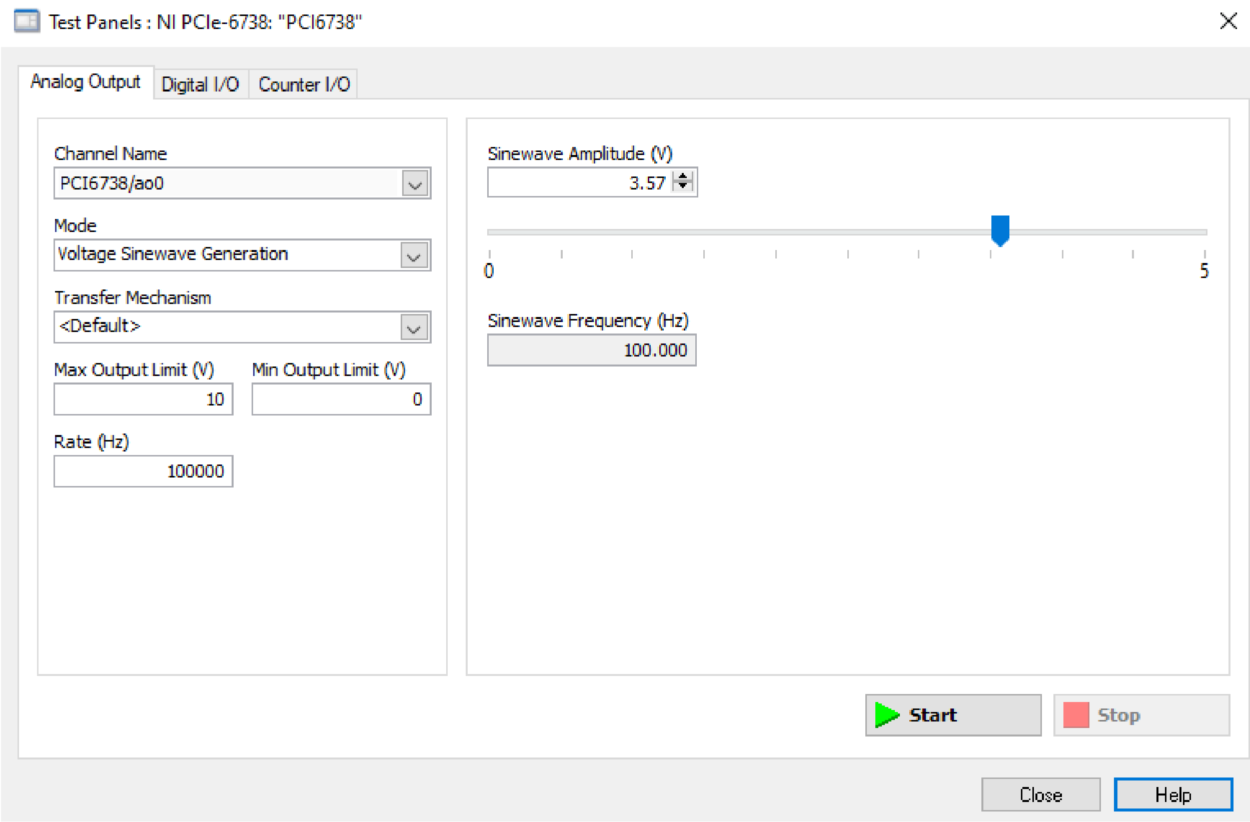

Fig. 24: **Figure 24:** How to generate analog output voltages with NI MAXX for testing purposes.

2. Plug the other end of that cable into the oscilloscope.
3. Verify that a sinewave output is seen on the oscilloscope.

If the PCI Board voltage is working as intended but the piezo position output doesn't seem to work, try ensuring that the piezo is set in External Input mode, and not Controller Input mode:

1. **Use the `PM Q?` (Our piezo corresponds to Q) command:**
   - the output was `Q = 0` originally, telling us that it's in Controller Input mode
2. **Use the `PM Q = 1` command to set the piezo into External Input mode:**
   - now the output of `PM Q?` is `Q = 1`

Another important step is to ensure that the configuration file associated with navigate is appropriately set up for your piezo. This involves setting the correct axis and voltage-to-distance mapping for the piezo. As an example our configuration file for navigate looks like the following for setting up our piezo:

```
type: GalvoNIStage
serial_number: 001
axes: [z]
axes_mapping: [PCI6738/ao0]
volts_per_micron: 0.1*x
min: 0.0
max: 10.0
distance_threshold: 5
settle_duration_ms: 5
controllername: na
stages: na
refmode: na
port: na
baudrate: 0
```

Fig. 25: **Figure 25:** Navigate configuration file for the piezo.

CHAPTER

SEVEN

### SYSTEM CHARACTERIZATION

#### 7.1 Beam Characterization and PSF Analysis

To characterize our constructed system, we first image the generated light-sheet using the sample chamber transmission configuration shown below, where the illumination objective is placed directly in front of the detection objective. The image of our light sheet is shown in (a) below, where the cross-sectional profile of the light-sheet is shown in (b) and reveals a z-FWHM of ~0.415 um.

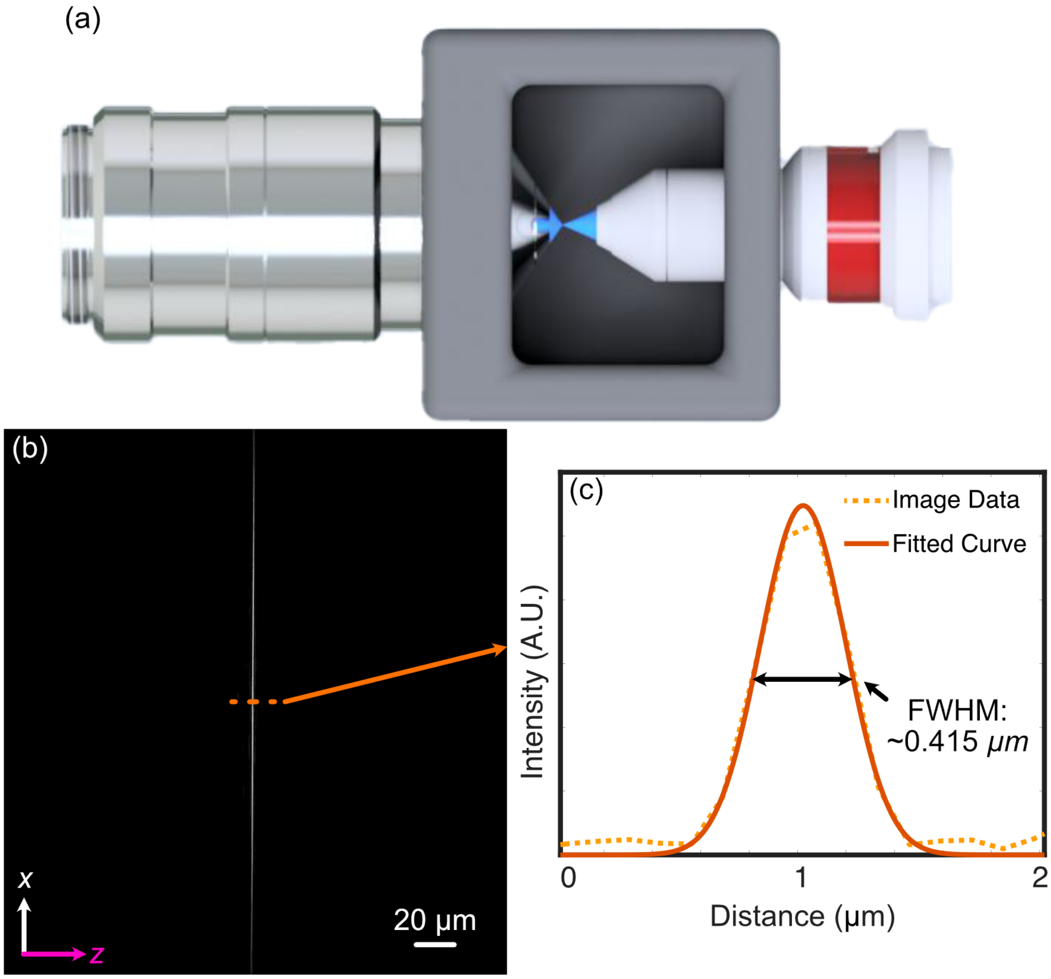

Fig. 1: **Figure 1:** Analysis of the experimental lightsheet characteristics

To characterize the resolution of our system, we utilize 100 nm YG Fluorescent Beads (ID: 17150-10) from Polysciences Inc. The beads are first affixed onto a 5 mm coverslip using the following protocol:

**Note: Affixation protocol for 100 nm beads**

1. Assemble petri dish, coverslip, and 5mM concentration (3-Aminopropyl)triethoxysilane (APTS)
2. Put 5 mm coverslip in petri dish
3. Apply ~100 microliters of (3-Aminopropyl)triethoxysilane (APTS) on top of coverslip
4. Allow APTS to incubate for ~10-30 minutes
5. Wash coverslip lightly with DI water 3 times
6. Put beads of desired dilution (typically 10^-3 or 10^-4 for a normal distribution, 10^-6 for a sparse distribution) onto coverslip and allow to incubate between 2-20 minutes. Longer incubation time allows for more beads to adhere to the coverslip
7. Wash lightly afterwards with DI water

After affixation, the beads are then imaged, the results of which are shown below. The PSF of an isolated bead is shown below in (a-c), where each image is a different orthogonal perspective of the bead's intensity distribution, and provide us insight into the resolution of our system in each orthogonal direction. We then provide Gaussian-fitted distributions of the FWHM of the population of fluorescent beads across a given z-stack in (d), both before and after applying deconvolution procedures. Prior to deconvolution, the average FWHM values across the bead population were 328 in x, 330 nm in y, and 464 nm in z. After deconvolution with PetaKit5D, these values improved to 235.5 nm in x, 233.5 nm in your, and 350.4 in z.

#### 7.2 Sample Biological Images

As a demonstration of Altair's biological imaging capabilities, we prepared and imaged mouse embryonic fibroblast (MEF) cells, where multiple subcellular structures were stained for 4 different channels corresponding to the excitation wavelengths of our laser source (405 nm, 488 nm, 561 nm, 638 nm). The staining protocol described in our initial Altair-LSFM paper was optimized for visualization of the nucleus (DAPI, 405 nm, cyan channel), microtubules (488 nm, gray channel), actin filaments (561 nm, gold channel), and the Golgi apparatus of the MEF cells (638 nm, magenta channel).

A deconvolved maximum intensity projection of a distribution of MEF cells is shown below in (a) and (f), with each individual channel displayed in (b-e) and (g-j). These results display fine nucleolar features within the nucleus, well-defined perinuclear Golgi structures, well-resolved stress fibers in the actin channel, and individually distinct microtubules, demonstrating Altair's ability to capture high-resolution images of cytoskeletal structures.

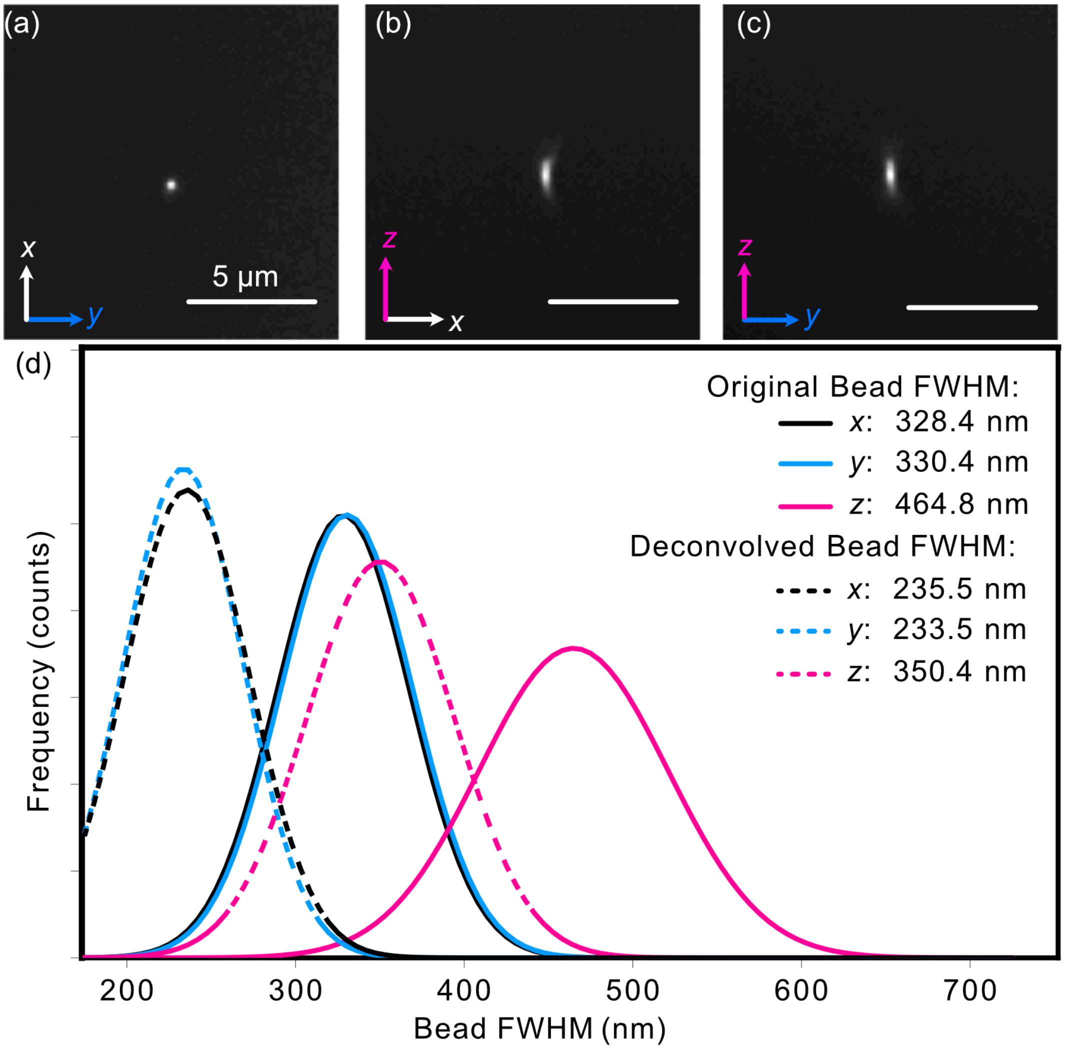

Fig. 2: **Figure 2:** Analysis of the experimental PSF characteristics

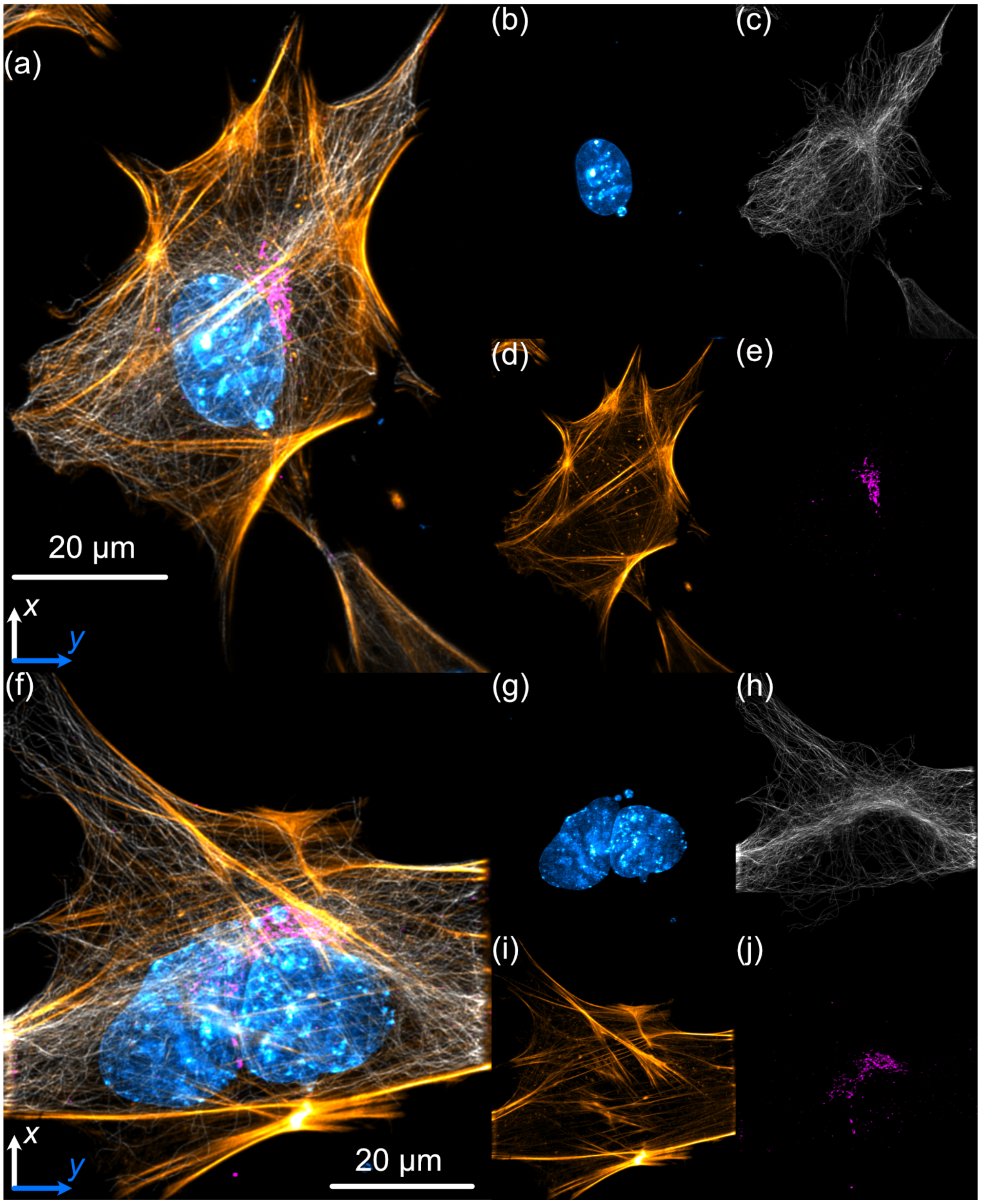

Fig. 3: **Figure 3:** Analysis of MEF Cells

CHAPTER

EIGHT

### IMAGING PROCESS

#### 8.1 Imaging Configurations

Our sample chamber features three ports that provide two distinct imaging configurations shown below: the first is a traditional light-sheet imaging scheme, where illumination and detection objectives are placed orthogonally to each other, and the second is one where the illumination and detection objective are placed coaxially with each other. The first configuration should be thought of as the default imaging setup for the microscope, and the second allows one to observe and characterize the produced light sheet itself. The port not in use should be sealed, which we do using a silicon or rubber seal that's able to be fixed onto the exterior of the port using screws. In addition, it should be noted that our sample chamber design utilizes two sequential layers of O-rings in each of the ports to both secure the objectives and prevent any leaking.

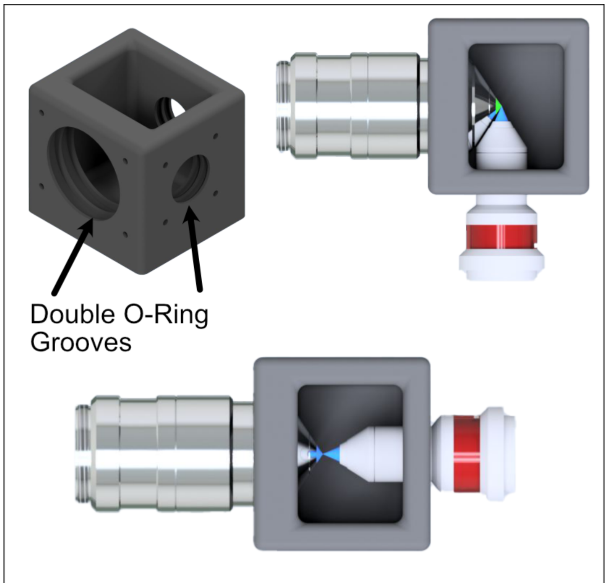

Fig. 1: **Figure 1** Two imaging configurations for the sample chamber design

#### 8.2 Visualization of Axes Mapping

In our system we essentially have 5 different translation stages at work: the standard x,y, and z axes, an additional stage along the z axis to control the focus of the detection path (f), and and axis associated with the piezo positioned such that its normal is 60.5 degrees away from the y-axis.

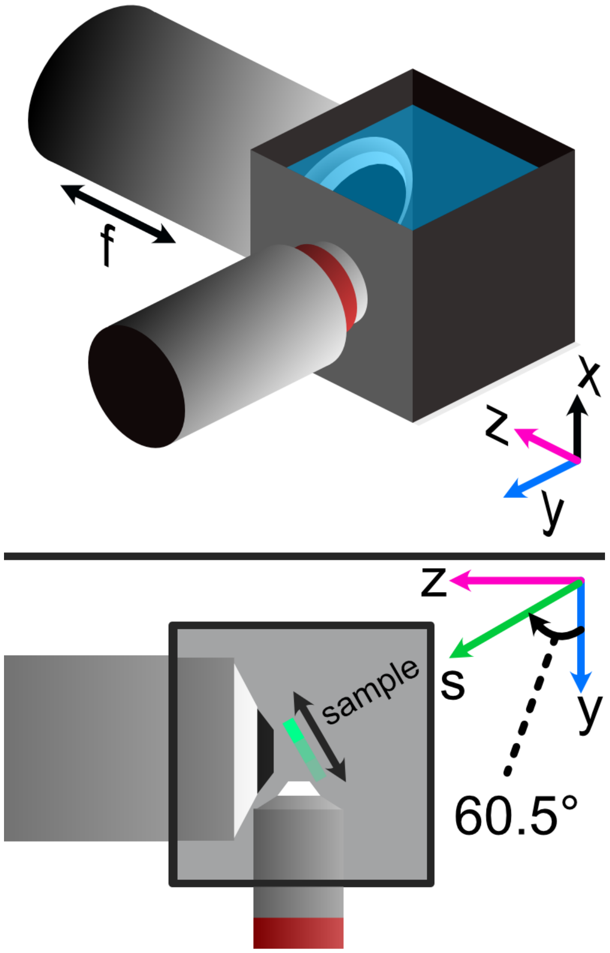

Fig. 2: **Figure 2** Layout of how the axis of the system are mapped

#### 8.3 Minimizing Spherical Aberrations

Once the system has been assembled to the point of being able to take image stacks, the process of minimizing the effects of spherical aberrations can begin. Spherical aberrations are typically introduced into optical systems due to the surface curvature of different lens elements. This type of aberration typically presents itself visually as a sort of stretching or bending of the focus of light in the system. Certain microscope objectives, such as the Nikon 25x/1.1 NA that we employ in this setup, have a built-in collar that can be adjusted to minimize spherical aberration.

In our system, we expect the effects of spherical aberrations to be along the axis of our detection path (defined as z in our imaging scheme). In order to visualize these effects and adjust the correction collar of our objective to mitigate them, we employ a process of taking a z-stack of fluorescent beads suspended in agarose and using ImageJ to quickly process those images.

1. Take a z-stack within navigate of your sample
2. Open up the z-stack within ImageJ
3. Reslice the z-stack (Image -> Stacks -> Reslice)
4. Do a maximum intensity project of the resliced stack (Image -> Stacks -> Z-Projection)
5. Take note if spherical aberration is present in the projected image.
6. If spherical aberration is still present, make slight adjustments to the objective correction collar and repeat Steps 1-5. Each time you adjust the correction collar, you will likely need to update the focus of the microscope.

As a note, observing the camera live-feed via navigate's "Continuous Scan" mode while adjusting the correction collar can help to get in the general vicinity of the correct placement of the correction collar. An example of how change in the correction collar affect live images are shown below for fluorescent beads. Aiming to get to get the beads near the expected light sheet position to be as in-focus as possible is a general guide for what direction to move the collar; however, true correction needs to be done with the z-projection method mentioned above.

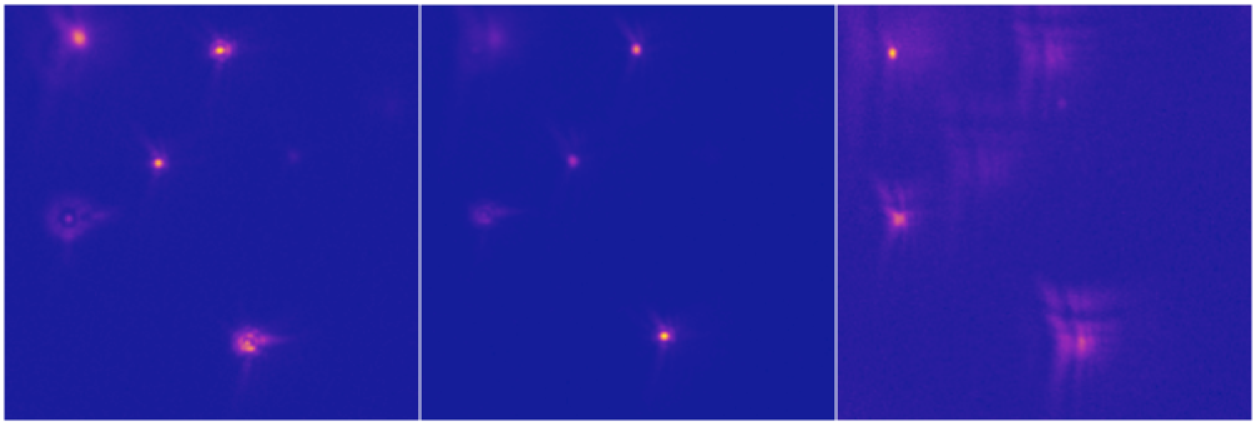

Fig. 3: **Figure 3:** Effects of adjusting the correction collar when imaging fluorescent beads

As a quick example of what an image of a z-projection could look like before and after trying to correct for spherical aberration is shown below. Here, one can see in the top panel that the bead features are essentially smoothed out and fuzzy due to aberrations, while in the bottom panel with adjustments made to the correction collar the beads appear much cleaner and focused.

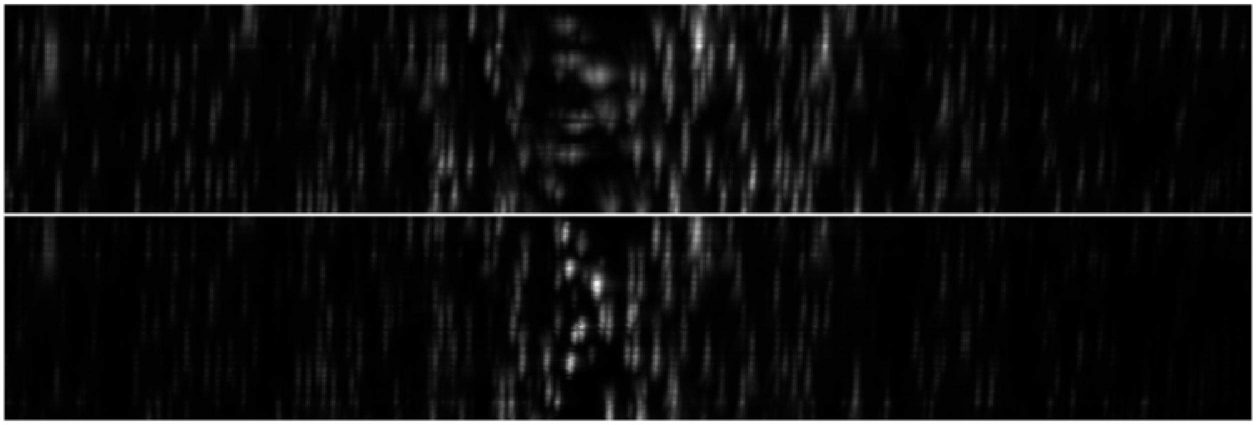

Fig. 4: **Figure 4:** Before and after of adjusting in Z-projections after adjusting the correction collar

#### 8.4 Sample Image Examples

When imaging samples, it is important to note the way in which the individual images comprising our z-stack might look as it might differ from what one might traditionally be accustomed to. In our imaging scheme, we essentially have a thin column of light spanning the vertical direction on our image. This results in each image in the z-stack being a thin snapshot of the whole sample, which can then be viewed through max intensity projections in ImageJ. To give a visual reference of what these individual images in a stack can look like for a biological sample, we provide the following images taken of a mouse embryonic fibroblast sample at a single z-position in a stack for 4 different imaging channels (gold = actin, gray = tubulin, cyan = nuclei, and magenta = Golgi apparatus).

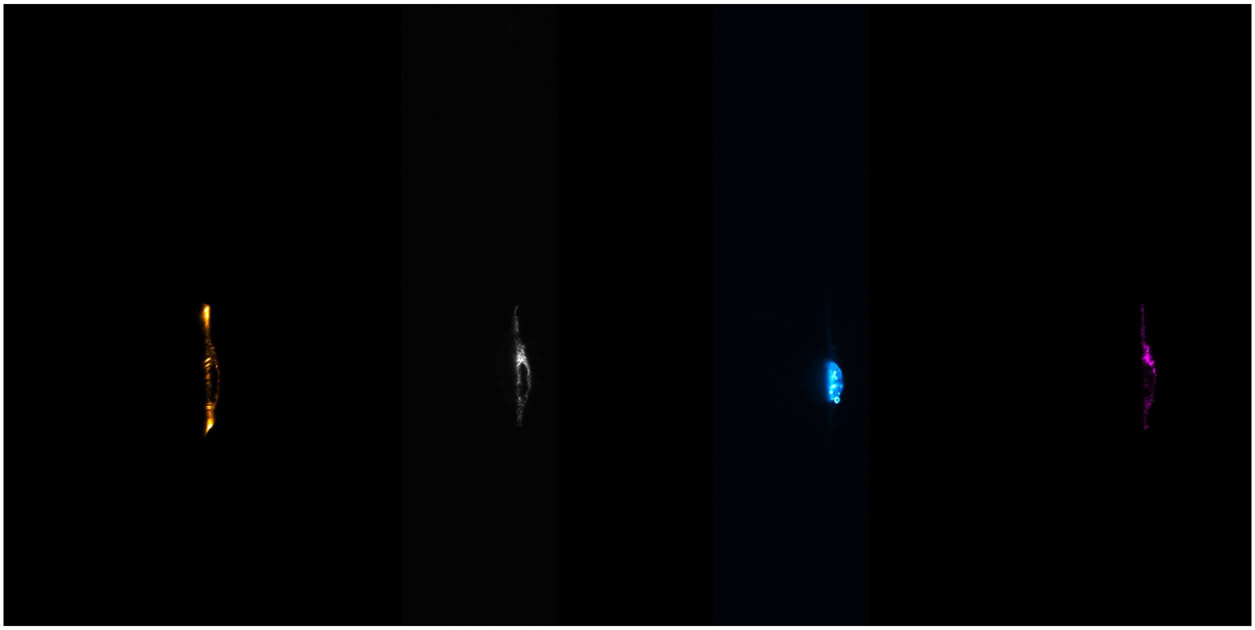

Fig. 5: **Figure 5:** Example individual images for MEF cells

##### 8.4.1 Image Stack Processing

#### 8.5 Deskewing

With an image stack acquired, some post processing is still required in order to remove the effects of shearing in our images. The root of this shearing is due to the angled method in which our sample is mounted and similarly, the angled path that the sample moves as the piezo is scanned. A basic visual idea of how deskewing affects the resulting image is shown below for 100 nm fluorescent beads. Here before deskewing for the same image plane (yz), the beads appear to be stacked in a straight line but oriented along an angle, which is not the most accurate representation of our system. On the deskewed image on the right, one can see that the beads are now properly angled correspond to our piezo angle mount, and that the PSFs of the beads is now correctly aligned along the z axis.

To do this deskew processing, we utilize custom-built python code via Jupyter notebooks available here. The user needs to provide the correct file path to the .tif image stack collected via navigate, as well as the parameters of the imaging system like z-step size, xy pixel size, and the angle that the images should be deskewed over. In our case, our deskew angle is equivalent to 90-60.5 degrees, where 60.5 degrees corresponds to the difference between the normal of our angle mount and the y-axis. If this value is unknown, one can use different values for the deskew angle until the bead PSFs are correctly aligned along the z-axis and not angled.

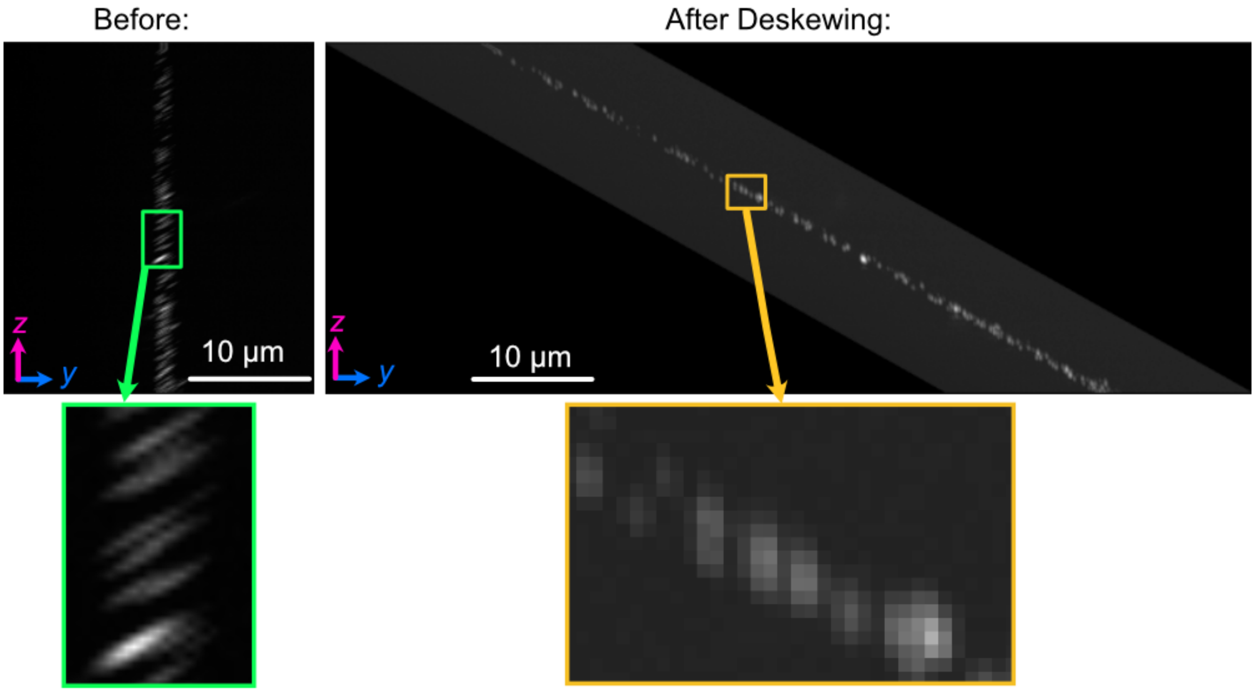

Fig. 6: **Figure 6:** Difference between an image set of 100 nm bead before deskewing (left) and after (right)

#### 8.6 Reslicing

Reslicing in ImageJ is a process that allows one to be able to reconstruct different planes of a 3D image set. In other words, it allows one to view the XY, XZ, and YZ projections of the same image set. In our system, our default viewing plane is the XY plane, and so we reslice to observe the XZ and YZ planes. The reslicing process within ImageJ is done after deskewing, and involves opening up the Reslicing panel (Image-> Stacks-> Reslice). Within this panel, one just needs to select the direction of the reslice (typically just top or left). For our system, top slicing provides us with the YZ plane view where one can observe the angled orientation of our sample setup after projection (Image-> Stacks-> Z Project). This is shown below for the same 100 nm bead samples used in the Deskewing and Rescaling portions of this page.

The same process can then be done to obtain the XZ plane view of our sample by reslicing left instead:

#### 8.7 Deconvolution

Deconvolution is an iterative post-processing technique that aims to enhance the resolution of a given image. Typically, in order to properly utilize deconvolution techniques one needs not only to have an image that they want to enhance, but also have an image of the corresponding point-spread-function (PSF) of the system used to take the image. We generate this PSF through taking an image stack of an isolated 100 nm fluorescent bead. For deconvolution we utilize PetaKit5D, which is open-source and MATLAB-based image processing code base. It should be noted that deconvolution techniques, while powerful, are also highly dependent on a variety of sensitive input parameters, and finding an effective combination of these parameters can often be a difficult process.

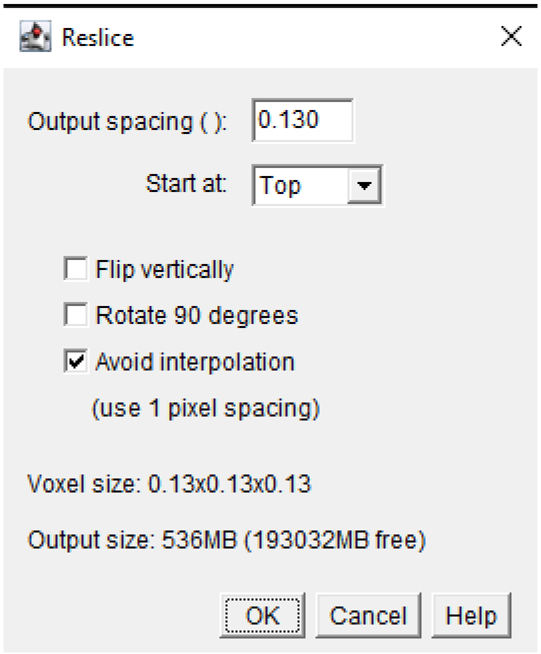

Fig. 7: **Figure 7:** Reslicing Panel for top reslicing

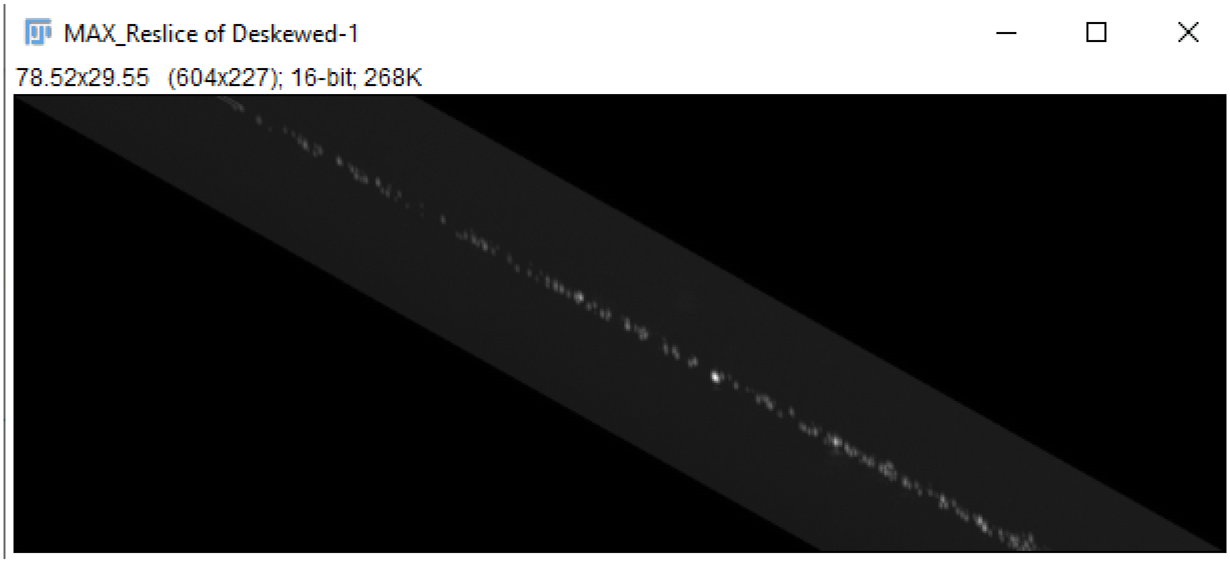

Fig. 8: **Figure 8:** The YZ projection of our bead images after reslicing.

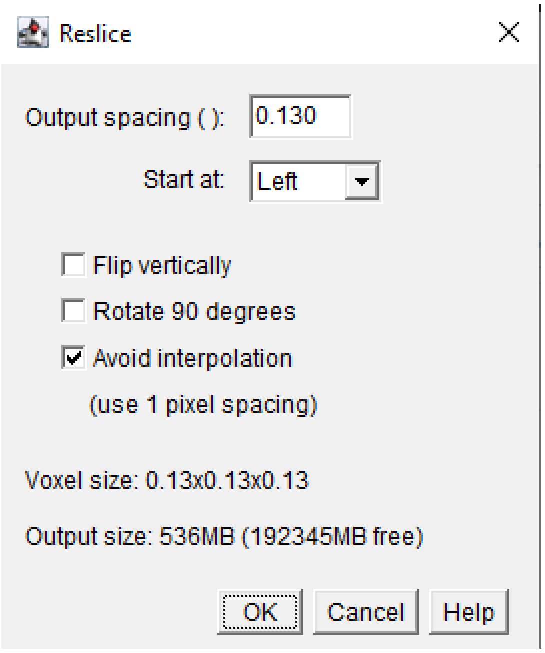

Fig. 9: **Figure 9:** Reslicing Panel for left reslicing

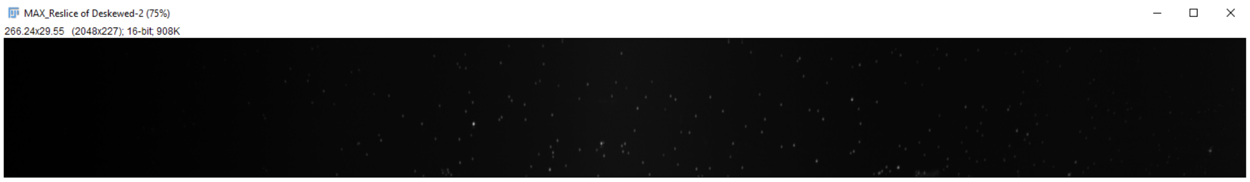

Fig. 10: **Figure 10** The XZ projection of our bead images after reslicing.

CHAPTER

NINE

### LIVE-CELL IMAGING

#### 9.1 Sample Chamber Design

In addition to fixed-cell imaging, there are also a variety of live-cell imaging applications where observing the evolution or behavior of a cell over time can be valuable. To be able to perform live-cell imaging, the most important aspect that needs to be addressed compared to fixed-cell imaging is that of a heat-regulated environment for the cells. In live-cell imaging, the typical temperature ranges that one would use are between 25-37C, with 90% of applications being done at 37C. We've traditionally done this temperature regulation in the past using a combined system comprised of thermocouples (for detection the ambient temperature), heating pads (to heat the cell sample chamber), and a heating controller (to manage signals to and from the thermocouples and heating pads and ensure the chamber remains at a specific temperature). It's also important to note that in addition to the sample chamber, gently heating the microscope objectives used is also advised to reduce potential effects of thermal drifting that effect imaging performance.

Our live-cell imaging chamber is meant to serve as an upgrade to our pre-existing sample chamber design and provide an all-in-one way to accurately heat the sample chamber as well as the two objectives used. The chamber is designed for the Thorlabs TL20X-MPL illumination objective and Nikon N25X-APO-MP detection objective pairing. As with our traditional chamber, there are two ports for these objectives with sets of o-ring grooves within them to form a water-tight seal around the objectives themselves.

In comparison to our traditional chamber design, there are a few changes made in this variant. The first is that there is not an additional third port for imaging of the light sheet itself. This allows us to place a heating pad along the two sides of the chamber with no ports and increase the overall surface area that is actively heated on the chamber. The other large change is that of additional large hoods around each of the objective ports. These hoods are designed for heating pads to be able to be wrapped around them to apply indirect air heating to the objectives.

#### 9.2 Parts List

Similarly to our philosophy when trying to source all of our components in our system from as few distinct vendors as possible, we sourced all of the heating elements for this system directly from Mcmaster Carr. There are ceratinly other options one can go with to accomplish similar heating capabilities, but we'll be basing our full heating assembly based on the following components:

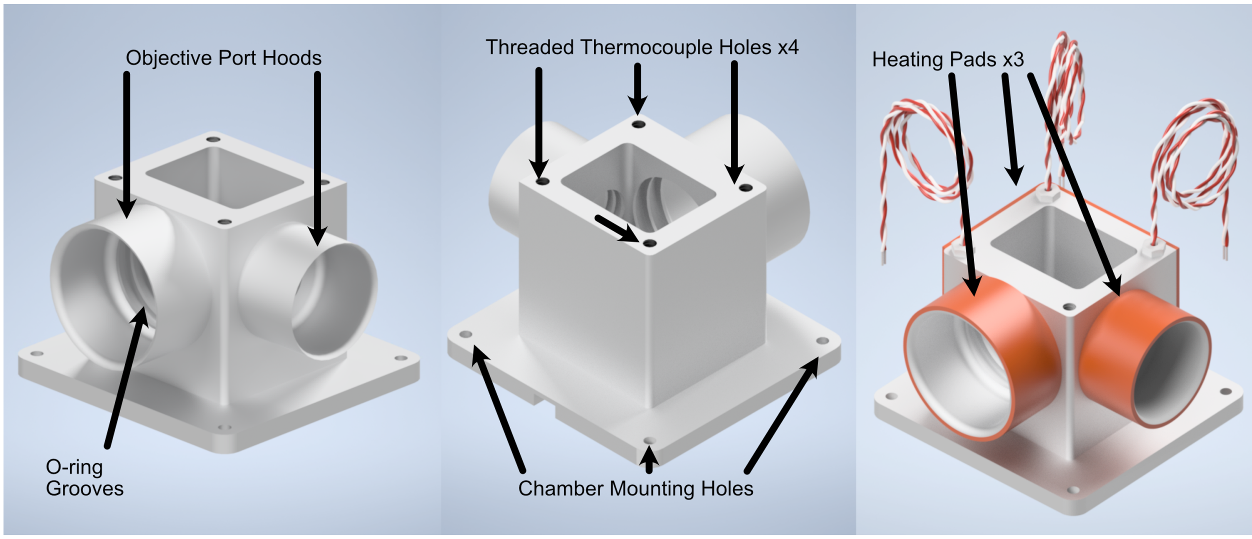

Fig. 1: **Figure 1** Our custom live-cell sample chamber

| Part | Vendor | Purpose |
|---|---|---|
| Live-Cell Sample Chamber | Xometry | Custom-designed live-cell sample chamber |
| Benchtop Autotuning Temperature Controller, Type J Thermocouple, 1 Bay | Mcmaster Carr (Tempco) | 1 Bay Temperature Controller for the larger heating pad on the sample chamber side |
| Benchtop Autotuning Temperature Controller, Type J Thermocouple, 2 Bay | Mcmaster Carr (Tempco) | 2 Bay Temperature Controller for the heating pads associated with the microscope objectives |
| 3" Dual Thermocouple Type J, 1/8" Diameter Probe | Mcmaster Carr (Reotemp) | Optional Dual-Probe Thermocouple for controlling objective temperatures |
| 1" Threaded Thermocouple 1/4"-20 Threading | Mcmaster Carr | Threaded Thermocouples to screw into sample chamber, 3 total if not using Dual-Probe thermocouple |
| 2x5" Adhesive Backed Heat Sheet | Mcmaster Carr (Benchmark Thermal) | Large heating pad to be placed on the outer surfaces of the sample chamber without objective ports |
| 5x1" Ultrathin Heat Sheet | Mcmaster Carr (All flex solutions) | Heating pad to wrap around illumination objective port |
| 6x1" Adhesive Backed Heat Sheet | Mcmaster Carr (Benchmark Thermal) | Heating pad to wrap around detection objective port |

### 9.3 Live-Cell Imaging Full Assembly

#### 9.3.1 Thermocouple Adapter Assembly

Thermocouples often come without the necessary adapter placed on the end of their wiring to connect them to a temperature controller. Below in Figure 2 we show the general process to attaching one of these adapters to the two thermocouple wires:

1. Unscrew the screws on the outer shell of the adapter.
2. Remove the outer shell element of the adapter.
3. Unscrew the inner screws over both terminals
4. Ensure that the inner screws are undone enough such that the metal plates underneath can be lifted.
5. Place the ends of the thermocouple wires beneath the metal plates of each terminal where red = - & white = +.
6. Screw the inner terminal screws tight.
7. Place the outer shell element back on the adapter and screw both screws into place on it.

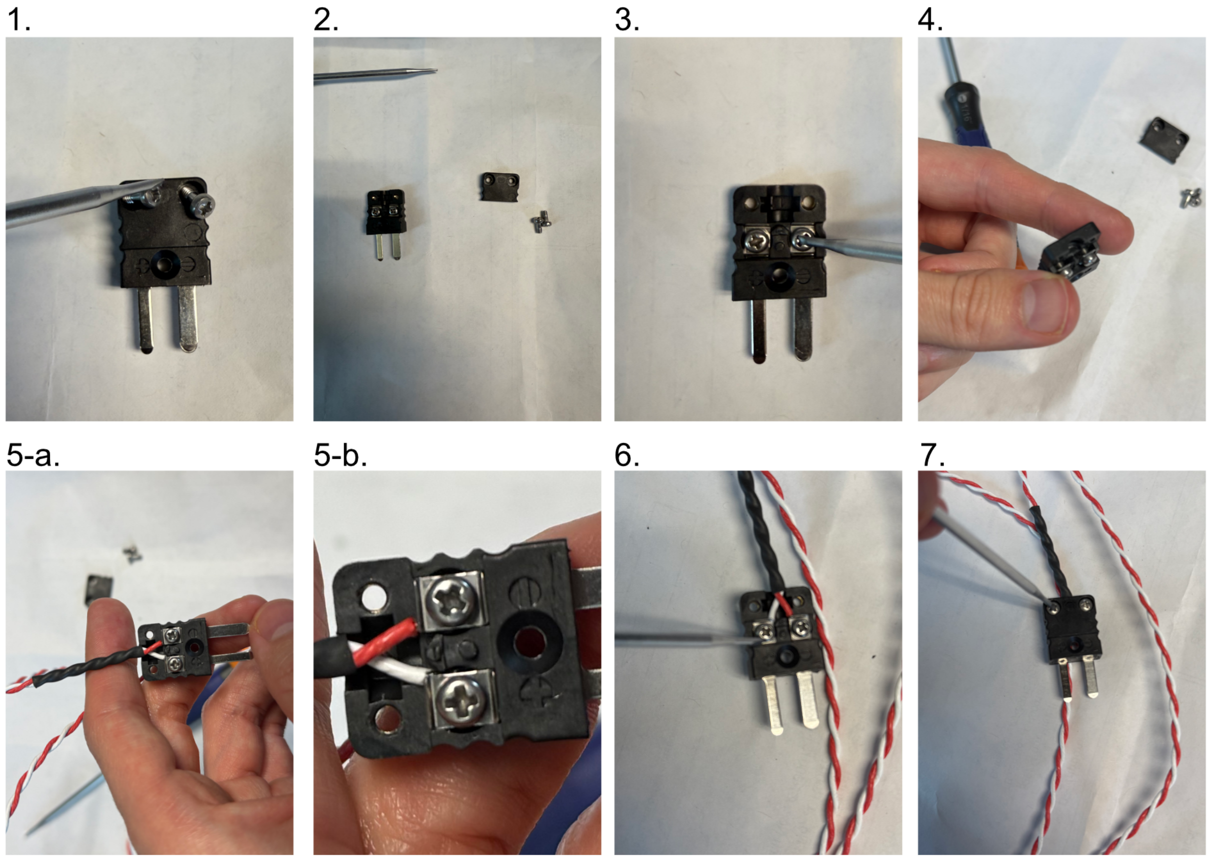

Fig. 2: **Figure 2:** Thermocouple adapter assembly process

##### 9.3.2 Heater Adapter Assembly

The heaters we utilize also lack the necessary adapter placed on the end of their wiring to connect them to a temperature controller. It also should be noted that for the heater elements, the two wires associated with them are unpolarized. Below in Figure 3 we show the general process to attaching one of these adapters to the two heater unit wires:

1. Unscrew the screws on the side of the adapter with the prongs sticking out
2. Remove the inner element of the adapter.
3. Unscrew the side screws over the gold and silver terminals on the inner element
4. Ensure that the plates within the terminals of the inner element are able to move
5. Thread the heater wires through the center hole of the detached front side of the adapter
6. Insert one heater wire into the silver terminal and one into the gold terminal and secure them by tightening the side screws.
7. Place the two adapter elements together, aligning the three screws on the inner element with the respective holes on the outer element, and tighten all the screws

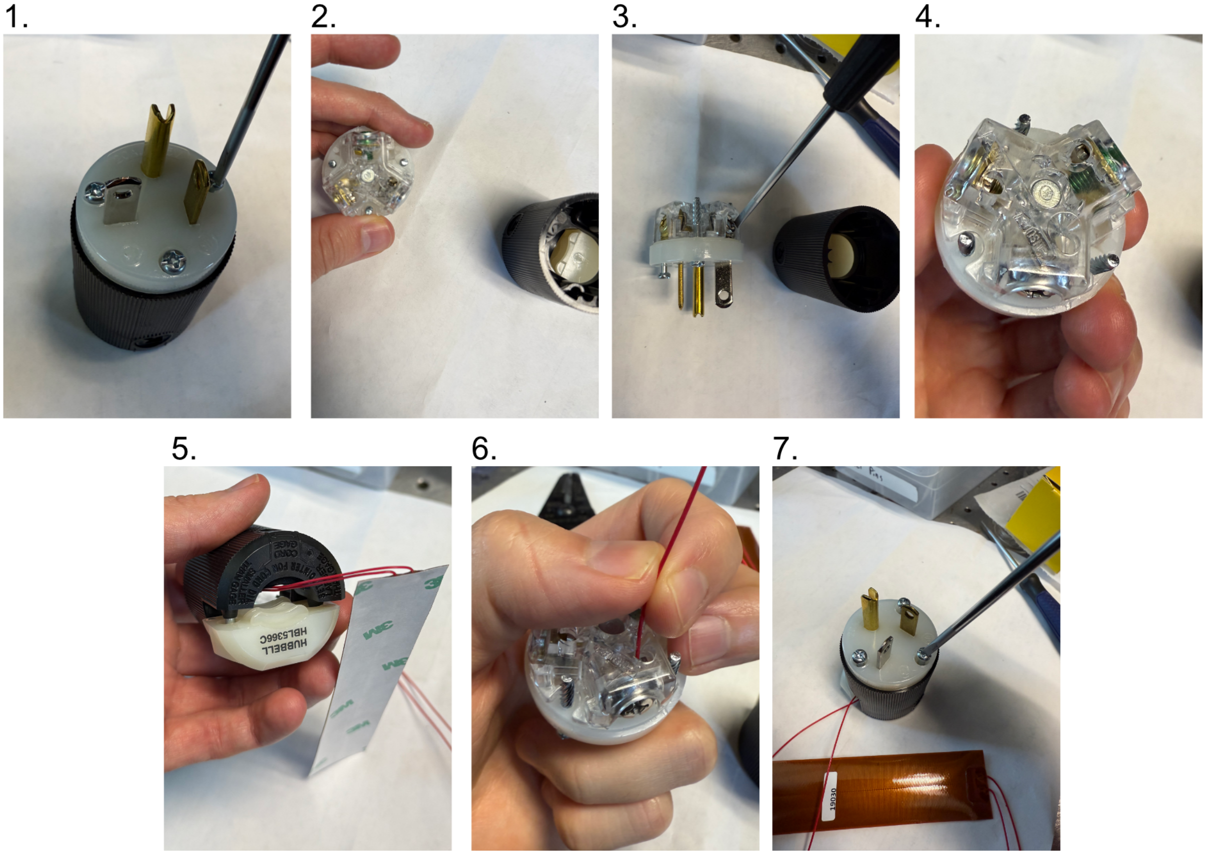

Fig. 3: **Figure 3:** Heater adapter assembly process

##### 9.3.3 Placing O-rings in the Sample Chamber

In order to ensure a watertight seal around our objectives, both of our objective ports feature two sets of o-rings surrounding their circumference. For our smaller port associated with the TL20X-MPL objective, we use oil-resistant Buna-N O-Rings with 11/16” inner diameter (ID) and 13/16” outer diameter (OD). For the larger port associated with the Nikon 20X objective, we used Buna o-rings with roughly 1.3” ID and 1.7” OD. These o-rings and their associated grooves are first coated with vacuum grease in the following process:

1. Unscrew vacuum grease container
2. Using either a finger or a cotton-tipped applicator, apply a layer of vacuum grease into and around the grooves on both ports
3. Put more vacuum grease on the cotton-tipped applicator
4. Using a finger or cotton-tipped applicator, take an o-ring and coat it fully in the vacuum grease.
5. Place the o-ring in the appropriate groove using a finger or tweezers to help ensure it sits within the groove
6. Repeat steps 3-5 for all 4 o-ring grooves in the chamber

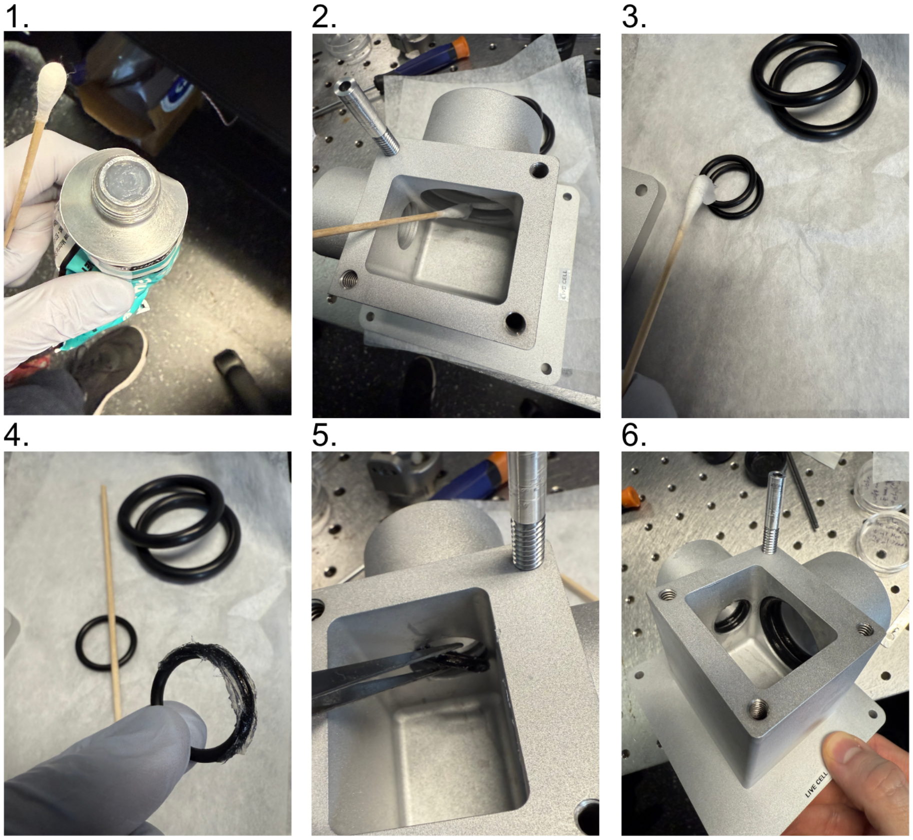

Fig. 4: **Figure 4:** Preparation and Placement of O-rings in the Sample Chamber

##### 9.3.4 Placement of the Heating Pads on the Sample Chamber

The placement of heating pads on the chamber is a straightforward process, particularly with the adhesive-backed pads we detailed in our parts list. In the application process, a general consideration should be made as to the direction the wires from the thermocouple are pointed in, as the actual wire lengths are relatively short you want to be as efficient as possible with them.The general assembly process is as follows:

1. Remove protective backing for the 5x1" ultrathin heat sheet
2. Wrap the 5x1" heat sheet carefully around the exterior of the smaller port on the sample chamber, oriented such that the wires are pointed towards the direction your heating controller will be.
3. Remove protective backing for the 2x5" heat sheet
4. Wrap the 2x5" heat sheet carefully around the exterior sides of the sample chamber with no ports on them, oriented such that the wires are pointed towards the direction of your heating controller.
5. Remove protective backing for the 6x1" ultrathin heat sheet
6. Wrap the 6x1" heat sheet carefully around the exterior of the larger port on the sample chamber, oriented such that the wires are pointed towards the direction your heating controller will be.

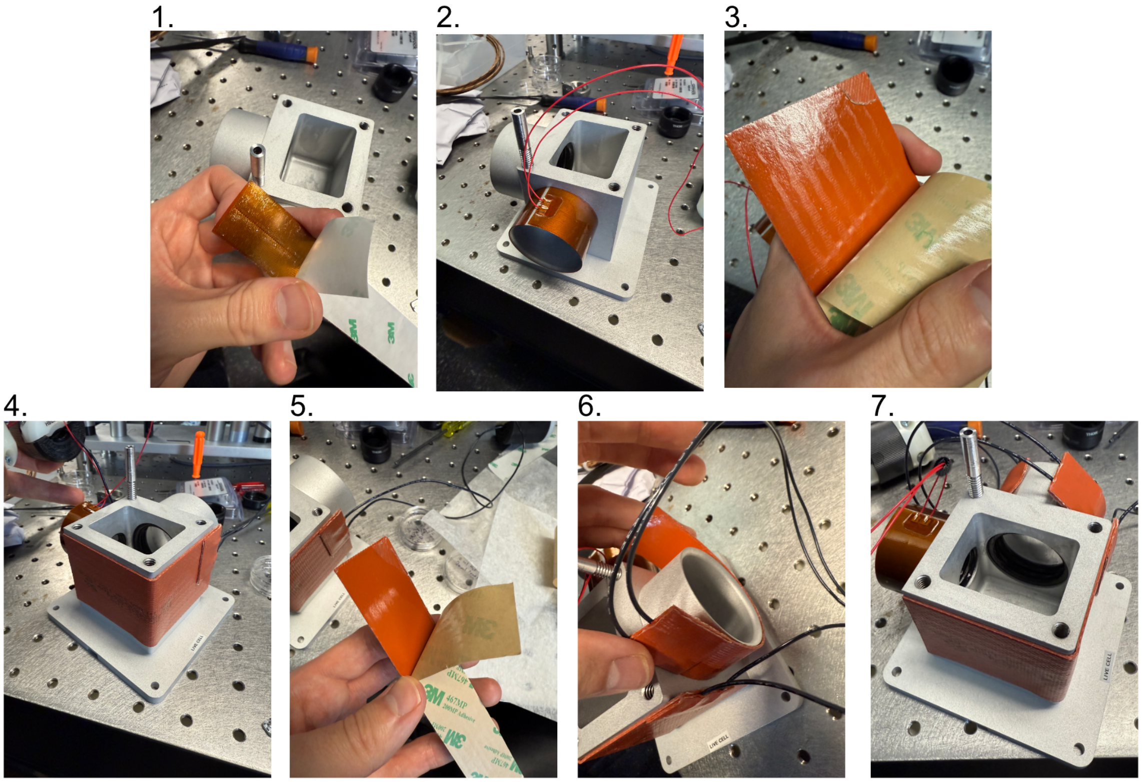

Fig. 5: **Figure 5:** Preparation and Placement of Heating Pads on the Sample Chamber

##### 9.3.5 Temperature Controller Assembly and Settings

With the individual components of our heating assembly prepared, the full assembly of the system is straightforward. There is some variation in terms of if one opts to use 3 threaded thermocouples or a combination of one dual-probe thermocouple and one threaded thermocouple, but in principle both configurations are valid and will yield the same system properties. In our case, we began by testing the dual-probe + 1 threaded thermocouple system which is reflected in our figure below. Our general process is as follows:

1. Mount the sample chamber and position it such that both objectives are able to enter the chamber (more detailed setup on aligning the objectives and positioning them can be found on our Microscope Assembly page).
2. Place chosen thermocouples into position in one of the 4 threaded thermocouple holes on the top of the sample chamber. We chose the dual-probe thermocouple to simultaneously control the heating of both objectives, so we placed it in the hole between the two objective ports. For our threaded thermocouple we chose it to correspond to our larger heating pad on the exterior of the chamber walls, and placed it in the corner that was closest to where we were able to physically place our individual controller for it.
3. Plug the thermocouple adapters for each thermocouple into their respective ports on their temperature controllers. In our case we had our two-bay controller for our two objective ports, and the one-bay controller for the larger exterior heating pad.
4. Plug the heating pad modules into the outlets on the back of the controllers corresponding to their respective ports
5. Plug the temperature controllers into an outlet and turn on the power
6. Using the two middle arrow buttons on the temperature controller display, adjust the temperature to the desired value. The controller will take temperature values from the thermocouples and automatically heat the heating pads accordingly to match the set value.

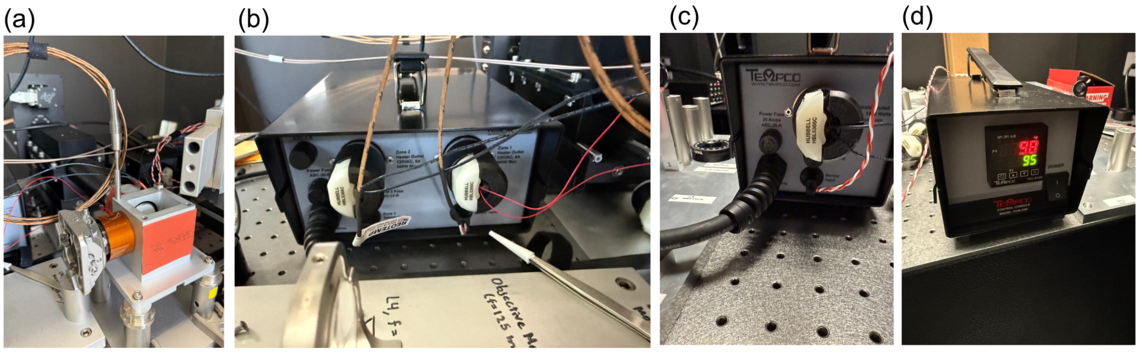

Fig. 6: **Figure 6:** Final Assembly and Setting of the Temperature Controllers

CHAPTER

TEN

### FUTURE RELEASES

The development of **Altair** is an ongoing effort, with future iterations aimed at increasing both **complexity and capability** to support volumetric imaging across a diverse range of specimens and experimental contexts. Each implementation will be **versioned** and accompanied by comprehensive documentation, ensuring that researchers can **replicate and modify** the system at a fraction of the cost of commercial alternatives. To foster **open development and community-driven innovation**, we encourage users to **request features and provide feedback** publicly via Altair GitHub Issues.

The projected design roadmap includes:

- **Sample-scanning light-sheet microscope** for imaging cells on glass coverslips.
- **Sample-scanning Axially Swept Light-Sheet Microscope (ASLM).**
- **Laser-scanning Axially Swept Light-Sheet Microscope.**
- **Sample-scanning Oblique Plane Microscope (OPM).**
- **Laser-scanning Oblique Plane Microscope.**

Beyond these core designs, we aim to develop **modular additions** that extend Altair's functionality, including **optogenetic stimulation** for targeted cellular activation and **multiplexed imaging** for simultaneous readouts of multiple probes. Through this iterative approach, Altair will continue evolving as a **highly adaptable, cost-effective, and open-source platform** for cutting-edge volumetric imaging.
